# Revealing the Neurobiology Underlying Interpersonal Neural Synchronization with Multimodal Data Fusion

**DOI:** 10.1101/2022.07.26.501562

**Authors:** Leon D. Lotter, Simon H. Kohl, Christian Gerloff, Laura Bell, Alexandra Niephaus, Jana A. Kruppa, Juergen Dukart, Martin Schulte-Rüther, Vanessa Reindl, Kerstin Konrad

## Abstract

Humans synchronize with one another to foster successful interactions. Here, we use a multimodal data fusion approach with the aim of elucidating the neurobiological mechanisms by which interpersonal neural synchronization (INS) occurs. Our meta-analysis of 22 functional magnetic resonance imaging and 69 near-infrared spectroscopy hyperscanning experiments (740 and 3,721 subjects) revealed robust brain-regional correlates of INS in the right temporoparietal junction and left ventral prefrontal cortex. Integrating this meta-analytic information with public databases, biobehavioral and brain-functional association analyses suggested that INS involves sensory-integrative hubs with functional connections to mentalizing and attention networks. On the molecular and genetic levels, we found INS to be associated with GABAergic neurotransmission and layer IV/V neuronal circuits, protracted developmental gene expression patterns, and disorders of neurodevelopment. Although limited by the indirect nature of phenotypic-molecular association analyses, our findings generate new testable hypotheses on the neurobiological basis of INS.

## 1. Introduction

Synchronization with the environment is a key mechanism that facilitates adaptation to varying environmental conditions in most living organisms, potentially providing an evolutionary advantage (Xue et al., 2019). In humans, adaptation to other people is a central survival mechanism and has been linked to *interpersonal synchronization* that occurs at multiple biobehavioral levels during human social interaction (Gordon et al., 2021; Harel et al., 2011; Mogan et al., 2017; V. Müller et al., 2013; Reindl et al., 2022). Interpersonal synchronization involves coordination of behavioral, physiological, or hormonal activities between people and may represent adaptive capacities that allow humans to access another’s internal arousal state (Mizugaki et al., 2015), share and regulate emotions, increase social affiliation, empathy, and prosocial commitment (Mogan et al., 2017), facilitate learning (Pan et al., 2021), and adapt to collective behaviors and group norms (Wiltermuth & Heath, 2009; Reinero et al., 2021). Revealing the neurobiology underlying interpersonal synchronization will improve our understanding of the fundamental mechanisms by which humans adapt to and engage with others. Knowledge on how these mechanisms work and how they fail has broad implications for educational sciences and developmental and social neurosciences on the one hand, and for intra- and intergroup conflict studies and psychiatric health care on the other.

Interpersonal synchronization has been extended to the *neural activity* of interacting individuals, often referred to as *interpersonal neural synchronization* (INS) (Jiang et al., 2015), interbrain synchrony, or brain-to-brain synchrony (Mu et al., 2016; Dikker et al., 2014). In a single brain, rhythmic oscillations of neurons may lead to neuronal signal coherence through synchronization of excitatory states, thereby enabling neuronal information transfer and interaction (Fries, 2005). Local neuronal oscillations have been linked to *excitation-inhibition (E/I) balance*, regulated by GABAergic and glutamatergic neuron populations (Gonzalez-Burgos & Lewis, 2008; Sears & Hewett, 2021), which may also be a driving factor for long-range synchronization (Stagg et al., 2014). Electrophysiologically, within-brain synchronization may be driven by excitatory cortico-cortical connections (Uhlhaas & Singer, 2006), together with subcortical structures, in particular the thalamus (Llinás & Steriade, 2006). Across brains, in analogy to the oscillations of individual neurons, our brains and their sensory systems may also rhythmically sample information from the environment. Information transfer is then not enabled via direct physical contact but indirectly through actions arising from an individual’s motor system (e.g., speech, sounds, gestures, or eye contact). These actions are transmitted through the environment and sampled by an interaction partner’s sensory system. In each individual in a dyad or group, rhythmical neuronal oscillations may then synchronize (Hasson & Frith, 2016).

By simultaneous brain recordings from two or more subjects, termed *hyperscanning* (Montague et al., 2002), it is now possible to quantify the temporal and spatial similarities of brain signals while the individuals engage in interpersonal interaction. Alternatively, one subject can be scanned after another in response to prerecorded stimuli of the first person, often termed *pseudohyperscanning* (Babiloni & Astolfi, 2014; Schoot et al., 2016). Methodologically, human hyperscanning experiments have been performed over the full spectrum of noninvasive electrical and hemodynamic brain imaging techniques, with electroencephalography (EEG), functional near-infrared spectroscopy (fNIRS), and functional magnetic resonance imaging (fMRI) being the most widely used (Babiloni & Astolfi, 2014; Czeszumski et al., 2020).

Previous human hyperscanning studies identified a variety of brain regions that contribute to INS, including the medial prefrontal cortex (PFC), anterior cingulate (Babiloni et al., 2007; Yun et al., 2012), superior temporal gyrus (STG) and right temporoparietal junction (rTPJ) (Stolk et al., 2014; Bilek et al., 2015; Kinreich et al., 2017), and insular cortex (Koike, Tanabe, et al., 2019). The first meta-analytic evaluation of 13 fNIRS hyperscanning studies involving interpersonal cooperation confirmed INS in the PFC and TPJ (Czeszumski et al., 2022). The observed brain region patterns suggest connections to brain networks known to be associated with mentalization (Schurz et al., 2021; Bilek et al., 2015), social cognition and interaction (Feng et al., 2021), predictive coding (Ficco et al., 2021; Shamay-Tsoory et al., 2019), and mirroring (Rizzolatti & Craighero, 2004; Schippers et al., 2010). These patterns indicate that INS involves complex cognitive processes, including theory of mind (ToM), mental modeling, prediction, emulation, and simulation of behavioral and affective states.

Developmentally, INS might be rooted early in human life, with synchronous caregiver-infant interactions being critical for establishing affiliative bonds (Feldman, 2017) and impacting long-term developmental outcomes (Atzil & Gendron, 2017). In the brain, on both cognitive and functional levels, INS has been embedded in a predictive coding framework (*social alignment system*), mediated by a three-component feedback loop consisting of an *observation-execution/alignment*, an *error-monitoring*, and a *reward system* thought to be activated by and to reinforce successful alignment (Shamay-Tsoory et al., 2019). As postulated in the *mutual prediction theory*, coherent patterns of brain activity in two interacting partners might result from the sum of neural activities from co-localized neurons (i) encoding self-behavior as well as (ii) encoding predictions of the partner’s behavior (Hamilton, 2021; Kingsbury et al., 2019). On the neurophysiological level, the connectivity between two brains or among multiple brains may be shaped by social contact in analogy to the Hebbian rule for synaptic connectivity (“fire together, wire together”) (Shamay-Tsoory, 2021). Here, the cortical activity of one subject engaged in a certain behavior would translate into the cortical activity of an interacting subject, with the repetition of this social interaction reshaping interbrain functional connectivity not only in dyads but potentially in entire social groups (Ramakrishnan et al., 2015). On the neurochemical level, oxytocin and dopamine have been discussed as the key neurotransmitter systems involved (Feldman, 2017; Gvirts & Perlmutter, 2020; Mu et al., 2016) given their pivotal roles in social functions (MacDonald & MacDonald, 2010), reward processing (Glimcher, 2011), and reciprocal interactions between the two systems in the mesolimbic tract (Baskerville & Douglas, 2010). Related to the *social alignment system* (Shamay-Tsoory et al., 2019), these mesolimbic neurotransmitter systems may regulate a *mutual social attention system* located in the PFC and TPJ, possibly enabling selective attention in social interactions through reward-related feedback mechanisms (Gvirts & Perlmutter, 2020).

The rapidly evolving hyperscanning research field and our inherent fascination with human social abilities and inabilities led to a steadily growing number of theoretical accounts attempting to explain the phenomenon of INS. However, robust evidence to ground these theories on is still lacking and many of the proposed frameworks have yet to be tested empirically. In particular, attempts to develop models moving beyond brain region correlates have been limited by the unavailability of empirical data. Given that social cognition, oxytocin signaling, and E/I balance are considered to be connected on neurophysiological levels (Lopatina et al., 2018), the extent to which these mechanisms underlying within-brain synchronization are involved in INS also remains to be explored.

The current study aimed to identify a common neural substrate and formulate new testable hypotheses regarding the neurophysiological mechanisms of INS. To achieve this goal, we used multimodal data fusion techniques as powerful tools to integrate data from imaging, genetic, and behavioral levels. Through integrative meta-analytic techniques, data fusion approaches, and null model-based hypothesis tests (Figure 1), we confirm robust spatial convergence of INS in the rTPJ as well as an involvement of the ventral PFC, and provide first evidence for an important and previously unacknowledged role of GABAergic neurotransmission and E/I balance in human INS.

**Figure 1:**
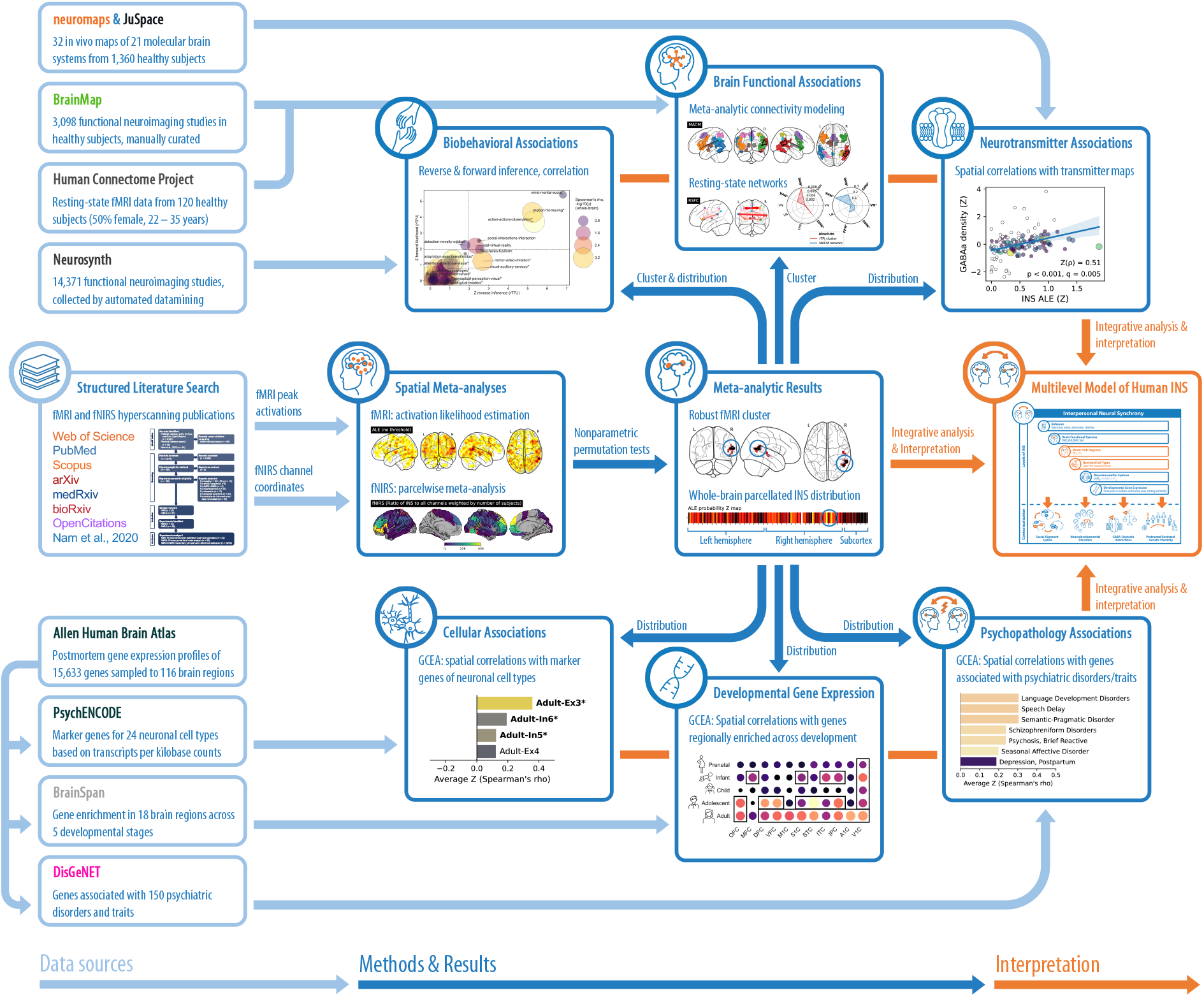
The multimodal data fusion approach to explore the neurobiology of human INS. The figure outlines the multimodal data fusion workflow applied in the present study. Depicted are data sources and major analysis steps applied to generate multilevel knowledge and new hypotheses about the neurobiological basis of INS. Abbreviations: INS = interpersonal neural synchrony, fMRI = functional magnetic resonance imaging, fNIRS = functional near-infrared spectroscopy, GCEA = gene-category enrichment analysis.

## 2. Materials and Methods

First, to identify a common brain regional correlate of INS, we collected the currently available fMRI and fNIRS hyperscanning data through an inclusive literature search and submitted it to spatial meta-analyses. Second, based on these brain correlates of INS, we assessed INS-related functional brain networks and biobehavioral association patterns on both brain regional and whole brain levels. Third, extending our results to a molecular level, we explored how the whole-brain INS distribution aligned with neurotransmitter systems and how spatially related gene expression patterns connected INS to specific neuronal cell types, brain development, and psychopathology (Figure 1).

### 2.1. Software, code, and data availability

The analyses were conducted in *Python* (3.8.8) and *Matlab* (R2021a) environments. The following software and packages were used: Literature search: *SetYouFree* (0.0.1) (Gerloff, Lotter, et al., 2022), *Cadima* (Kohl et al., 2018). Neuroimaging meta-analysis and image manipulation: *NiMARE* (0.0.12rc7) (Salo et al., 2018), *Nilearn* (0.9.1) (Abraham et al., 2014), *AtlasReader* (0.1.2) (Notter et al., 2019). FNIRS probe reconstruction: *AtlasViewer* (2.12.12) (Aasted et al., 2015). FMRI data processing: *CONN* (20b) (Whitfield-Gabrieli & Nieto-Castanon, 2012). Nuclear imaging/mRNA expression data retrieval and spatial correlation analyses: *JuSpace* (1.3) (Dukart et al., 2021), *neuromaps* (0.0.2) (Markello et al., 2022), *JuSpyce* (0.0.1) (Lotter & Dukart, 2022), *brainSMASH* (0.11.0) (Burt et al., 2020)*, abagen* (0.1.3) (Markello et al., 2021), *ABAnnotate* (0.1.0) (Lotter et al., 2022). Visualizations: *Nilearn, Matplotlib* (3.4.3) (Hunter, 2007), *seaborn* (0.11.2) (Waskom, 2021), *surfplot* (0.1.0) (Gale et al., 2021; Vos de Wael et al., 2020), *GO-Figure!* (1.0.1) (Reijnders & Waterhouse, 2021)*, pyvis* (0.2.1), *WordCloud* (1.8.1). Furthermore: *scipy* (1.8.1) (Virtanen et al., 2020), *statsmodels* (0.13.1), *numpy* (1.22.3),*pandas* (1.4.2).

We provide all code and data necessary to reproduce our results in a GitHub repository (https://github.com/LeonDLotter/MAsync). Raw Human Connectome Project neuroimaging data (Van Essen et al., 2013) are openly accessible otherwise (https://db.humanconnectome.org). All code can be found in an annotated Jupyter notebook, available in the repository and in HTML format (https://leondlotter.github.io/MAsync/MAsync_analyses.html).

### 2.2. Ethics

All analyses conducted and reported here rely on third-party data that were acquired in accordance with the respective institute’s ethical guidelines. The ethics committee of the RWTH Aachen University, Germany approved the use of these data (EK 188/22).

### 2.3. Literature search and data extraction

Currently published fMRI and fNIRS hyperscanning experiments were identified in a two-step semi-automated literature search (Gerloff, Lotter, et al., 2022). Methodologically, we focused on fMRI and fNIRS, as both methods rely on the hemodynamic signal, provide a relatively high spatial resolution (as compared to EEG), and together form the currently largest body of hyperscanning literature (Nam et al., 2020; Czeszumski et al., 2020). In the following, we will use the terms *publication* to refer to original studies and *experiment* to refer to sets of data obtained from independent study subjects which can cover data from multiple publications (Tables S1 and S2).

#### 2.3.1. Literature search process

An initial semi-structured literature search relying on PubMed, Web of Science, and Scopus and a manual screening of Google Scholar citation lists were performed in spring 2021 and updated through PubMed alerts during the following months. The final search was conducted on December 12, 2021, through open-access APIs of PubMed, Scopus, arXiv, bioRxiv, and medRxiv using SetYouFree (https://github.com/ChristianGerloff/set-you-free) followed by forward- and backward-citation searches of the resulting records in the OpenCitations database (Peroni & Shotton, 2020) (excluding records without abstracts). The results of both literature searches, together with references from a related review (Nam et al., 2020), were imported into Cadima (https://www.cadima.info/) for manual screening, eligibility assessment, and a final inclusion decision following PRISMA 2020 guidelines (Page et al., 2021). First, titles and abstracts were screened by one of five independent reviewers (LDL, LB, AN, JK, and CG) followed by a full text assessment by two of four reviewers unaware of each other’s inclusion/exclusion decisions (LDL, LB, AN, and JK).

#### 2.3.2. Study inclusion and data extraction

We searched for (i) fMRI or fNIRS hyperscanning or pseudohyperscanning publications assessing (ii) temporal synchrony at (iii) whole-brain (fMRI) or channelwise (fNIRS) levels between (iv) hemodynamic brain signals of (v) healthy adults (18–65 years) engaging in (vi) uni- or bidirectional interactions (Supplement 1.1.1–3). While pseudohyperscanning, in which typically one subject is scanned after another in response to pre-recorded stimuli of the first person (Babiloni et al., 2007; Schoot et al., 2016), allow for a more precise control of the experimental stimuli, it may not fully capture potential neurobiological representations unique to real-life reciprocal social interactions. However, we included both hyperscanning and pseudohyperscanning studies, as the latter may still shine light on certain aspects of social interaction in the sense that the unidirectional communicative aspect of, e.g., a subject listening to a speaker in a two-person communicative setting can be seen as a subaspect of an actual bidirectional social interaction.

Brain coordinates or group-level imaging data depicting INS-related foci were extracted from the included experiments (Supplement 1.1.4–6), requested from the authors, or, in case of multiple fNIRS studies, derived by reconstructing reported probe setups (see below). We were interested in analyses contrasting INS during interpersonal interaction with rest, control, or randomization conditions, independent of the type of interaction, as we aimed to identify the common neural substrate of INS (e.g., if a study contrasted INS during *cooperation, competition*, and *control* conditions, we included the combined result as *cooperation/competition > control*). We additionally included studies that reported only more specific contrasts (e.g., *INS after feedback > INS prior to feedback)* and evaluated, in post-hoc assessments, how the inclusion of these studies influenced the meta-analytic results (only relevant for fNIRS studies). Methodologically, we included studies independent of the connectivity estimator (i.e., timeseries correlation or prediction, wavelet coherence) if the method captured temporal synchrony. If studies investigated the effect of temporally shifting subject timeseries, we aimed to include only results reflecting zero-lag relationships to increase homogeneity. Therefore, the broad concept of “significant INS” studied here can be summarized as *similarity in the temporal variation of brain-derived blood oxygenation-dependent signals measured in humans during interpersonal interaction relative to non-interaction conditions*.

### 2.4. Spatial meta-analysis of fMRI experiments

We used activation likelihood estimation (ALE) to identify consistent spatial correlates of INS. Briefly, ALE provides brain-wide convergence maps combining the experiment-level activation maps modelled from the reported INS foci by convolving each focus with a sample-size dependent Gaussian kernel (Eickhoff et al., 2012; Turkeltaub et al., 2002, 2012). A nonparametric permutation procedure then distinguishes true convergence of INS foci from random spatial patterns (5,000 permutations) (Eickhoff et al., 2012, 2016).

#### 2.4.1. ALE

All coordinate based meta-analyses were performed using the Neuroimaging Meta-Analysis Research Environment (NiMARE; https://github.com/neurostuff/NiMARE). All contrasts and coordinates derived from the same sample (experiment) were concatenated. For each experiment, an activation map in 2-mm isotropic Montreal Neurological Institute (MNI)-152 space (Fonov et al., 2011) was estimated by convolving each focus with a Gaussian kernel. The width of the kernel, at half of the maximum of the height of the Gaussian, was determined based on the sample sizes of each experiment (Eickhoff et al., 2012; Turkeltaub et al., 2002, 2012). If foci from the same experiment overlapped, only the maximum voxelwise values were retained (Eickhoff et al., 2012). The union of these experiment-level data constituted the meta-analytic convergence map. Voxelwise statistical significance was determined based on an empirically derived null distribution (Eickhoff et al., 2012), a primary threshold of *p* < .001 was used to form clusters (extended by a threshold of *p* < .01 to increase sensitivity for weak effects) (Eklund et al., 2016), and a null distribution of cluster masses (5,000 iterations) was generated by randomly drawing coordinates from a gray matter template. By comparison of the actual cluster masses to the null distribution of cluster masses, each cluster was assigned a familywise error (FWE)-corrected *p* value and significant clusters were retained by thresholding the cluster map at –*log*_10_(*p*) > ~1.3. We relied on comparisons of cluster masses (the sum of voxel values) to estimate significance, as this has previously been shown to be more powerful than cluster inference based on size (the number of voxels) (H. Zhang et al., 2009). All subsequent analyses relied either on cluster-level FWE-corrected and binarized ALE clusters (depicting brain regions of significant spatial convergence of INS) or on the unthresholded Z-maps generated from ALE-derived voxel-level *p* values (reflecting the continuous probability of observing INS for every voxel).

#### 2.4.2. Influence of individual experiments and risk of publication bias

To estimate experimentwise influences on the overall ALE result (Eickhoff et al., 2016), we iteratively calculated the contribution of experiment x as 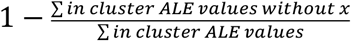. Noting that this approach is not exact due to the nonlinear ALE union computation, from a practical perspective, it is sufficient to approximate contributions and identify the exaggerated influence of individual experiments (Eickhoff et al., 2016). We then calculated the spatial conjunction of all resulting thresholded and binarized maps (Nichols et al., 2005) to demonstrate which clusters persisted in each iteration.

Clusterwise robustness against publication bias was estimated as the fail-safe-N (Acar et al., 2018). For each cluster, noise experiments in which the foci did not contribute to the cluster were generated. We then estimated the minimum number of noise experiments needed to render the cluster insignificant, reflecting the number of negative studies that could have “remained in the file drawer” (Supplement 1.2).

### 2.5. Spatial meta-analysis of fNIRS experiments

As, to date, the fNIRS field is still limited with respect to methodological standardization and availability of specific meta-analytic techniques, we developed a meta-analytic fNIRS evaluation in accordance with the ALE approach. In brief, for each of 100 cortical brain parcels (Schaefer et al., 2018), we collected information on whether or not INS was observed in fNIRS channels sampling the respective regions along with the overall number of subjects and experiments contributing to this information. We then calculated a parcelwise “fNIRS index” incorporating all available information for further evaluation and tested for parcelwise significance by randomizing channel-parcel-assignments (1,000 permutations).

#### 2.5.1. Coordinate extraction and reconstruction

Most fNIRS studies use probe arrays with standard formats positioned on the participant’s head according to coordinates within the international EEG positioning system. Commonly used methods to derive approximate locations on the brain surface are (i) registration using an anatomical MRI scan of one or more subjects, (ii) registration after digitization of channel positions using a 3D digitizer or (iii) *virtual registration* based on a digital model of the optode array and a reference database (Tsuzuki et al., 2007; Tsuzuki & Dan, 2014). The full workflow we followed to extract fNIRS coordinates is outlined in Supplement 1.1.6. When possible, we included coordinates as reported or sent to us by the authors. When necessary, we obtained coordinates from a database (Tsuzuki et al., 2007), or from other studies conducted by the same research groups, or we reconstructed the optode positions using AtlasViewer (Table S2). Experiments for which this workflow failed were excluded.

#### 2.5.2. FNIRS data analysis

For fNIRS data, standardized results reporting systems are currently being developed (Yücel et al., 2021), and no specific meta-analytic analysis techniques are available. To approximate a meta-analytic evaluation of fNIRS INS findings, we used a 100-parcel volumetric cortical atlas (Schaefer et al., 2018) to summarize fNIRS data by region. We assigned the nearest atlas parcel to each fNIRS channel using a kd-tree (Virtanen et al., 2020). Then, for each parcel, we collected the overall number of channels, the number of channels showing INS, and the corresponding numbers of experiments and subjects contributing to that information. To compare the results between parcels, we used three indices calculated as

i. *N_significant channels_*,
ii. 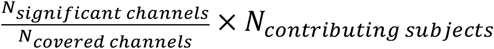, and
iii. 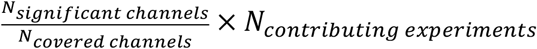.

We focused on the second index as it incorporated all available fNIRS data. To identify regions with the highest probability of the observed indices not being due to chance, we then permuted the channel-parcel assignment (1,000 iterations), estimated exact one-sided *p* values for each parcel and each “fNIRS index”, and applied FDR correction across parcels per index. We preferred this ALE-like approach over effect size-based meta-analyses for each parcel (Czeszumski et al., 2022), as the latter would have severely limited eligible studies due to their methodological heterogeneity. The results were visualized on fsaverage surface templates after surface transformation (Markello et al., 2022; Wu et al., 2018). By using parcellation-level instead of voxel-level data, we aimed to approximate the spatial resolution of fNIRS data, taking into account the added spatial uncertainty due to post-hoc reconstruction of channel coordinates without detailed information on head shape, size, and probe positioning.

Second, we conducted a joint meta-analysis of all neuroimaging data by evaluating fNIRS foci along with fMRI foci by means of an ALE. To adapt fNIRS data to the ALE method, we kept the ALE kernel size to a constant 10-mm FWHM for all fNIRS experiments as the kernel’s sample size-FWHM function was developed for fMRI data.

To further incorporate the spatial uncertainty of fNIRS data in our analyses, we iteratively (1,000 iterations) recalculated parcelwise and fNIRS-ALE meta-analyses after randomization of fNIRS coordinates (10 mm radius constrained to the cortical surface; Supplement 1.3).

### 2.6. Meta-analytic coactivation and resting-state functional connectivity

To establish the role of our meta-analytic findings in a whole-brain functional context, we constructed a co-activation network using meta-analytic connectivity modeling (MACM) (Eickhoff et al., 2011; Langner et al., 2014) on BrainMap data (Laird et al., 2011). As MACM does not provide information on interregional connection strength, we assessed resting-state functional connectivity (RSFC) patterns within the MACM network (Whitfield-Gabrieli & Nieto-Castanon, 2012).

#### 2.6.1. MACM

We performed MACM by performing an ALE on all BrainMap experiments (Laird et al., 2009, 2011) that had at least one activation focus within the robust rTPJ cluster (voxel-level *p* < .001). Only this cluster was used as the other INS-related ALE clusters proved unstable or, in the case of fNIRS analysis, did not survive multiple-comparison correction. Data were constrained to *activations* from *normal mapping* studies (i.e., those involving healthy control participants) and downloaded via Sleuth (3.0.4, https://brainmap.org/sleuth/). The resulting patterns resemble the network of task-related coactivation associated with the region of origin and are closely related to functional networks derived by RSFC analysis (Eickhoff et al., 2011). We relied on the BrainMap database as experiments and coordinates were manually screened by a dedicated team, promising greater precision and specificity of the resulting networks compared to automated data mining approaches.

To identify the most specific regions coactivated with the INS cluster, we used a specific coactivation likelihood estimation (Langner et al., 2014) in a separate analysis constructing a MACM network controlled for the baseline activation rate of all included BrainMap studies (3,098 experiments).

To validate the MACM results, and to assess whether the resulting activation patterns mirrored those of the original INS data, we computed the spatial correlation pattern between Z-maps derived from MACM and INS analyses after parcellation into 116 functionally defined whole-brain parcels [100 cortical (Schaefer et al., 2018) and 16 subcortical parcels (Tian et al., 2020)].

#### 2.6.2. RSFC

As MACM is not suited to quantify the connection strength between pairs of regions, we elucidated the functional connectivity patterns within the coactivation network using resting-state fMRI, thereby validating the presence of the network in single-subject data. For this, we relied on open access data from 120 Human Connectome Project subjects (50% female, 20 subjects per sex randomly drawn from each age group: 22–25, 26–30, and 31–35 years). The subject-level average timeseries of each MACM cluster was extracted and correlated using semipartial Pearson correlations. The *p* values resulting from connectionwise one-sample t tests were thresholded at *p* < .05 (Bonferroni-corrected). Only positive connections were retained to exclude potential artifacts introduced through noise regression (Murphy et al., 2009) (Supplement 1.4).

### 2.7. Functional contextualization

To explore the functional context of the observed activation patterns, we characterized relationships to established brain-wide resting-state networks (Yeo et al., 2011; Chen et al., 2018), determined associations between our INS-findings and biobehavioral domains in the Neurosynth database (Yarkoni et al., 2011) labeled with functional domain-related terms (Poldrack et al., 2012), and finally assessed relationships to meta-analytic networks underlying INS-associated constructs.

#### 2.7.1. Overlap with major resting-state networks

To assess spatial relationships between the INS-data and seven established resting-state networks covering the cortex, striatum, and thalamus (Yeo et al., 2011; Choi et al., 2012; Yeo, 2020), we calculated the relative and absolute distributions of ALE-derived clusters and the MACM-network within each of these reference networks (Chen et al., 2018). The relative distribution refers to the proportion of activated voxels within a reference network compared to all activated voxels, while the absolute distribution was calculated as the proportion of activated voxels compared to all voxels within a reference network. We evaluated the results using a permutation procedure (Supplement 1.5).

#### 2.7.2. Functional decoding and comparison to related meta-analytic brain networks

To reach an objective, data-driven interpretation of functional domains associated with regions found in our ALE analyses, we relied on the Neurosynth database (Yarkoni et al., 2011), the largest corpus of annotated neuroimaging data available to date (version 7; 14,371 studies). Studies in the database are mapped to *topics* generated by Latent Dirichlet Allocation (Poldrack et al., 2012) based on the frequency of topic-associated terms in the study’s full text (default: at least every 1000^th^ word annotated to the topic). We excluded 109 topics comprising mainly anatomical, disease-related, or too general terms (e.g., “resonance magnetic mechanisms”) from the 200-topic version of the database and applied two complementary functional decoding approaches. First, we decoded clusters resulting from ALE analyses based on reverse and forward inference (V. I. Müller et al., 2013) as implemented in NiMARE. For each topic and each cluster, all Neurosynth studies reporting at least one coordinate within the cluster were collected. The forward likelihood is the result of a binominal test examining whether the probability of topic-related activation in the cluster, *P*(*Activation*|*Topic*), is higher than the baseline probability of observing activation in the cluster, *P*(*Activatiori*). The reverse probability was derived from a chi-squared test assessing the probability of finding a particular topic given activation in the cluster, *P*(*Topic*|*Activation*), which was derived using Bayes’ rule. Both resulting sets of *p* values were FDR-corrected and Z transformed. Second, we adopted the “Neurosynth” approach (https://neurosynth.org/decode/) based on whole-brain spatial correlations between a brain volume of interest and topic maps derived from the spatial meta-analysis of all studies annotated to a topic. We calculated meta-analytic maps for each of 91 topics using the multilevel kernel density analysis chi-square algorithm implemented in NiMARE (Wager et al., 2007) and calculated spatial Spearman correlations to the INS Z-map after parcellation as described above. By comparison of these correlation coefficients to null distributions derived from 10,000 spatial autocorrelation-corrected topic null maps (Burt et al., 2020; Markello et al., 2022) using JuSpyce (https://github.com/LeonDLotter/JuSpyce), we estimated empirical *p* values and applied FDR correction.

To further confirm these associations, we then calculated the relative and absolute distributions of the INS-related cluster and network within meta-analytic networks of social interaction (Feng et al., 2021), ToM (Schurz et al., 2021), and predictive coding (Ficco et al., 2021) and with a previously published representation of the rTPJ in which it was parcellated into two subunits (Bzdok et al., 2013). Except for the predictive coding network, which we generated from coordinates (ALE, voxel-level *p* < .001, cluster-mass), volumetric data were obtained from the cited authors.

### 2.8. Biological contextualization

We then explored the neurobiological mechanism of INS by conducting a series of wholebrain spatial correlation analyses explicitly testing for positive associations, i.e., systems that showed their highest density in brain areas identified as subserving INS. Briefly, we first assessed relationships to neurotransmitter atlases quantified by spatial correlation analyses adjusted for spatial autocorrelation (Burt et al., 2020; Lotter & Dukart, 2022; Markello et al., 2022) and partial volume effects (Dukart & Bertolino, 2014). Second, we validated these analyses based on spatial associations with neuronal cell type distributions as obtained from human cell marker genes (Darmanis et al., 2015; Lake et al., 2016; D. Wang et al., 2018) using a neuroimaging-specific method for gene-category enrichment analysis (GCEA) (Subramanian et al., 2005) based on gene null-ensembles (Fulcher et al., 2021; Lotter et al., 2022). To assess the extent to which these molecular and cell-level systems could explain INS, we used dominance analysis (Azen & Budescu, 2003), a method that quantifies the relative contributions of each predictor to the overall explained variance in a multivariate regression model. Further GCEAs were directed at INS-associated developmental gene expression patterns, relationships to psychopathology, and INS-related molecular processes.

#### 2.8.1. Sources and processing of nuclear imaging and gene expression atlases

Invivo neurotransmitter atlases derived from nuclear imaging in various healthy adult cohorts (overall, 32 brain maps involving data from 1,360 subjects) were collected from JuSpace (https://github.com/juryxy/JuSpace) and neuromaps (https://github.com/netneurolab/neuromaps), parcellated into 116 brain regions, and Z-standardized atlaswise. Multiple atlases using the same tracers were combined by calculating the parcelwise mean weighted by the number of subjects contributing to each atlas (Hansen et al., 2022) forming 21 averaged atlases (Table S3). Parcelwise *Allen Human Brain Atlas* (ABA) mRNA expression data (https://portal.brain-map.org/) (Hawrylycz et al., 2012) were retrieved and processed with abagen (https://github.com/rmarkello/abagen/) using the default settings (Markello et al., 2021; Arnatkevičiūtė et al., 2019) (Supplement 1.6).

#### 2.8.2. Spatial associations with neurotransmitter systems

To relate the brain-wide INS distribution to molecular brain systems, spatial correlations between the INS ALE-Z map and nuclear imaging-derived brain maps were calculated as partial Spearman correlations of parcellated whole-brain data using JuSpyce. As parametric *p* values resulting from these analyses suffer from exaggerated false positive rates due to inflated degrees of freedom and spatial autocorrelations, we assessed significance by comparisons of “true” correlations to the right tails of empirically estimated null distributions of correlation coefficients derived from correlation with atlaswise null maps (5,000 iterations) (Burt et al., 2020; Lotter & Dukart, 2022; Markello et al., 2022). The resulting positive-sided empirical *p* values were FDR-corrected. To control for partial volume effects, correlations were adjusted for parcel-wise grey matter estimates derived from the MNI-152 template (Dukart et al., 2021; Dukart & Bertolino, 2014).

In sensitivity analyses, significant associations were repeated while (i) using only subcortical parcels and (ii) adjusting for functional baseline activation rate. For the second approach, a map of baseline activation rate (meta-analytic map of 14,370 Neurosynth experiments) was additionally included in the partial correlation analyses. Finally, the observed association to GABA_A_ receptors was replicated using ABA gene expression data (Supplement 1.7).

Combined with the averaged neuronal cell type maps introduced below, we finally estimated the amount of INS variance explained by neurotransmitter and neuronal cell type distributions associated with INS using dominance analysis (Azen & Budescu, 2003) as implemented in JuSpyce (Supplement 1.8).

#### 2.8.3. GCEA

GCEA was applied according to an approach specifically adopted for neuroimaging data (Fulcher et al., 2021). We adopted a previously published toolbox (https://github.com/benfulcher/GeneCategoryEnrichmentAnalysis/), originally designed for annotation of neuroimaging data to GO categories (Ashburner et al., 2000; The Gene Ontology Consortium et al., 2021), to perform GCEA on any given set of genes (ABAnnotate, https://github.com/LeonDLotter/ABAnnotate). First, a whole-brain volume (the INS ALE-Z map) and the complete ABA dataset (including mRNA expression data for 15,633 genes) were parcellated into 116 brain regions. Next, spatial autocorrelation-corrected null maps (5,000) were generated from the phenotypic data (Burt et al., 2018; Dukart et al., 2021). After matching category and ABA genes based on gene symbols, Spearman correlations between the phenotypic map, the null maps, and all mRNA expression maps were calculated. For each null map and each category, null category scores were obtained as the mean Z-transformed correlation coefficients. Positive-sided *p* values, representing the association of the phenotypic map to each category, were calculated from comparisons of the “true” category scores with the null distribution and FDR-corrected. This approach has been shown to sufficiently control false positive rates potentially caused by spatial autocorrelation present in the phenotypic data and within-category coexpression in the genetic data (Fulcher et al., 2021).

The following sets of gene-category annotations were used (Table S4): neuronal cell type markers (*PsychENCODE)* (Darmanis et al., 2015; Lake et al., 2016; D. Wang et al., 2018), genes associated with psychiatric disorders (*DisGeNET)* (Jiao et al., 2012), developmental regional gene enrichment (*BrainSpan)* (Miller et al., 2014; Grote et al., 2016), and GO biological processes (Ashburner et al., 2000; The Gene Ontology Consortium et al., 2021; Jiao et al., 2012). To aid interpretation, GO results were clustered as described in Supplement 1.9.

## 3. Results

### 3.1. Literature search and data extraction

Searching for an extensive list of INS-related terms (Supplement 1.1.1), the initial literature search resulted in 2,575 unique records from which 79 publications were found eligible for metaanalysis (Figure 2A for exclusion reasons). Finally, we included 14 hyperscanning fMRI publications (Bilek et al., 2015; Gordon et al., 2021; Koike et al., 2016; Koike, Sumiya, et al., 2019; Koike, Tanabe, et al., 2019; Miyata et al., 2021; Saito et al., 2010; Salazar et al., 2021; Shaw et al., 2018, 2020; Spiegelhalder et al., 2014; Špiláková et al., 2020; L.-S. Wang et al., 2022; Xie et al., 2020; Yoshioka et al., 2021), 8 pseudohyperscanning fMRI publications (Anders et al., 2011; Dikker et al., 2014; Kostorz et al., 2020; Liu et al., 2021, 2022; Silbert et al., 2014; Smirnov et al., 2019; Stephens et al., 2010), 54 hyperscanning fNIRS publications, and 3 pseudohyperscanning fNIRS publications [Figure 2B (interactive version available online), see Tables S1 and S2 for detailed information and fNIRS references]. INS foci coordinates were extracted from the above publications, requested from the authors, drawn from a virtual registration database (Tsuzuki et al., 2007), or derived from manual reconstruction of fNIRS probe setups (Aasted et al., 2015). Detailed information is provided in the methods, Figure 2, Supplement 1.1, Tables S1 and S2, and Figures S1 and S2.

**Figure 2:**
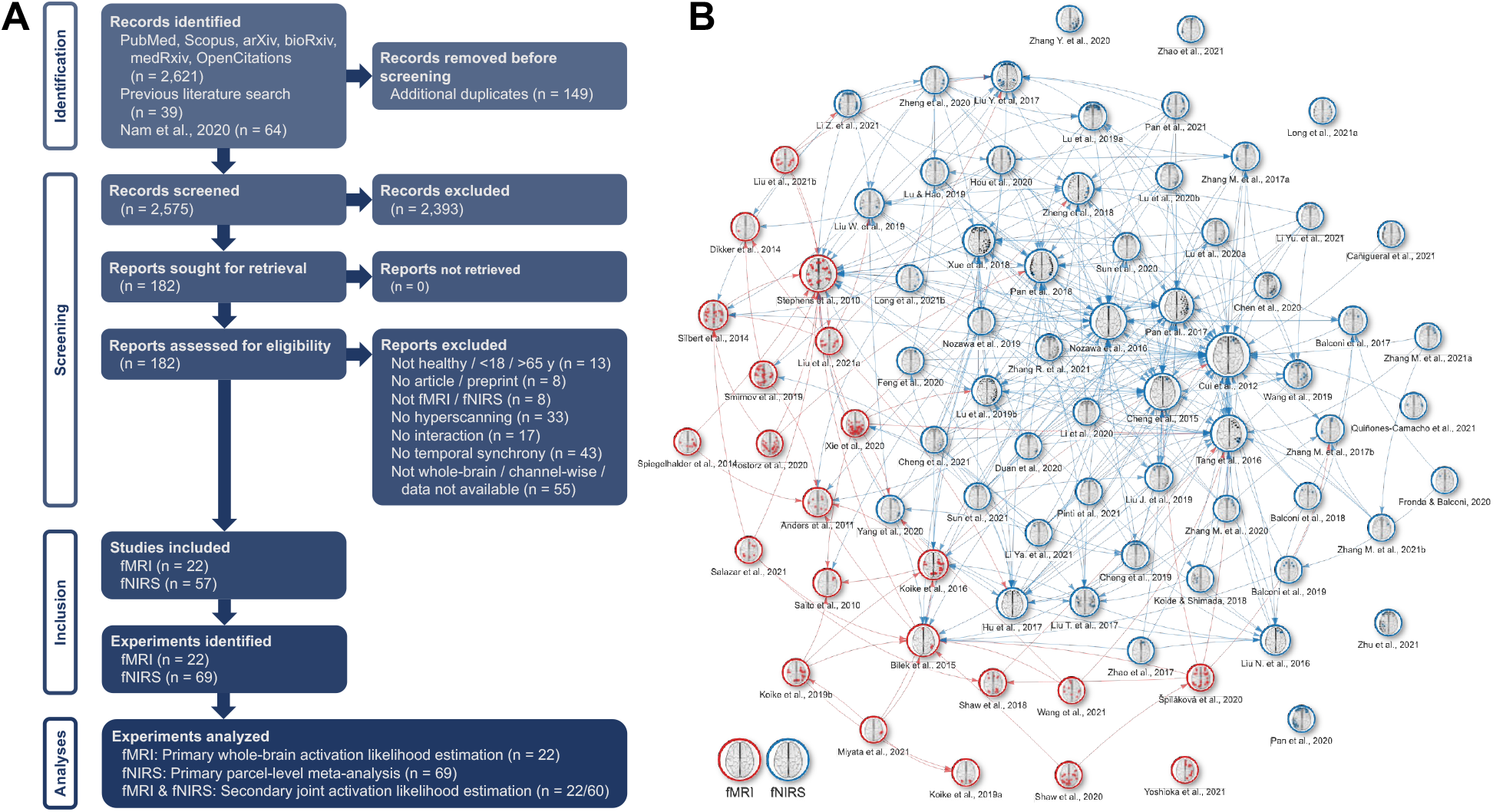
Structured literature search. **A**: Flow chart depicting the literature search process in line with the PRISMA 2020 statement. SetYouFree was used for the automatic literature search, duplicate detection and the cross-reference search. The resulting records, together with results from other sources, were submitted to Cadima to manually identify eligible studies. Note that the exclusion criteria listed in *Reports excluded* were not mutually exclusive. *Records* are entries in the publication lists resulting from the main search and screened on the abstract level, *reports* are publications screened in full, *studies* are included publications, and *experiments* are sets of data derived from independent study samples and can cover data from multiple studies. **B**: Citation network generated from OpenCitations data, including overview figures of reported INS foci and fNIRS probe setups. An interactive version with metadata for each individual study is available at https://leondlotter.github.io/MAsync/citenet. Note that the OpenCitations database only contains citations and references made openly accessible by the publishers and thus does likely not include all existing links among publications. Abbreviations: fMRI = functional magnetic resonance imaging, fNIRS = functional near-infrared spectroscopy, y = years, INS = interpersonal neural synchronization.

After taking data-reuse into account (Tables S1 and S2), 22 fMRI and 69 fNIRS experiments reporting 297 and 228 brain foci derived from data of 740 and 3,721 unique subjects were included. Task domains of these experiments varied widely, targeting communication, joint attention/action, cooperation/competition, learning, imitation, reward, and decision-making.

### 3.2. Spatial meta-analysis of fMRI and fNIRS INS experiments

To identify brain areas consistently associated with INS, we performed separate spatial metaanalyses on INS brain coordinates reported in eligible fMRI and fNIRS experiments. For fMRI data, we relied on the well-established ALE method, while for fNIRS experiments, we developed a meta-analytic procedure comparable to the ALE approach.

#### 3.2.1. Robust spatial convergence of INS revealed by fMRI hyperscanning studies

The ALE INS map revealed a mainly cortical distribution focused on right-sided parietotemporal-insular brain areas. After applying standard voxel-level thresholding (*p* < .001, uncorrected), two clusters with significant spatial convergence emerged in the rTPJ and right STG. Then, after applying a more liberal threshold (*p* < .01) (Eklund et al., 2016), we observed increased cluster sizes and an additional cluster in the right insula (Figure 3A, Table S5). Sensitivity analyses confirmed robust spatial convergence of INS in the rTPJ by showing that the cluster (i) remained after excluding pseudohyperscanning experiments (Figure S3A, Table S5), (ii) was stable against the exclusion of single experiments using a jackknife approach (12 of 22 experiments contributed relevantly to the cluster with a maximum contribution of 16%; Supplement 2.1.1; Table S1, Figure 3A), and (iii) was robust against the potential influence of publication bias (Acar et al., 2018) (*failsafe-N* of 66; Supplement 2.1.2). The right STG and insular clusters did not prove stable in these analyses.

**Figure 3:**
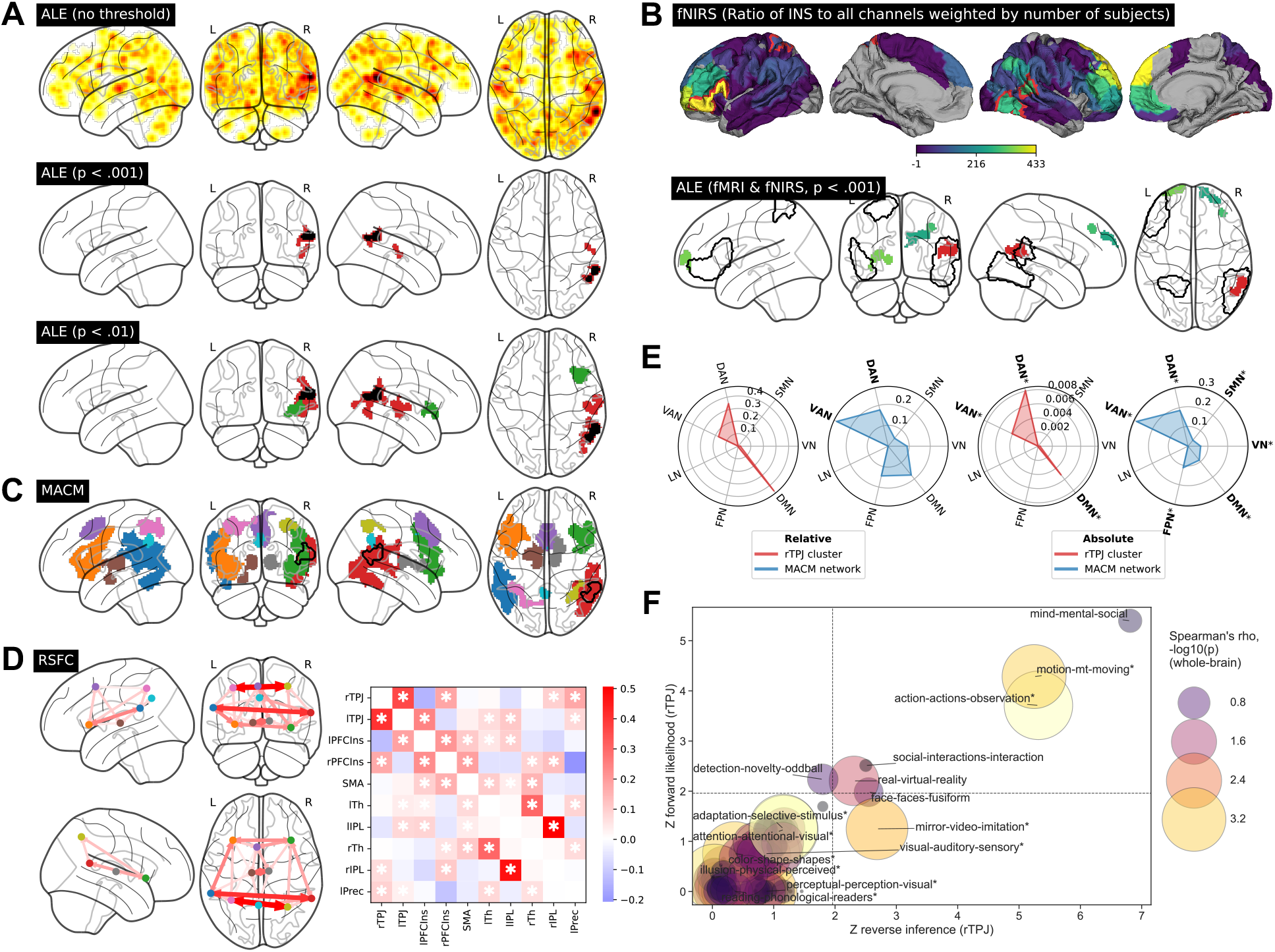
Brain-functional INS correlates resulting from fMRI and fNIRS meta-analyses with their neuronal connectivity and neurobehavioral association patterns. **A**: Results of the main fMRI INS meta-analysis. **Upper**: Unthresholded Z-map derived from ALE *p* values. **Middle/lower**: Significant INS clusters after thresholding using voxel-level thresholds of either *p* < .001 or *p* < .01. Black overlays: Spatial conjunctions of all maps derived from jackknife-analyses. **B**: **Upper**: Parcelwise fNIRS INS results. Colors: Number of “INS channels” relative to all channels sampled in the region, multiplied by the total number of subjects involved in the contributing experiments. Red outlines: Parcels showing *p* < .05 estimated in the permutation test. **Lower**: Results of a preliminary ALE analysis of the combined fMRI and fNIRS data. Black outlines: Significant parcels estimated in the fNIRS-only analysis. **C**: Meta-analytic coactivation network using the rTPJ INS cluster as a seed (black contour). **D**: Functional resting-state connectivity between MACM clusters. Colors: Semipartial Pearson correlation coefficients between clusters. Asterisks: Positive functional connections significant after Bonferroni correction (*p* < .05). **E**: Relationships of the rTPJ cluster and MACM network to major resting-state networks. *Relative:* Proportion of “INS-voxels” within a given network vs. all “INS-voxels”. *Absolute:* Proportion of “INS-voxels” within a given network vs. all voxels within the network. Bold print: *p* < .05. *: *p* < .05. **F**: Functional decoding of INS-related activation using Neurosynth topics. Three alternative approaches are presented: x-axis: Z-transformed, FDR-corrected *p* values derived from decoding of the rTPJ cluster using reverse inference, P(Topic|Activation); y-axis: similar but with forward likelihood, P(Activation|Topic) > P(Activation). Point sizes and colors: Nonparametric *p* values according to Spearman correlations between whole-brain INS distribution and meta-analytic topic maps (*: *q* < .05). Dashed lines: Alpha level of *q* < .05. All FDR-corrected significant topics are annotated. Abbreviations: fMRI = functional magnetic resonance imaging, fNIRS = functional near-infrared spectroscopy, INS = interpersonal neural synchronization, ALE = activation likelihood estimation, MACM = meta-analytic connectivity modeling, RSFC = resting-state functional connectivity, r/lTPJ = right/left temporoparietal junction, rSTG = right superior temporal gyrus, rIns = right insula, r/lPFCIns = right/left prefrontal cortex-insula, SMA = supplementary motor area, r/lTh = right/left thalamus, r/lIPL = right/left inferior parietal lobule, lPrec = left precuneus, DAN/VAN = dorsal/ventral attention network, SMN = somatomotor network, VN = visual network, DMN = default mode network, FPN = frontoparietal network, LN = limbic network.

#### 3.2.2. Spatial convergence of INS revealed by fNIRS hyperscanning studies

Meta-analytic evaluation of fNIRS data revealed four significant parcels covering the right inferior temporal gyrus (4/19 INS channels/total channels, *n* = 1,339 subjects, *p* = .017), left inferior frontal gyrus (IFG; 11/56, *n* = 2,205, *p* = .020) and superior parietal gyrus (3/5, *n* = 154, *p* = .047), as well as right STG that overlapped with the fMRI-derived rTPJ cluster (5/35, *n* = 2,145, *p* = .048; Figures 3B and S5A, Table S5). None of the derived exact *p* values survived false discovery rate (FDR) correction. An exploratory ALE analysis, including combined INS coordinates from 22 fMRI and 60 fNIRS experiments, resulted in four significant clusters covering the rTPJ, left anteroventral superior frontal gyrus, and right middle and superior frontal gyri (Figure 3B, Table S5). The fNIRS studies contributed to both the prefrontal clusters and the rTPJ cluster (Tables S1 and S2). Evaluation of alternative indices derived from fNIRS data further pointed to bilateral prefrontal and left temporoparietal brain regions (Supplement 2.2.1, Figure S5A and B, Table S6). Sensitivity analyses, accounting for bias in study selection and spatial uncertainty of fNIRS data, demonstrated generally comparable patterns. However, concerning the fNIRS-only meta-analysis, the left superior frontal cluster showed the highest stability, while, in the combined fNIRS-fMRI meta-analysis, left superior frontal and rTPJ were the most stable locations (Supplement 2.2.2–3, Tables S1, S2, and S5, Figure S5).

Summarizing, in line with prior findings and models of INS, we identified the rTPJ as a robust and task domain-independent hub region of INS supported by both fMRI and fNIRS data. FNIRS meta-analysis additionally indicated involvement of the left inferior PFC in INS.

### 3.3. INS-related neuronal connectivity and biobehavioral association patterns

To establish the functional context of the identified INS hub region within large-scale brain networks and biobehavioral domains, we conducted a set of brain- and task-functional association analyses capitalizing on different open neuroimaging databases (Figure 1). First, a MACM network of brain regions likely to co-activate with the rTPJ hub was constructed from the BrainMap database (Laird et al., 2011) and RSFC patterns within this network were evaluated (Van Essen et al., 2013; Whitfield-Gabrieli & Nieto-Castanon, 2012). To then explore the functional context of the observed activation patterns and aid interpretation, we characterized relationships to established brain-wide resting-state networks (Yeo et al., 2011; Chen et al., 2018), biobehavioral domains in the Neurosynth database (Yarkoni et al., 2011), and previously published meta-analytic networks of INS-associated constructs, i.e., social interaction (Feng et al., 2021), empathy and ToM (Schurz et al., 2021), and predictive coding (Ficco et al., 2021). Finally, we assessed how the rTPJ cluster related to a parcellation of the rTPJ (Bzdok et al., 2013).

#### 3.3.1. Task-based coactivation and resting-state connectivity networks

148 BrainMap studies reported at least one activation focus within the rTPJ cluster. The meta-analytic coactivation network involved primarily bilateral frontotemporal cortical regions, with the largest clusters placed on bilateral TPJs, insulae and dorsolateral PFCs, supplementary motor areas, and thalami (Figure 3C, Table S5). Within this network, RSFC was strongest between temporoparietal clusters, while subcortical regions showed functional connections primarily to insulae but not to the TPJ hub regions (Figure 3D). An additional analysis controlling for baseline activation probability also indicated the TPJs as unique hub regions of the observed INS-related network (Supplement 2.3.1, Figure S3B). Comparing whole-brain patterns of INS-ALE maps and MACM maps, bilateral TPJs, insulae, and dorsal PFCs showed the highest activation likelihood in both maps, indicating a possible role of the MACM network in INS beyond the rTPJ activation (Supplement 2.3.2, Figure S3C). In line with interregional connectivity patterns, the rTPJ cluster and the associated coactivation network showed the strongest spatial associations to the default mode and attention resting-state networks (Supplement 2.4, Figure 3E).

#### 3.3.2. Functional decoding of INS-related networks

We observed significant associations between the rTPJ and topics related to ToM, action, observation, and social interaction. On the whole-brain level, the strongest associations were found with topics related to attention and sensory domains (Figure 3F, Table S7). In line with that, INS-associated activation showed a general alignment with (affective) ToM and social interaction networks, and relatively greater overlap with the posterior rTPJ subunit, which itself had previously been related to ToM and social cognition (Bzdok et al., 2013). While the predictive coding network did not include the rTPJ, it strongly resembled the INS-related MACM network (Figure S4).

In summary, spatial associations analyses embedded meta-analytic INS results in the context of large-scale brain networks mainly related to attentional, sensory, and mentalizing processes. While the TPJs again emerged as hub regions, we cannot exclude the possibility that INS may also be present in the extended INS-related network involving mainly the insulae, PFCs, and potentially thalami.

### 3.4. Relationships of INS with neurophysiological brain systems

To extend our meta-analytic approach from the macroscale level of brain regions to meso- and microscale neurophysiological functions, and thereby identify biological processes potentially underlying INS, we conducted a series of whole-brain spatial association analyses. We first assessed relationships between INS and neurotransmitter systems by spatial correlations with neurotransmitter atlases (Dukart et al., 2021; Markello et al., 2022; Hawrylycz et al., 2012). Using GCEA, we identify INS-related neuronal cell types (Darmanis et al., 2015; Lake et al., 2016; D. Wang et al., 2018), tested for potential gene-mediated associations between INS and psychopathology (Piñero et al., 2020), quantified the regional enrichment of INS-related genes across neurodevelopment (Miller et al., 2014; Grote et al., 2016), and identified INS-related molecular processes (Ashburner et al., 2000; Fulcher et al., 2021; The Gene Ontology Consortium et al., 2021).

#### 3.4.1. Spatial associations with neurotransmitter systems and neuronal cell types

We observed significant positive spatial associations between INS and the distributions of GABAergic (GABA_A_) and glutamatergic (mGluR5) receptors as well as between INS and synaptic density (SV2a). Without FDR-correction, INS was further associated with serotonergic (5-HT_2A_) and cholinergic components (M1; Figures 4A and S6). The remaining neurotransmitter atlases were not significantly associated (Figure S6, Table S8). Furthermore, GCEA indicated significant associations with a specific excitatory neuron class, *Ex3* (*Z* = .36, *p* < .001, *q* < .001), and two classes of inhibitory neurons, *In5* (*Z* = .12, *p* < .001, *q* = .012) and *In6* (*Z* = .19, *p* = .001, *q* = .032; Figures 4B, Table S9). In the original publication that identified the applied cell markers (Lake et al., 2016), *Ex3* was enriched in visual brain areas and cortical layer IV, and classified as granule neuron. *In5* and *In6* were widely distributed, concentrated in layers II/III (*In5*) and IV/V (*In6*), and the latter was identified as the parvalbumin-expressing interneuron subclass. Dominance analysis including all FDR-corrected significant nuclear imaging and cell type maps indicated an overall explained variance of 31.4% with the strongest relative contributions by *Ex3* (26.2%), followed by GABA_A_ (19.8%) and mGluR5 (18.5%; Figure S8).

**Figure 4:**
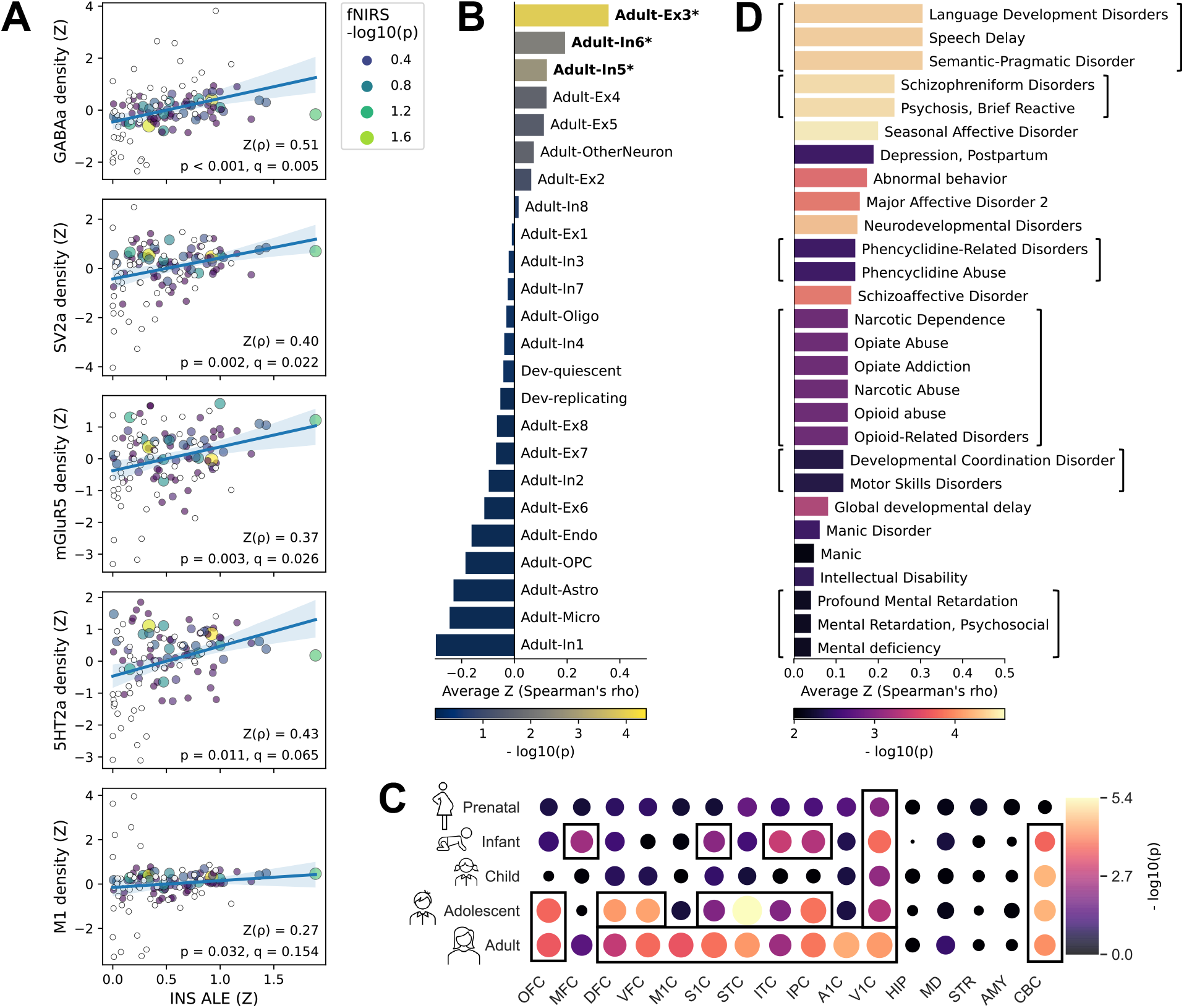
Spatial associations of INS to neurotransmitter receptor and synaptic density distributions, as well as genetic markers of neuronal cell types, brain development and psychiatric disorders. **A**: Correlation of the whole-brain INS Z-map (x-axis) with in vivo nuclear imaging-derived neurotransmitter receptor and synaptic density (SV2a) distributions. Statistics: Partial Spearman correlations between INS and nuclear imaging maps adjusted for local grey matter volume with exact *p* values estimated via nonparametric permutation-corrected for spatial autocorrelation. Points: Value pairs associated representing 116 whole-brain parcels. Point size and color: *p* values derived from the fNIRS INS meta-analysis as shown in the upper panel of Figure 2B. The two yellow and the rightmost points correspond to the significant fNIRS atlas parcels. White points: Parcels not included in the fNIRS analysis. Lines: Linear regression lines with 95% confidence intervals. **B, C, and D**: Spatial associations of INS to postmortem mRNA expression distributions estimated by GCEA. Bar plots: Associations of INS with neuronal cell type markers (**B**) and disease markers (**D**). Bars: Average Z-transformed Spearman correlation coefficient of all genes annotated to a category. Bar color: Uncorrected *p* values. For cell types (**B**), all categories are shown, and those significant at FDR-corrected *p* < .05 are highlighted. For disease markers (**D**), only significant categories are shown. Brackets: Categories comprising exactly the same genes. **C**: INS-related regional developmental gene enrichment across five developmental stages (y-axis) and 16 brain regions (x-axis). Point size: Average r-to-Z transformed correlation coefficient. Point color: Uncorrected *p* values. Rectangles: Categories significant after FDR correction. Abbreviations: INS = interpersonal neural synchrony, ALE = activation likelihood estimation, OFC = orbital frontal cortex, MFC = anterior (rostral) cingulate (medial prefrontal) cortex, DFC = dorsolateral prefrontal cortex, VFC = ventrolateral prefrontal cortex, M1C = primary motor cortex, S1C = primary somatosensory cortex, STC = posterior (caudal) superior temporal cortex, ITC = inferolateral temporal cortex, IPC = posteroventral (inferior) parietal cortex, A1C = primary auditory cortex, V1C = primary visual cortex, HIP = hippocampus, MD = mediodorsal nucleus of thalamus, STR = striatum, AMY = amygdaloid complex, CBC = cerebellar cortex, GCEA = gene category enrichment analysis.

Further sensitivity analyses targeting neurotransmitter associations showed (i) that additional adjustment for functional baseline activation rate generally led to a slight increase in effect sizes (Table S8), (ii) that only the associations with GABA_A_ and SV2a remained significant after exclusion of subcortical parcels to estimate effects of a potential cortical-subcortical density gradient in the neurotransmitter data (Table S8), and (iii) that the GABAA-association was replicated in mRNA expression data (Figure S7, Table S8). The latter pointed to a relationship between INS and genes coding for the α1-GABA_A_ receptor subunit and parvalbumin (Supplement 2.5.1).

#### 3.4.2. Spatial associations with developmental gene expression, genetic disease markers, and molecular processes

Significant developmental gene enrichment was found in 29 categories that revealed a gene expression pattern that was pronounced in adult cortical sensory brain areas but was detectable from the postnatal stage (Figure 4C, Table S9). Of note, we found no significant enrichment prenatally, or in subcortical areas. Furthermore, INS was strongly related to genes previously associated with neurodevelopmental disorders and secondary with affective disorders (9/16 and 5/16 unique significant categories, respectively; Figure 4D, Table S9). Finally, semantic clustering analyses (Reijnders & Waterhouse, 2021) based on 474 GO categories spatially associated with the INS distribution (Table S9) indicated processes related to neuron and general cell development, and neuronal signal transmission (Table S10, Figure S8).

Concluding, our results pointed towards a neurophysiological basis of INS consisting of major inhibitory and excitatory neuronal systems involved in sensory processing. In line with that, INS-related genes (i) were most expressed in cortical sensory brain areas starting in postnatal brain development and (ii) have previously been related to primarily neurodevelopmental disorders.

## 4. Discussion

In recent years, synchronization of brain activities between interacting partners has been acknowledged as a central mechanism by which we foster successful social relationships as well as a potential factor involved in the pathogenesis of diverse neuropsychiatric disorders. Based on the results generated by our multimodal data fusion approach (see Figure 5), we hypothesized that human INS is tightly linked to social attentional processing, subserved by the rTPJ as a sensory-integration hub at the brain system level, and potentially facilitated by GABA-mediated E/I balance at the neurophysiological level.

**Figure 5:**
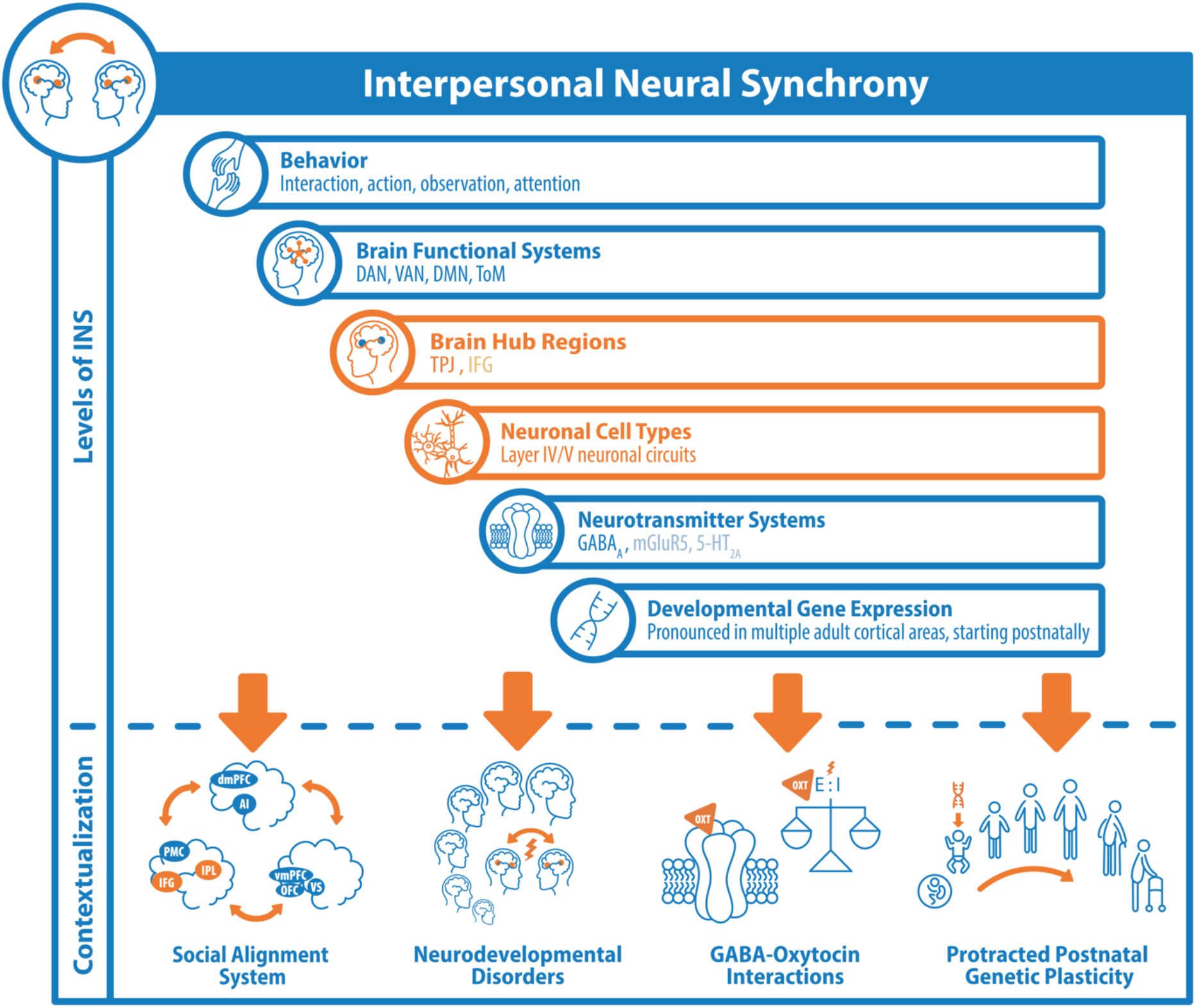
Summary of results and hypotheses. Abbreviations: INS = interpersonal neural synchronization, DAN/VAN = dorsal/ventral attention network, DMN = default mode network, ToM = theory of mind network, TPJ = temporoparietal junction, IFG = inferior frontal gyrus, dm/vmPFC = dorsomedial/ventromedial prefrontal cortex, AI = anterior insula, PMC = premotor cortex, IPL = inferior parietal lobule, OFC = orbitofrontal cortex, VS = ventral striatum, OXT = oxytocin, E/I = excitation/inhibition balance.

### 4.1. Brain functional systems facilitating INS

Our results confirmed the central role of the rTPJ and of the left IFG in the facilitation of INS. Both regions have been proposed as central hubs of the *mutual social attention system*, thought to allow for “coupling” between interacting partners (Gvirts & Perlmutter, 2020), and the *observation-execution/alignment* subunit of the *social alignment system* (Shamay-Tsoory et al., 2019). On the whole-brain level, the connection to the *alignment system* was further supported by our data indicating a clear association of INS to biobehavioral domains, such as “motion”, “action”, “observation”, “imitation”, and “attention”. On the brain regional level, we also demonstrated links between the rTPJ and its functional connected network with the brain’s attention networks. Here, attention might generally enhance the processing of relevant information, regulate the overall cortical responsiveness (Lakatos et al., 2008), and more specifically increase the computational weight of prediction error units via synaptic gain enhancement (Kok et al., 2012). Both, the rTPJ as well as the associated network, were further linked to the default mode network as well as networks associated with ToM and social interaction and showed strong associations with ToM-related terms (e.g., “mind”, “social”, and “interaction”). Consistent with the previously reported twofold functional specialization of the rTPJ (Bzdok et al., 2013), our findings highlight the link between interpersonal synchronization and both *alignment of behavior* (Shamay-Tsoory et al., 2019) as well as *alignment of mental states* (i.e., ToM) (Gallotti et al., 2017). Thus, our ALE findings are largely in line with the assumptions of the mutual prediction theory suggesting that on the brain network level, INS might reflect the sum of brain activity involved in encoding self-behavior (e.g., right-sided anterior insula) (Seth et al., 2012), other’s behavior and mental state (e.g., rTPJ) and dynamic interpersonal alignment (e.g., left IFG).

While the most robust effect was found in the rTPJ, the left IFG became evident as an additional region of convergence when findings from fNIRS hyperscanning experiments were included. The IFG might be involved in both simple mirroring of behavior and brain activities (Hasson & Frith, 2016) and facilitating dynamic interpersonal alignment, e.g., by containing motor representations of actions (Shamay-Tsoory, 2021; Shamay-Tsoory et al., 2019). While all three components of the *social alignment system* have been proposed to be involved in INS (Shamay-Tsoory et al., 2019), we found evidence only for INS in the *alignment* subsystem but not in the *misalignment-detection* and *reward* sub-systems. Although these latter systems might also be involved in social alignment processes (Shamay-Tsoory et al., 2019), our finding could indicate that they might not necessarily exhibit synchronized neural activity. The question of whether systems underlying misalignment detection and experienced reward mainly *form and motivate* biobehavioral synchrony or actually *become activated simultaneously* in interacting individuals will have to be addressed in future research.

Given our approach of searching for the common spatial correlate of INS across a wide range of study types, tasks, analysis protocols, and social interaction settings, two aspects deserve further consideration. On the one hand, based on our results, we cannot draw conclusions about situation-specific spatial INS patterns that might occur bound to specific interaction settings. On the other hand, the identified brain regions might be considered *core regions* of INS that simultaneously activate in interacting humans *independent* of the specific interaction context. Reiterating the aforementioned connection to attentional processing and the *social alignment system*, and referring to the following discussion on potential neurophysiological mechanisms, we hypothesize that a core component of social interaction must be a system that supports the reciprocal alignment of attention and sensory processing between interacting people. INS in the core brain areas supporting these cognitive functions might represent this mutual alignment process, thus representing the correlate of successful interpersonal communication.

### 4.2. Comparing fMRI and fNIRS approaches to measuring INS

Given the currently available data and the meta-analytic methods applied, our study is not suited to rule out the possibility that INS is present in a larger network but was not detected in these other areas. Relatedly, we also found no evidence for prefrontal INS based on fMRI hyperscanning data alone. Comparing specific confounds affecting the comparison between fNIRS and fMRI data, we noted that the fNIRS data had only limited cortex coverage and prefrontal regions were relatively oversampled. Furthermore, fNIRS data shows higher spatial uncertainty resulting mainly from variations in head shape, virtual registration methods, and post-hoc reconstruction of probe coordinates (Aasted et al., 2015; Cooper et al., 2012; Tsuzuki et al., 2007; Tsuzuki & Dan, 2014) and lower image quality due to limited methodological standardization (Tachtsidis & Scholkmann, 2016). On the other hand, upright body posture has been shown to affect body physiology, brain activation, and cognitive performance, potentially increasing the ecological validity of fNIRS to capture neurophysiological processes underlying social interactions (Thibault & Raz, 2016). In addition to several methodological differences inherent to fMRI and fNIRS (Czeszumski et al., 2020), variation may also stem from differing experimental designs and settings. First, experimental designs allowed generally only for limited face-to-face contact in both modalities, however only fNIRS experiments could establish real-life interactional settings. Second, while both hyperscanning techniques covered a similarly broad spectrum of task domains with *communication* tasks being most prevalent, a bias was found toward more *joint attention* tasks for fMRI experiments and toward *cooperation/competition* conditions for fNIRS experiments. Keeping in mind these limitations but considering that our fNIRS hyperscanning dataset far exceeds its fMRI pendant in terms of sample size (3,721 versus 740 subjects), the prefrontal activation observed in fNIRS data might indeed arise from interpersonal collaboration paradigms (Czeszumski et al., 2022) which should be tested in future fMRI hyperscanning research.

### 4.3. Future pathways for the hyperscanning research field

We identified several pathways that can be built upon in the future. Methodologically, it seems mandatory for the field to move toward standardized and reproducible data acquisition, task protocols, and analysis pipelines (Yücel et al., 2021). Dissecting the definition of *interpersonal synchronization* both in a methodological and in a conceptual sense (Schirmer et al., 2021) will help us delineate the brain regional and behavioral domain-dependent nuances of INS. To (i) ensure that the phenomenon of INS goes beyond the representation of shared sensory environments, (ii) delineate the interaction between behavioral interpersonal synchronization and INS, and (iii) facilitate effective interpretation and formulation of predictions for future research, carefully designed experiments, including multimodal (behavior, body, and brain) data collection and integrative data analysis within a neurocognitive framework, constitute necessary steps for the future (Hamilton, 2021). As the current state of the fMRI hyperscanning field did not allow us to conduct metaanalyses that differentiate task domains or examine the influence of, e.g., sex, gender, and age, the impact of these factors on INS will have to be clarified in future meta-analyses. Most studies included here focused on temporal synchrony between homologous brain regions or on the univariate coherence between one region and multiple other regions. Analogous to the development of “traditional” fMRI research, moving toward network-based analyses in which functionally connected networks are tracked *across* subjects (Gerloff, Konrad, et al., 2022) will elucidate the role of the rTPJ and its functional connections between individuals. Furthermore, while the current meta-analysis of fNIRS data was focused on spatial information, our openly accessible dataset could easily be extended to include statistical information, thus enabling regional effect size-based meta-analyses (Czeszumski et al., 2022).

### 4.4. GABA-mediated E/I balance as a potential mechanism underlying INS

To generate hypotheses on the neurophysiological processes underlying INS, we developed and applied state of the art data fusion methods drawing robust spatial associations between wholebrain INS patterns and potentially underlying cellular, molecular, and genetic systems (Hansen et al., 2022; Dukart et al., 2021; Markello et al., 2022; Fulcher et al., 2021; Lotter et al., 2022; Lotter & Dukart, 2022). We note, however, that our inferences are based on spatial correlation analyses involving heterogeneous data obtained from often relatively small samples and, thus, must be treated cautiously. These analyses, by no means, provide a reliable basis for causal claims and cannot replace nuclear imaging methods, brain stimulation protocols, pharmacological investigations, or histological approaches. We nevertheless consider these analyses to be sufficient to provide a basis for future hypothesis-driven research based on the currently available hyperscanning data.

We found converging evidence for an association of spatial INS patterns with GABA_A_ receptors and layer IV/V interneuron density, suggestive of (“fast-spiking”) parvalbumin-expressing interneurons. These interneurons constitute the largest group of cortical inhibitory neurons (Tremblay et al., 2016), contribute to feedback and feedforward inhibition, and are critically involved in the generation of network oscillations (Hu et al., 2014). In accordance with the association between INS and attention systems, GABAergic neurotransmission has previously been linked to general attentional processing and to visual attentional selectivity in particular (Lockhofen & Mulert, 2021). Furthermore, parvalbumin-expressing interneurons, along with layer V pyramidal cells, have been shown to express 5-HT_2A_ receptors which we also found spatially related to INS (Andrade & Weber, 2010). Together with *Ex3* (“granule”) excitatory neurons, which explained the largest amount of spatial INS information in the present study and were also located in cortical layer IV (Lake et al., 2016), these INS-associated neuron classes may form *thalamocortical feedforward inhibition circuits* (Tremblay et al., 2016) between thalamocortical afferents (Sherman & Guillery, 2002), GABAergic interneurons, and pyramidal cells which have crucial roles in encoding spatial and temporal sensory information (Gabernet et al., 2005; Tremblay et al., 2016). Relatedly, layer IV neurons have previously been linked to computations of prediction errors (Bastos et al., 2012). Finally, in line with the potential relevance of thalamocortical neuronal circuits, the INS-associated meta-analytic network involved the bilateral thalami as the sole subcortical structures.

Here, we could not confirm the involvement of oxytocin and dopamine in neurobiobehavioral synchrony that was hypothesized previously (Feldman, 2017; Gvirts & Perlmutter, 2020; Mu et al., 2016). In contrast, we consider our findings to be more in line with the hypothesis that INS relies on neurobiological mechanisms similar to those previously reported for within-brain synchronization processes: *GABA-mediated E/I balance* (Sears & Hewett, 2021). Given links between GABAergic interneurons and local gamma oscillations (Gonzalez-Burgos & Lewis, 2008) as well as between local oscillations, GABA concentration, and long-range functional connectivity (Stagg et al., 2014; Rajkumar et al., 2021), we hypothesize that INS might be maintained by GABA-regulated signal transmission that relies on thalamocortical pathways transferring sensory information obtained during social interaction to an INS-associated cortical network centered in the rTPJ. Importantly, while we focused on blood oxygenation signals, electrophysiological studies also localized the source of gamma power synchronization in the TPJ (Hoehl et al., 2021; Kinreich et al., 2017). Of note, indirect involvement of oxytocin might be plausible given that both oxytocin and GABA signaling may mediate *E/I balance* in the contexts of social interaction and brain development (Lopatina et al., 2018), that GABA_A_ receptors might have oxytocin binding sites (Gough, 2015), and that oxytocin regulates GABA_A_ neurosteroid binding and expression patterns (Kaneko et al., 2016; Koksma et al., 2003).

The indirect nature of our evidence for INS mediation by GABAergic signaling and E/I balance requires thorough confirmation or falsification with experimental data. Future electrophysiological, and especially magnetoencephalography-based (Watanabe et al., 2022) hyperscanning experiments could investigate the role of E/I balance in the development of INS. GABAergic involvement as well as regulatory mechanisms could be tested using pharmacological challenges with safe oxytocin-regulating or GABA-regulating agents or multibrain stimulation protocols to show causal roles in the regulation and maintenance of human INS.

### 4.5. Associations to neurodevelopment and neurodevelopmental disorders

We demonstrated that genes which’s spatial expression pattern aligned with the INS distribution were expressed particularly in cortical sensory brain areas starting at the postnatal stage. The GABAergic system undergoes significant changes in early postnatal development, switching from excitatory to inhibitory action, thus establishing E/I balance (Lopatina et al., 2018), enabling intrapersonal cortical synchronization (Warm et al., 2022) and potentially also INS. If the observed developmental gene expression pattern reflects the development of this neurophysiological system, it might indicate long-lasting plasticity in this system; however, it is also compatible with ideas about neurobiological instantiation through early postnatal infant-caregiver interactions (Feldman, 2017).

We observed gene-mediated spatial associations of INS with psychopathology, i.e., primarily with disordered language development, general neurodevelopment, and schizophrenia spectrum disorders and, secondarily, with affective disorders. Indeed, both developmental and affective disorders have been suggested as disorders of social interaction (Schilbach, 2016; Kruppa et al., 2021) and a prior study reported reduced INS during interaction between people with and without autism spectrum disorder (Tanabe et al., 2012). Moreover, autism and schizophrenia spectra have repeatedly been linked to disrupted GABA signaling and E/I balance (Gonzalez-Burgos & Lewis, 2008; Tang et al., 2021; Lopatina et al., 2018) as well as thalamotemporal functional and structural dysconnectivity (Ameis & Catani, 2015; Woodward et al., 2017), and GABAergic agents have been shown to affect both clinical presentation and brain E/I balance in patients with autism (L. Zhang et al., 2020). Our findings provide further evidence for a pathophysiological role of (disrupted) INS in these disorders, with relevance for both the identification of social biomarkers and the development of neurobiologically based treatment strategies.

### 4.6. A multilevel model of human INS

Taken together, our findings suggest a model of human INS bridging multiple levels of organization from social interaction to molecular processes (Figure 5). Interindividual synchrony in humans, as in many group-living species, amplifies social rapport and affective bonds and arises both intentionally and spontaneously (Hoehl et al., 2021). Our actions broadcast our behaviors through the environment, i.e., by motion, touch, gesture, mimics, speech, or sounds. Our brain’s sensory systems, subserved by neuronal attention networks, rhythmically sample information from the environment and can detect these actions. Synchronizing the actions of one individual with the perceptions of another and vice versa might then lead to interpersonal synchronization of brain oscillations (Hasson & Frith, 2016). From these sensory inputs, we not only observe and predict the actions of our interaction partner but also infer mental states. The rTPJ may serve as the brain’s hub for processing attention information and integrating this information into a broader context, communicating with brain networks relevant for controlling selective attention, reactive action, inference of mental states, and affective responses (Downar et al., 2000; Bzdok et al., 2013). Neurophysiologically, these interpersonally synchronized oscillations may, within each brain, arise from GABA-mediated E/I balance, receiving sensory input from thalamic afferences. In an attempt to incorporate prior hypotheses on the involvement of oxytocin and dopamine or social reward processes (Feldman, 2017; Gvirts & Perlmutter, 2020; Shamay-Tsoory et al., 2019), one might now speculate that these aspects could act as influencing factors on the rTPJ, functionally associated brain networks, or GABA-mediated neuronal oscillations, without directly partaking in INS (Lopatina et al., 2018; Mu et al., 2016). Relatedly, dysregulation of this multilevel representation of INS might be involved in the pathophysiology of neurodevelopmental disorders (Ameis & Catani, 2015; Gonzalez-Burgos & Lewis, 2008; Lopatina et al., 2018; Tang et al., 2021; Woodward et al., 2017). If further validated and fully understood, our findings may lay the foundation of a neurobiologically based concept of “typical” INS, with relevance for the study of human social development, communication, and interpersonal learning processes. Understanding how INS works will also help us understand how its mechanisms fail, which might have critical implications for how we view and treat disordered social interaction on both individual levels and societal levels.

Closing and taking a general perspective, our methodological approach demonstrated the value of multimodal association analyses not only for psychiatric research but also to advance current models of cognition in typically developing populations. We designed this study with reuse of our resources in mind to provide a path for similar future endeavors.

## 5. Acknowledgements

We thank everyone involved in the creation of the original publications that constitute the basis of this work. Special thanks go to those authors who responded to our requests and provided additional information and data to their publications. We thank Julia Camilleri and Simon Eickhoff for providing an initial version of the functional decoding analyses as well as Jacqueline Gädtke and Marius Kaufmann for their help with extraction of fNIRS publication data.

LDL was supported by the Federal Ministry of Education and Research, Germany (BMBF) and the Max Planck Society, Germany (MPG).

