## Supplementary materials for "Revealing the Neurobiology Underlying Interpersonal Neural Synchronization with Multimodal Data Fusion"

### 1 Supplementary Materials and Methods

#### 1.1 Literature search and data extraction

##### *1.1.1 Search string*

```
(([hyperscanning] OR
  [neural entrainment] OR
  (([brain-to-brain] OR
    [interbrain] OR
    [inter-brain] OR q
    [interpersonal]) AND
   ([synchrony] OR
    [synchronization] OR
    [coupling] OR
    [connectivity] OR
    [networks] OR
    [alignment] OR
    [coherence]))) AND
([fNIRS] OR
 [NIRS] OR
 [near-infrared spectroscopy] OR
 [fMRI] OR
 [MRI] OR
 [magnetic resonance imaging]))
```

##### *1.1.2 Study inclusion criteria*

- **Article type**: original research, peer-reviewed or preprint, English language (exclude: conference paper, review, meta-analysis, opinion)
- **Subjects**: human, healthy, between 18 and 65 years
- **Imaging Technology**: fNIRS or fMRI
- **Methodological study design**: “hyperscanning” or “pseudohyperscanning”
- **Task**: uni- or bidirectional interaction between ≥ two subjects
- **Analysis**: temporal synchrony between data derived from hemodynamic signals of ≥ two subjects
    - **fMRI**: analysis of data on whole-brain level (exclude: “region of interest” approaches)
    - **fNIRS**: analysis of channel-wise signals (exclude: data analysis after averaging data from multiple channels to form a “region of interest”)
- **Data availability**:
    - **fMRI**: reporting of coordinates in Montreal Neurological Institute (MNI) or Talairach space, or availability of coordinates or 3D volumes on request
    - **fNIRS**: reporting of coordinates in MNI or Talairach space, or availability of coordinates or fNIRS probe setups on request, or reporting of ≥ one coordinate in

international 10-10 system, alignment to a reference plane, and usage of a square probe setup, thus allowing for reconstruction of the probe positioning

##### *1.1.3 Extracted study information*

- **General**: authors, publication year, title, journal, identifier
- **Sample:** number of subjects, number of dyads, male/female ratio, relationship between interacting subjects, mean age, age standard deviation, age range
- **Method:** imaging technology, hyperscanning or pseudohyperscanning, general analysis technique, applied contrasts
    - **fMRI:** repetition time, time lag, analysis frequency band, analysis level (voxelwise, independent component analysis, seed-to-voxel, whole-brain parcellation), analysis software
    - **fNIRS:** device, wavelength of emitted light, sampling rate, analyzed data (HbO, HbR), analysis frequency band, time lag, optode array format and positioning
- **Task and setting:** general domain, exact applied task, setting
    - **fMRI:** (live) audio/video contact
    - **fNIRS:** positioning of subjects to each other (side-by-side, face-to-face, separated)
- **Coordinate data:** coordinates in MNI or Talairach space, author contact information if no coordinates were available
    - **fNIRS:** source of coordinates (virtual registration, digitization, MRI-registration)

##### *1.1.4 Sample data extraction*

Sample data were extracted as reported in included studies. If a study reported exclusion of subjects, e.g., due to low data quality, sample sizes after exclusion of these subjects were retained. When, e.g., in case of pseudo-hyperscanning fMRI and some fNIRS studies, demographic data was provided for the study sample and a small group of “special” participants who were scanned repeatedly, sample sizes were determined by the overall sum of participants, while the number of dyads equaled the number of formed dyads, and age mean and standard deviation were calculated as weighted averages. For example, if a study included n = 20 students (mean age 20 years) and n = 2 teachers (mean age 24 years), forming n = 20 dyads with teachers being scanned repeatedly, we set the sample size to 22, the number of dyads to 20, and the mean age to 20.37 years.

When a study split its sample in multiple groups with independent subjects and independently analyzed data, these groups were included as separate experiments. When sample information (i.e., male/female ratio, age) were reported only for the whole sample but not for subgroups, this information was applied to all subgroups at the risk of small inaccuracies (which only apply to the results Tables S1 and S2 but have no influence on our meta-analytic results).

When samples between different studies overlapped, i.e., data derived from the same subjects was analyzed and published multiple times, these data, and reported brain coordinates, were concatenated into one experiment. If it was most likely that (according to the descriptions of sample characteristics, tasks, and the publishing authors), a sample was analyzed twice but this was not mentioned in the papers and the authors did not answer requests, we chose a conservative approach and concatenated respective data into one experiment. When concatenating data, the lowest reported sample size of all individual studies was used for the resulting "experiment" to be included in meta-analyses.

##### *1.1.5 fMRI coordinate data extraction*

Coordinates provided in MNI space were not modified. Coordinates in Talairach space were converted to MNI using the Lancaster transform (*1*).

When authors sent thresholded volumetric imaging data, we extracted a maximum of three peak coordinates with a minimum distance of 10mm from each cluster containing at least 10 voxels.

When authors sent independent component analysis results in the form of volumetric data depicting independent components, we Z-normalized these maps while excluding zero voxels, applied a cluster-threshold of $Z > 2.5$ and retained only clusters exceeding a volume of 0.1% of all non-zero in-brain voxels. From all studies applying independent component analysis, for each reported or estimated cluster, a maximum number of three peak coordinates with a minimum distance of 10mm were retained for meta-analysis.

When data was analyzed using a whole-brain parcellation, for each parcel showing INS, the center of mass was used as focus coordinates.

When authors analyzed INS not only for homologous brain areas (voxels, parcels, independent components) but in a pairwise fashion for the whole brain, foci representing all connected regions were retained for meta-analysis (e.g., when significant INS was found between right temporoparietal junction of subject 1 and right insula of subject 2, we would include foci representing both regions).

##### *1.1.6 fNIRS coordinate data extraction and reconstruction*

For studies that did not report coordinates, we obtained MNI coordinates according to the following workflow:

When studies reported to have used the virtual registration method (*2*) and used a common static rectangular optode placeholder, we used coordinates provided by Tsuzuki et al. (http://brain-lab.jp/wp/?page_id=58). If studies did not report to have used the virtual registration method but their description of optode placements matched one of the probe setups for which virtual registration results were available, we used these for the analyses [e.g., Cui et al., 2012 (*3*) matched Fpz_low3_HorSag_3x5].

In some cases, a probe set configuration reported by study A was used in study B as well and study B was published by authors from the same research group or University. When coordinates for study B were available, we used these for study A as well [e.g., for Liu et al, 2019 (*4*), we used the temporoparietal coordinates from Lu et al, 2020b (*5*)].

When this was not possible, we reconstructed the probe sets based on the spatial information provided by the studies using AtlasViewer (https://openfnirs.org/software/homer/) and registered it to a scaled version of the Colin27 MNI-atlas. Since head size information were not provided by the studies in question, we could not register the Colin27 atlas in AtlasViewer to study-specific head sizes but had to rely on a standard head size for all reconstructions. In order to get an appropriate approximation of the average head shape, we calculated the average head sizes for white and

Asian women and men (560.01mm) based on data provided by Ball (*6*) (https://repository.tudelft.nl/islandora/object/uuid:2d038418-8923-4605-92e8-ca3df57ea731) and Harrison & Robinette (*7*) (https://apps.dtic.mil/sti/pdfs/ADA406674.pdf). To be suited for reconstruction, studies had to report at least two anatomical landmarks of the EEG 10-10 or 10-5 system. We rated the "reconstructability" of studies as *high* if EEG channel positions for every single optode were reported, *medium* if at least two EEG channel positions for each optode array were given, or l*ow* if one EEG channel position and some additional spatial information about the alignment of the probe set, e.g., "15° angle between probe patches and transverse plane", was reported. When only one landmark but no additional information or no information at all was reported, these studies were excluded.

When studies analyzed and reported fNIRS data in voxel space and reported significant voxel-level MNI coordinates, we obtained coordinates according to the workflow lined out above and retained, for each reported significant voxel, coordinates of the three fNIRS channels most closely located to this each voxel.

#### 1.2 Calculation of the fail-safe-N

The robustness against publication bias of each cluster was estimated as the fail-safe-N (*8*). For each cluster, noise experiments with sample sizes and foci numbers drawn from the actual included fMRI experiments were generated. Foci coordinates for these noise studies were randomly drawn from all gray matter voxels excluding those in the brain quadrant where the respective cluster's center of mass was located. Based on preset minimum and maximum numbers of added noise experiments (the number of contributing experiments and the number of noise studies needed to reach a minimum contribution of 10% of studies) (*8*), we iteratively searched for the minimum number of noise experiments needed to render the cluster insignificant while applying standard voxel-level and cluster-level thresholds ($p < .001$ and $p < .05$). The resulting number mirrors the hypothetical minimum of negative studies that could have "remained in the file drawer" necessary for the respective cluster to fail significance thresholds.

#### 1.3 Modelling of spatial uncertainty in fNIRS meta-analyses

To incorporate spatial uncertainty of fNIRS data in fNIRS meta-analyses, we iteratively recalculated both the parcel-wise fNIRS meta-analysis and the combined ALE after randomization of fNIRS coordinates within a 10mm radius and a strongly constrained cortical MNI-152 template (1,000 iterations). The radius choice was based on previous data showing a localization error of 18mm in preregistered model-based fNIRS-fMRI registration (*9*, *10*).

To evaluate the fNIRS-only meta-analytic results, we calculated, for each parcel, the percentage of sub-threshold $p$ values ($p < .05$) relative to the total number of iterations as well as the median $p$ value resulting from all iterations.

The combined ALE was repeated with randomized coordinates in the same fashion as the main analysis. However, we adopted the cluster mass threshold estimated from the maximum cluster size distribution of the original fNIRS-fMRI ALE instead of recalculating individual thresholds for each fNIRS coordinate randomization iteration to reduce the required computational power.

We then evaluated the results by calculating the proportion of thresholded ALE maps for which nonzero ALE values were present within the clusters derived from the main combined fMRI-fNIRS ALE.

##### 1.4 Resting-state fMRI data processing

To estimate the functional connectivity pattern present within the task-based coactivation network, $N = 120$ unrelated subjects from the Human Connectome Project S900 release (*11*) were randomly selected from age and gender groups (20 females and 20 males from each age group: 22–25, 26–30, and 31–35 years). We used the S900 extensively preprocessed volumetric data (ICA-FIX denoised) thoroughly described in the reference manual (*12*). Further processing in CONN (https://web.conn-toolbox.org/) included: resampling of the data to a 3mm isotropic MNI-152 template, concatenation of the first two resting-state sessions (resulting in 30 min), linear detrending and bandpass filtering (0.01–0.08Hz), averaging of voxelwise timeseries across MACM clusters, and calculation of semipartial correlations between clusters (*13*). After exclusion of subjects exceeding framewise displacement cutoffs of 2mm translation or 2 degree rotation ($n = 5$), we calculated two-sided one-sample t-tests against zero on the r-to-Z transformed semipartial correlation coefficients representing each (directed) functional connection. To identify the strongest functional connections, the resulting $p$ value matrix was thresholded at FWE-corrected $p < .05$ (Bonferroni) and only positive connections were interpreted to exclude potential spurious negative connections introduced through noise regression (*14*).

##### 1.5 Relative and absolute distributions within major resting-state networks

To characterize the ALE clusters and the associated MACM networks in respect to their spatial overlap with established brain-wide resting-state networks (*15–17*), we adopted a frequently used method characterizing the absolute and relative distributions of our target volumes across these networks (*18*). The *relative distribution* refers to the proportion of activated voxels within a reference network compared to all activated voxels, while the *absolute distribution* is calculated as the proportion of activated voxels compared to all voxels within a reference network. To estimate significance of these spatial associations, we then permuted the input coordinates (1,000 repetitions; fMRI INS coordinates and rTPJ-associated BrainMap coordinates, respectively) within a grey-matter mask (MNI-152, thresholded at > .2), performed ALE analyses on these coordinates, and iteratively reassessed the resting-state network overlaps to generate null distributions of the overlap metrics. From these null distributions, we calculated empirical $p$ values denoting the probability of a false positive finding under the null hypothesis of random spatial localizations of INS/MACM coordinates. The $p$ values were FDR-corrected for each metric across the combined rTPJ and MACM results. To reduce computational cost, we did not run the full cluster mass permutation procedure as in the main ALE/MACM analyses but extracted the cluster mass thresholds from the primary analyses and used these to perform cluster-level inference on the null data after applying a voxel-level threshold of $p < .001$.

##### 1.6 Processing of Allen Brain Atlas mRNA expression data

Regional microarray expression data were obtained from 6 post-mortem brains (1 female, age range 24.0–57.0 years, mean age 42.50 ± 13.38 years) provided by the Allen Human Brain Atlas (https://human.brain-map.org) (*19*). Data were processed with the abagen toolbox (version 0.1.3; https://github.com/rmarkello/abagen) (*20*) using a volumetric atlas in MNI space covering 116 cortical and subcortical brain regions (*21*, *22*).

First, microarray probes were reannotated using data provided by Arnatkevičiūtė et al. (*23*); probes not matched to a valid Entrez ID were discarded. Next, probes were filtered based on their expression intensity relative to background noise (*24*), such that probes with intensity less than the background in ≥ 50% of samples across donors were discarded, yielding 31,569 probes. When multiple probes indexed the expression of the same gene, we selected and used the probe with the most consistent pattern of regional variation across donors [i.e., differential stability (*25*)], calculated with:

$$\Delta_S(p) = \frac{1}{\binom{N}{2}} \sum_{i=1}^{N-1} \sum_{j=i+1}^{N} \rho\left[B_i(p), B_j(p)\right]$$

where $p$ is Spearman's rank correlation of the expression of a single probe, p, across regions in two donors $B_i$ and $B_j$, and N is the total number of donors. Here, regions correspond to the structural designations provided in the ontology from the Allen Human Brain Atlas. The MNI coordinates of tissue samples were updated to those generated via non-linear registration using the Advanced Normalization Tools (https://github.com/chrisfilo/alleninf). Samples were assigned to brain regions in the provided atlas if their MNI coordinates were within 2mm of a given parcel. All tissue samples not assigned to a brain region in the provided atlas were discarded. Inter-subject variation was addressed by normalizing tissue sample expression values across genes using a robust sigmoid function (*26*):

$$x_{norm} = \frac{1}{1 + exp - \left(\frac{(x - \langle x \rangle)}{IQR_x}\right)}$$

where $\langle x \rangle$ is the median and $IQR_x$ is the normalized interquartile range of the expression of a single tissue sample across genes. Normalized expression values were then rescaled to the unit interval:

$$x_{scaled} = \frac{x_{norm} - \min(x_{norm})}{\max(x_{norm}) - \min(x_{norm})}$$

Gene expression values were then normalized across tissue samples using an identical procedure. Samples assigned to the same brain region were averaged separately for each donor and then across donors, yielding a regional expression matrix with 116 rows, corresponding to brain regions, and 15,633 columns, corresponding to the retained genes.

##### 1.7 Validation analyses for in-vivo INS-$GABA_A$ associations

To validate the spatial association found between INS and the $GABA_A$ receptor distribution, the following list of GABA-related genes was collected (*27*, *28*), and data were extracted from the

Allen Brain atlas: GABRA1, GABRA2, GABRA3, GABRA4, GABRA5, GABRB1, GABRB2, GABRB3, GABRG1, GABRG2, GABRG3, GABRD, GABRE, GAD1, GAD2, PVALB, SST, VIP, CCK, NPY, CALB1, CALB2, NOS1, RELN, ADRB2, LHX6, TAC1, TAC3, TAC4, SLC6A1, SLC6A13, SLC6A12, SLC32A1, GABBR1, GABBR2.

Following genes were not available after preprocessing of Allen Brain Atlas data: GABRA6, GABRP, GABRQ, GABRR1, GABRR2, GABRR3, LAMB5, SLC6A11.

The expression data from all genes were pairwise correlated using Spearman correlations and the correlation matrix was clustered using the default unsupervised hierarchical clustering method implemented in scipy (*29*), based on Euclidean distances and estimating cluster proximity using the nearest point algorithm. Normalized expression values of all genes within a cluster were Z-standardized, averaged and correlated with the INS ALE-Z map using JuSpyce (*30*) as described in the main methods section (partial Spearman correlations, adjusted for local gray matter volume). Resulting $p$ values were FDR-corrected.

##### 1.8 Dominance analysis on nuclear imaging and neuronal cell type data

Dominance analysis is designed to determine the relative contribution of each predictor in a multivariate regression model to the overall explained variance as an intuitive measure of predictor importance (*31*). It does so by calculating all subset models of a multivariate linear model, i.e., re-calculating the regression analysis with all possible combinations of predictors. We implemented the method in JuSpyce to estimate three different dominance statistics of which we focused on two. *Total dominance* is the average contribution of a predictor to the total $R^2$ across all subset models and can be interpreted as the amount of explained variance of a predictor relative to the total explained variance. For comparison, we further evaluated *individual dominance*, which is the $R^2$ resulting from the univariate regression of a single predictor on the target variable and thus quantifies the amount of information explained by a predictor alone. As the main goal of this analysis was to quantify the amount of INS variance explained by PET and cell type maps, we submitted only those maps to dominance analysis that showed significant positive relationships to INS (FDR-corrected) in the previous analyses. PET data was atlas-wise parcellated, Z-standardized, and averaged across atlases using the same tracer. Neuronal cell type maps were generated by Z-standardizing each parcelwise gene expression vector and calculating the average gene expression per cell type category. Parcelwise gray matter volume was regressed out of the INS map and each predictor map before performing the regression analysis to align with the main correlation analyses.

##### 1.9 Clustering and visualization of INS-associated GeneOntology categories

Results from GeneOntology (GO) gene-category enrichment analyses as obtained from ABAnnotate (*32*) were further clustered based on semantic similarity using GO-Figure! which was described in detail elsewhere (*33*, *34*). Briefly, the dimensionality of the list of significantly associated GO terms is reduced by calculating pairwise semantic similarity scores based on (i) the distance of two GO terms within the directed acyclic organization of GO categories and (ii) the frequency of each GO term in a large database of genes (equaling “specificity” of each GO term).

Based on an arbitrary similarity threshold, GO terms are then grouped into clusters, a representative term is selected, and a multidimensional scaling algorithm is applied to generate a two-dimensional visualization. Here, we chose a liberal threshold of $\geq .2$ to capture the overall biological functions of the identified GO clusters at the cost of specificity.

### 2 Supplementary Results

#### 2.1 FMRI meta-analysis sensitivity analyses

##### *2.1.1 Influence of individual experiments*

To estimate the influence of individual experiments on observed interpersonal neural synchrony (INS) clusters, we applied a jacknife-approach and recalculated ALE analyses while iteratively excluding one experiment at a time.

Using these data, we estimated that 12 of 22 experiments contributed to the right temporoparietal junction (rTPJ) cluster, with a maximum contribution of 16% using the conservative threshold ($p < .001$). In contrast, the right superior temporal and right insula clusters were driven by a lower number of studies with corresponding stronger contributions (Table S1). Accordingly, the spatial conjunction of all jackknife-derived thresholded maps proofed only the right temporoparietal junction (rTPJ) cluster as stable against the influence of individual experiments (Figure 2A).

##### *2.1.2 Risk of publication bias*

Coordinate-based meta-analytic techniques, such as ALE, only allow for limited possibilities for structured bias assessment. However, to assess the robustness of ALE results against publication bias, calculation of the cluster-wise *fail-safe-N* was proposed (*8*). Adopting this method, we tested how many noise-experiments generated with characteristics similar to the original INS data we could add to the INS ALE analysis before the originally observed clusters were no longer significant.

Based on the inclusion of a minimum of 12 (the number of contributing experiments) and a maximum of 98 ($\frac{12}{98+22} = 10\%$) of noise-experiments, we observed a fail-safe-N of 66 for the rTPJ cluster indicating that even if 66 "negative" experiments were not available for this analysis due to publication bias, we would still have observed spatial convergence of INS in the rTPJ. By contrast, the cluster in the right superior temporal gyrus did not survive the inclusion of the defined minimum of 12 additional noise-experiments.

#### 2.2 FNIRS meta-analysis sensitivity analyses

##### *2.2.1 Alternative "fNIRS-Indices"*

In addition to the "INS-to-all-channel-ratio" weighted by subject number we also evaluated the ratio weighted by the number of experiments as well as the raw count of "INS-channels" as parcelwise indices. We however emphasize that only the first index which is reported in the main paper incorporates all available information (presence of "INS-channels" in relation to the times a region was sampled and number of subjects)

Based on exact $p$ values estimated from permutation of parcel-channel assignments, we observed significant results for left inferior-anterior prefrontal regions across all evaluated measures. Additionally, while evaluation of the channel ratio weighted by subjects indicated right temporal brain regions, results based on the INS channel count pointed to left temporal and parietal regions and based on the channel-ratio weighted by the number of experiments we found a significant right frontomedial parcel (Figure S5). Again, no $p$ value survived false discovery rate (FDR) correction.

###### *2.2.2 Modelling of spatial uncertainty*

To account for spatial uncertainty of fNIRS data in all fNIRS meta-analyses, we iteratively repeated the parcelwise fNIRS meta-analysis as well as the ALE analysis with spatially randomized fNIRS coordinates (1cm radius, 1,000 iterations) and assessed the percentages of iterations in which significant results at the original brain locations were observed.

In the fNIRS-only meta-analysis, we generally observed a high sensitivity of cluster significance towards randomization of parcel-coordinates. Especially the relatively small rTPJ parcel was highly sensitive, showing sub-threshold $p$ values in only 7.9% of iterations ($median\ p = .28$) in the main evaluated fNIRS index. The highest consistency was found for the right inferior temporal parcel showing persistence in 48% of iterations ($median\ p = .052$). The overall results for all fNIRS indices are displayed in Table S6 and Figure S5. Concerning the combined fMRI-fNIRS ALE, coordinate randomization least affected the rTPJ and the left and right superior frontal clusters which emerged in 100% of iterations, while the right middle frontal cluster was found in 89.1% of iterations.

###### *2.2.3 Restricted set of fNIRS experiments*

To confirm our fNIRS-results, we restricted the included experiments to a more conservative selection including only experiments explicitly contrasting interaction with control, rest, or randomization conditions and recalculated all analyses.

We observed a generally comparable pattern but in the parcellation-based fNIRS-only meta-analysis only prefrontal parcels showed significance for the "INS channel ratios" weighted by number of subjects or experiments while the fNIRS evaluation based on the raw INS channel count remained stable ($p$ uncorrected; Figure S5). In the combined fMRI-fNIRS ALE, we mainly observed a reduction of cluster sizes (Tables S1 and S2).

##### 2.3 MACM sensitivity analyses

###### *2.3.1 Controlling for baseline activation probability*

Specific coactivation likelihood estimation (SCALE) is an alternative meta-analytic connectivity modeling (MACM) algorithm controlling for the baseline probability of observing coactivation independent of the chosen seed (*35*). We calculated a SCALE analysis using all studies in the BrainMap database as baseline (N = 3,098), to estimate the brain regions most uniquely coactivating with the rTPJ.

Here, we observed the left TPJ region, and to a lesser extent right insula, as specific functional connections of the rTPJ, further indicating the TPJs as hub regions of the observed INS-related network (Figure S3B).

###### *2.3.2 Spatial alignment between INS-ALE and MACM activation patterns*

To estimate whether the MACM network mirrors a brain-wide activation pattern that was already present in the original meta-analytic INS map beyond the rTPJ activation, we correlated the parcellated whole-brain maps derived from both analyses.

We observed a spatial alignment pattern driven by bilateral TPJs, insulae as well as dorsolateral prefrontal cortices, possibly indicating a role of these regions in INS but a lack of power to detect these as areas of spatial convergence in the main meta-analysis (Figure S3C).

##### 2.4 Spatial relationships to established resting-state networks

We characterized the rTPJ cluster and the MACM network in respect to their spatial overlap with established brain-wide resting-state networks (*15*). We adopted a frequently used method characterizing the *absolute* and *relative distributions* of our target volumes across these networks (see method). We then permuted the input coordinates within all gray matter voxels (1,000 iterations), tested the resulting maps for meta-analytic convergence, and calculated absolute and relative distributions for each null map. From these null distributions, empirical $p$ values were estimated.

The result is shown in Figure 2E. Looking at the absolute distributions, the rTPJ cluster was significantly associated with the default mode network ($p = .001, q = .002$), the dorsal attention ($p = .001, q = .002$) and the ventral attention network ($p = .026, q = .040$). The MACM network showed a widespread pattern (default mode, dorsal attention, ventral attention, frontoparietal, and somatomotor network: $p = .001, q = .002$). Concerning the relative distributions, the rTPJ cluster did not show significant associations while, in confirmation of the relationship between INS and attention networks, associations significant at an uncorrected alpha-level were observed between the MACM network and the ventral ($p = .004, q = .056$) and dorsal ($p = .047, q = .243$) attention networks.

##### 2.5 Neurotransmitter-associations sensitivity analyses

###### *2.5.1 GABA$_A$-related mRNA expression*

To validate the strong association found to the GABA$_A$ receptor, and further clarify which subtype of GABA$_A$ receptors may drive the association, we collected a list of GABA-related genes (*27*, *28*), clustered these according to their spatial co-expression profiles (Figure S7A), and assessed spatial correlation patterns between INS and cluster-wise mRNA expression data.

Two GABA gene clusters were significantly associated with INS ($Z = .43, p = .007, q = .014$ and $Z = .45, p = .005, q = .014$). In line with prior data on the molecular target of the applied GABA$_A$ tracer (*27*), one of these clusters comprised the α1-GABA$_A$ receptor subunit (GABRA1) and parvalbumin (PVALB), of which the latter is considered a marker of fast-spiking parvalbumin-expressing interneurons, the largest group of cortical inhibitory neurons (*36*) (Figure S7B, clusters 2 and 3; Table S8).

152. The Gene Ontology Consortium, S. Carbon, E. Douglass, B. M. Good, D. R. Unni, N. L. Harris, C. J. Mungall, S. Basu, R. L. Chisholm, R. J. Dodson, E. Hartline, P. Fey, P. D. Thomas, L.-P. Albou, D. Ebert, M. J. Kesling, H. Mi, A. Muruganujan, X. Huang, T. Mushayahama, S. A. LaBonte, D. A. Siegele, G. Antonazzo, H. Attrill, N. H. Brown, P. Garapati, S. J. Marygold, V. Trovisco, G. dos Santos, K. Falls, C. Tabone, P. Zhou, J. L. Goodman, V. B. Strelets, J. Thurmond, P. Garmiri, R. Ishtiaq, M. Rodríguez-López, M. L. Acencio, M. Kuiper, A. Lægreid, C. Logie, R. C. Lovering, B. Kramarz, S. C. C. Saverimuttu, S. M. Pinheiro, H. Gunn, R. Su, K. E. Thurlow, M. Chibucos, M. Giglio, S. Nadendla, J. Munro, R. Jackson, M. J. Duesbury, N. Del-Toro, B. H. M. Meldal, K. Paneerselvam, L. Perfetto, P. Porras, S. Orchard, A. Shrivastava, H.-Y. Chang, R. D. Finn, A. L. Mitchell, N. D. Rawlings, L. Richardson, A. Sangrador-Vegas, J. A. Blake, K. R. Christie, M. E. Dolan, H. J. Drabkin, D. P. Hill, L. Ni, D. M. Sitnikov, M. A. Harris, S. G. Oliver, K. Rutherford, V. Wood, J. Hayles, J. Bähler, E. R. Bolton, J. L. De Pons, M. R. Dwinell, G. T. Hayman, M. L. Kaldunski, A. E. Kwitek, S. J. F. Laulederkind, C. Plasterer,

M. A. Tutaj, M. Vedi, S.-J. Wang, P. D'Eustachio, L. Matthews, J. P. Balhoff, S. A. Aleksander, M. J. Alexander, J. M. Cherry, S. R. Engel, F. Gondwe, K. Karra, S. R. Miyasato, R. S. Nash, M. Simison, M. S. Skrzypek, S. Weng, E. D. Wong, M. Feuermann, P. Gaudet, A. Morgat, E. Bakker, T. Z. Berardini, L. Reiser, S. Subramaniam, E. Huala, C. N. Arighi, A. Auchincloss, K. Axelsen, G. Argoud-Puy, A. Bateman, M.-C. Blatter, E. Boutet, E. Bowler, L. Breuza, A. Bridge, R. Britto, H. Bye-A-Jee, C. C. Casas, E. Coudert, P. Denny, A. Estreicher, M. L. Famiglietti, G. Georghiou, A. Gos, N. Gruaz-Gumowski, E. Hatton-Ellis, C. Hulo, A. Ignatchenko, F. Jungo, K. Laiho, P. Le Mercier, D. Lieberherr, A. Lock, Y. Lussi, A. MacDougall, M. Magrane, M. J. Martin, P. Masson, D. A. Natale, N. Hyka-Nouspikel, S. Orchard, I. Pedruzzi, L. Pourcel, S. Poux, S. Pundir, C. Rivoire, E. Speretta, S. Sundaram, N. Tyagi, K. Warner, R. Zaru, C. H. Wu, A. D. Diehl, J. N. Chan, C. Grove, R. Y. N. Lee, H.-M. Muller, D. Raciti, K. Van Auken, P. W. Sternberg, M. Berriman, M. Paulini, K. Howe, S. Gao, A. Wright, L. Stein, D. G. Howe, S. Toro, M. Westerfield, P. Jaiswal, L. Cooper, J. Elser, The Gene Ontology resource: enriching a GOld mine. *Nucleic Acids Research*. **49**, D325–D334 (2021).

### 4 Supplementary Tables

| Publication | DOI | Experiment ID | Type | Area | Task | A/V | Contrast | Subjects | Dyads | Female | Age | | Space | Method | TR | Lag | Band | Foci | ALE cluster contributions: fMRI (p < .001) | | fMRI (p < .01) | | | fMRI+fNIRS (p < .001) | | | |
|---|---|---|---|---|---|---|---|---|---|---|---|---|---|---|---|---|---|---|---|---|---|---|---|---|---|---|---|
| | | | | | | | | [N] | [N] | proportion | mean [y] | SD [y] | | | | | | [N] | rTPJ [%] | rSTG [%] | rTPJ [%] | rSTG [%] | rIns [%] | rTPJ [%] | lSFG [%] | rSFG [%] | rMFG [%] |
| **Anders et al., 2011** | 10.1016/j.neuroimage.2010.07.004 | Anders 2011 | pseudo | emotion | express emotions/ emphasize | V | prediction of perciever's from sender's voxel-wise activity | 12 | 6 | 0.50 | 23.00 | n.a. | MNI | Seed | 2000 | 0 | n.a. | 7 | **0.00** | **0.00** | **0.00** | **0.00** | **0.00** | **0.00** | **0.00** | **0.00** | **0.05** |
| **Bilek et al., 2015** | 10.1073/pnas.1421831112 | Bilek 2015 1 | hyper | joint attention | show target via button press | V | (dyad-INS during interaction > no interaction) > random-INS | 26 | 13 | 0.62 | 24.50 | 4.60 | MNI | wb-ICA | 1550 | 0-1.55 | n.a. | 2 | **5.63** | **0.00** | **4.08** | **0.00** | **0.00** | **4.67** | **0.00** | **0.07** | **0.00** |
| | | Bilek 2015 2 | hyper | joint attention | show target via button press | V | (dyad-INS during interaction > no interaction) > random-INS | 50 | 25 | 1.00 | 23.40 | 3.30 | MNI | wb-ICA | 1550 | 0-1.55 | n.a. | 1 | **9.28** | **0.00** | **5.66** | **0.00** | **0.00** | **7.03** | **0.00** | **0.00** | **0.00** |
| **Dikker et al., 2014** | 10.1523/JNEUROSCI.3796-13.2014 | Dikker 2014 | pseudo | communica tion | isten to image description | A | speaker/listener-INS > 0 | 10 | 9 | 0.80 | 25.05 | 5.00 | MNI | ts-VW | 1500 | 0 | n.a. | 1 | **0.00** | **0.00** | **0.00** | **0.00** | **0.00** | **0.00** | **0.00** | **0.00** | **0.00** |
| **Koike et al., 2016/ 2019b** | 10.1016/j.neuroimage.2015.09.076 | Koike 2016 1 3 + 2019b | hyper | joint attention | look at target cued by partner's gaze | V | dyad-INS during JA > random-INS | 64 | 32 | 0.59 | 22.40 | 5.08 | MNI | wb-VW | 2500 | 0 | .01-.08 | 20 | **12.70** | **0.00** | **13.22** | **0.03** | **11.35** | **11.51** | **0.00** | **0.00** | **0.00** |
| | 10.1093/scan/nsz087 | Koike 2016 2 | hyper | joint attention | look at partner & think about feelings | V | dyad-INS during mutual gaze without JA task > random-INS | 30 | 15 | 0.47 | 20.60 | 2.92 | MNI | wb-VW | 2500 | 0 | .01-.08 | 1 | **2.49** | **0.00** | **3.44** | **0.00** | **0.00** | **1.55** | **0.00** | **0.00** | **0.00** |
| **Koike et al., 2019a** | 10.1523/ENEURO.0284-18.2019 | Koike 2019a | hyper | joint attention | look at partner & think about feelings | V | dyad-INS during mutual gaze with live video > delayed video | 28 | 14 | 0.36 | 21.80 | 2.17 | MNI | wb-VW | 1000 | 0 | >.008 | 6 | **0.00** | **0.00** | **0.00** | **0.00** | **0.00** | **0.00** | **0.00** | **0.00** | **0.00** |
| **Kostorz et al., 2020** | 10.1016/j.neuroimage.2020.116659 | Kostorz 2020 | pseudo | learning | watch origami folding & memorize | V | intructor-observer-INS > 0 | 29 | 28 | 0.52 | 27.20 | n.a. | MNI | wb-VW | 590 | 0 | >.001 | 27 | **8.92** | **0.00** | **10.73** | **0.00** | **0.00** | **7.94** | **0.00** | **0.00** | **0.00** |
| **Liu et al., 2021a/ 2021b** | 10.1007/s00429-021-02271-2 10.1101/2021.03.02.433669 | Liu 2021a + 2021b | pseudo | communica tion | listen to auto-biographic story | A | speaker/listener-INS > 0 | 33 | 32 | 0.52 | 23.00 | n.a. | MNI | wb-Atlas | 2000 | 0 | >.008 | 15 | **0.00** | **20.70** | **0.05** | **25.06** | **0.00** | **0.00** | **0.00** | **0.00** | **0.00** |
| **Miyata et al., 2021** | 10.1016/j.neuroimage.2021.117916 | Miyata 2021 | hyper | joint action | show or imitate facial expression | V | dyad-INS during imitation > random-INS | 32 | 16 | 0.69 | 22.42 | n.a. | MNI | wb-VW | 3500 | 0 | >.008 | 6 | **0.05** | **0.00** | **1.72** | **0.00** | **0.00** | **0.02** | **0.00** | **0.00** | **0.00** |
| **Saito et al., 2010** | 10.3389/fnint.2010.00127 | Saito 2010 | hyper | joint attention | look at target cued by partner's gaze | V | dyad-INS during JA > random-INS | 38 | 19 | 0.00 | 24.50 | 4.10 | MNI | wb-VW | 3000 | 0 | >.008 | 3 | **0.00** | **0.00** | **0.00** | **0.00** | **19.60** | **0.00** | **0.00** | **0.00** | **0.00** |
| **Salazar et al., 2021** | 10.1016/j.neuroimage.2020.117697 | Salazar 2021 | hyper | joint action | try to say the same word | A | dyad-INS during JAct > random-INS | 44 | 22 | 0.45 | 26.80 | 3.80 | MNI | ts-ICA | 2000 | 0 | >.008 | 6 | **0.00** | **0.00** | **0.00** | **0.00** | **0.00** | **0.00** | **0.00** | **0.00** | **0.00** |
| **Shaw et al., 2018** | 10.1038/s41598-018-29233-9 | Shaw 2018 | hyper | decision making | iterated ultimatum game | no | dyad-INS during ultimatum game > control | 38 | 19 | 0.00 | 24.60 | 3.70 | MNI | wb-VW | 2000 | 0 | σ = 60 | 12 | **0.00** | **0.00** | **0.00** | **0.00** | **8.66** | **0.00** | **1.01** | **0.00** | **0.19** |
| **Shaw et al., 2020** | 10.1371/journal.pone.0232222 | Shaw 2020 | hyper | coop/comp | interactive pattern game | no | dyad-INS during coop/comp > random-INS | 54 | 27 | 0.00 | 34.90 | 10.08 | MNI | ts-ICA | 2300 | 0 | σ = 50 | 19 | **7.79** | **20.46** | **5.06** | **9.92** | **0.00** | **6.95** | **0.00** | **0.00** | **0.00** |
| **Silbert et al., 2014** | 10.1073/pnas.1323812111 | Silbert 2014 | pseudo | communica tion | listen to auto-biographic story | A | speaker/listener-INS > random correlation | 14 | 11 | n.a. | 30.50 | n.a. | TAL | wb-VW | 1500 | 0 | "high-pass" | 22 | **10.53** | **0.33** | **9.80** | **8.26** | **13.48** | **7.69** | **0.00** | **0.00** | **0.00** |
| **Smirnov et al., 2019** | 10.1002/hbm.24736 | Smirnov 2019 | pseudo | communica tion | listen to auto-biographic story | A | speaker/listener-INS > random correlation | 18 | 16 | 1.00 | 25.00 | n.a. | MNI | wb-VW | 1700 | 0 | .04-.07 | 34 | **0.01** | **4.24** | **0.15** | **11.65** | **0.00** | **0.02** | **0.01** | **0.00** | **0.00** |
| **Spiegelhalder et al., 2014** | 10.1016/j.bbr.2013.10.015 | Spiegelhalder 2014 | hyper | communica tion | listen to auto-biographic story | A | dyad-INS during speak/listen > control | 22 | 11 | 1.00 | 27.20 | 2.90 | MNI | Seed | 2660 | 0 | >.008 | 9 | **8.48** | **0.65** | **5.39** | **1.43** | **0.00** | **6.52** | **0.00** | **0.00** | **0.00** |
| **Špiláková et al., 2020** | 10.1002/hbm.24861 | Spilakova 2020 | hyper | coop/comp | interactive pattern game | no | dyad-INS during coop/comp > random-INS | 38 | 19 | 0.42 | 22.44 | 1.90 | MNI | ts-ICA | 2000 | 0 | σ = 50 | 18 | **15.99** | **21.90** | **10.09** | **10.87** | **0.00** | **12.69** | **0.00** | **6.12** | **15.00** |
| **Stephens et al., 2010** | 10.1073/pnas.1008662107 | Stephens 2010 | pseudo | communica tion | listen to story | A | speaker-listener-INS > listener-listener-INS | 14 | 22 | n.a. | 25.50 | n.a. | TAL | wb-VW | 1500 | 0-4 | "high-pass" | 20 | **8.43** | **0.01** | **6.39** | **8.05** | **12.15** | **6.43** | **0.00** | **0.01** | **0.00** |
| **Wang et al., 2022** | 10.1101/2021.07.21.452832 | Wang 2021 | hyper | coop/comp | "treasure chest" game | no | dyad-INS during coop/comp > 0 | 66 | 33 | n.a. | 23.40 | 2.90 | TAL | Seed | 2000 | 0 | "high-pass" | 9 | **0.38** | **16.28** | **2.01** | **12.98** | **0.00** | **0.49** | **3.33** | **0.00** | **0.00** |
| **Xie et al., 2020** | 10.1073/pnas.1917407117 | Xie 2020 | hyper (3) | coop/comp | draw picture & guess drawing | no | dyad-INS during collaborative drawing > random-INS | 36 | 12 | 0.44 | 27.44 | 4.98 | MNI | wb-Atlas | 2000 | 0 | .008-.09 | 42 | **5.00** | **0.01** | **9.39** | **0.09** | **0.00** | **4.58** | **0.00** | **0.00** | **0.00** |
| **Yoshioka et al., 2021** | 10.1093/scan/nsab082 | Yoshioka 2021 | hyper | joint attention | direct attention to place or object | A/V | dyad-INS (task-dependent and independent) > random-INS | 44 | 22 | 0.55 | 21.27 | 2.38 | MNI | wb-VW | 2500 | 0 | >.008 | 17 | **3.48** | **14.84** | **12.21** | **11.24** | **34.40** | **1.85** | **0.00** | **3.63** | **0.00** |
| ***Summary (weighted)*** | | | | | | | | ***740*** | ***423*** | ***0.42*** | ***24.79*** | ***3.28*** | | | | | | ***297*** | ***99.15*** | ***99.44*** | ***99.40*** | ***99.58*** | ***99.64*** | ***79.94*** | ***4.35*** | ***9.83*** | ***15.23*** |

**Table S1: fMRI experiments included for meta-analysis**

See the separate supplementary file for the original table in Excel format. Data from 14 hyperscanning fMRI publications (*37–51*), 8 pseudohyperscanning fMRI publications (*52–59*) were included in the analyses. *Publications* refers to the single publications (n = 22 publications with n = 26 experiments) that were included in the meta-analyses. *Experiment ID* refers to the aggregated experiments when considering reanalyses of existing data (n = 22 experiments). Note that the experiment "Wang 2021" corresponding to Wang et al. (2022) (*49*) was included at the publication's preprint stage and updated later. The results did not differ between pre- and postprint

versions. *A/V* indicates whether participants in the experiments had contact per audio and/or video during fMRI scanning (live transmission in the case of hyperscanning experiments). *Method* refers to the general type of analysis method: Seed: seed-to-voxel (whole-brain), foci within the “target-brain” were included. Wb-ICA: ICA on whole-brain level, foci were peak coordiantes of reported independent components. ts-ICA: only independent components sensitive to the applied task were analyzed. wb-VW: voxelwise analysis on whole-brain level. ts-VW: voxelwise analysis on voxels sensitive to the applied task. wb-Atlas: a whole-brain atlas was applied, foci correspond to center-of-masses of reported parcels. *Lag* refers to whether time-series of interacting subjects were analyzed with time lag. If possible, we restricted the included foci to those from “zero lag” analyses. *Band* indicates whether a band pass or high pass filter was applied. *Contributions* refers to the relative contribution of each study to each cluster in the main fMRI ALE using the conservative voxellevel threshold ($p < .001$) or the lenient threshold ($p < .01$), and the combined fMRI and fNIRS ALE using the conservative thresholding. The ALE method incorporates nonlinear procedures causing the percentages to not add up to 100%.
Abbreviations: ALE = activation likelihood estimation, hyper(3) = hyperscanning (in one case between 3 subjects), pseudo = pseudo-hyperscanning, A = audio, V = video, INS = interpersonal neural synchronization, JA = join attention, JAct = joint action, y = years, MNI = montral neurological institute space, TAL = talairach space, wb = whole-brain, ts = task-specific, ICA = independent component analysis, VW = voxel-wise, TR = repitition time, rTPJ = right temporoparietal junction, rSTG = right superior temporal gyrus, rIns = right insula, r/lSFG = right/left superior frontal gyrus, lMFG = left medial frontal gyrus.

| Publication | DOI | Experiment ID | Type | Area | Task | Setting | Contrast | Subjects | Dyads | Female | Age | | Device | Length | Rate | Hb | Band | Coverage | Array | Source | Channels | | Contributions fMRI+fNIRS (p < .001) | | | |
|---|---|---|---|---|---|---|---|---|---|---|---|---|---|---|---|---|---|---|---|---|---|---|---|---|---|---|
| | | | | | | | | [N] | [N] | proportion | mean [y] | SD [y] | | [nm] | [Hz] | | | | | | all [N] | INS [N] | rTPJ [%] | lSFG [%] | rSFG [%] | rMFG [%] |
| **Balconi et al., 2017** | 10.1371/journal.pone.0187652 | Balconi 2017 | hyper | cooperation | synchronize sustained attention, pos feedback | side-by-side | ISC post positive feedback > ISC pre positive feedback | 26 | 13 | 0.50 | 24.08 | 1.78 | NIRScout | 760, 850 | 6.25 | HbO | .01-.03 | frontRL | 2x 2x2 | AV(3) | 8 | 2 | **0.00** | **0.00** | **0.00** | **0.00** |
| **Balconi et al., 2018** | 10.1016/j.bandc.2018.02.009 | Balconi 2018 | hyper | cooperation | synchronize sustained attention, neg feedback | side-by-side, screen | ISC pre negative feedback/control > ISC post negatove feedback | 26 | 13 | n.a. | 25.89 | 1.21 | NIRScout | 760, 850 | 6.25 | HbO | .01-.03 | frontRL | 2x 2x2 | AV(3) | 8 | 2 | **0.00** | **0.00** | **0.00** | **0.00** |
| **Balconi et al., 2019** | 10.1007/s12144-019-00247-4 | Balconi 2019 | hyper | cooperation | synchronize selective attention after gift exchange | side-by-side, screen | ISC after material gift </> ISC after experiential gift | 32 | 16 | 1.00 | 22.59 | 1.83 | NIRScout | 760, 850 | 6.25 | HbO | .01-.03 | frontRL | 2x 2x2 | AV(3) | 8 | 4 | **0.00** | **0.00** | **0.02** | **0.63** |
| **Cañigueral et al., 2021** | 10.1016/j.neuroimage.2020.117572 | Canigueral 2021 | hyper | communication | answering statement about oneself, shared vs. not shared | face-to-face, screen | prediction of partner's ISC from rTPJ during shared > private conditions | 30 | 15 | 0.73 | 28.20 | 7.33 | LABNIRS | 780, 805, 830 | 27 | HbR | whole | fronttempparRL | custom | Dig(voxel) | 58 | 3 | **0.00** | **0.00** | **0.00** | **0.00** |
| **Chen et al., 2020** | 10.1002/hbm.25173 | Chen 2020 1 | hyper | deception | spontaneous sender–receiver deception task, female dyads | face-to-face, screen | female dyad ISC during deception > rest | 44 | 22 | 1.00 | 21.30 | 2.50 | ETG-7100 | n.a. | 10 | HbO | .02-.07 | frontRL, tempR | 4x4, 4x4 | VR, AV(1) | 48 | 1 | **0.00** | **0.00** | **0.27** | **0.00** |
| | | Chen 2020 2 | hyper | deception | spontaneous sender–receiver deception task, male dyads | face-to-face, screen | male dyad ISC during deception > rest | 38 | 19 | 0.00 | 21.30 | 2.50 | ETG-7100 | n.a. | 10 | HbO | .02-.07 | frontRL, tempR | 4x4, 4x4 | VR, AV(1) | 48 | 3 | **0.00** | **0.00** | **0.00** | **0.00** |
| **Cheng et al., 2015** | 10.1002/hbm.22754 | Cheng 2015 | hyper | coop/comp | synchronize button press | side-by-side | ISC synchronize > press faster/rest in all participants | 90 | 45 | 0.51 | 21.96 | 2.15 | ETG-4000 | n.a. | 10 | HbO | .08-.31 | frontRL | 3x5 | VR | 22 | 2 | **0.00** | **6.09** | **7.97** | **0.00** |
| **Cheng et al., 2019** | 10.3389/fnins.2019.01071 | Cheng 2019 | hyper | cooperation | joint drawing task (joint controlling of brush) | face-to-face, wall | ISC during interpersonal coordination > rest | 62 | 31 | 0.76 | 21.39 | 2.36 | ETG-7100 | n.a. | 10 | HBO | .08-.31 | frontRL | 2x5 | VR | 13 | 7 | **0.00** | **15.23** | **0.01** | **0.00** |
| **Cheng et al., 2021** | 10.1016/j.neuroimage.2021.118777 | Cheng 2021 | hyper | cooperation | repeated trust game with subject's "social status" pre-determined | face-to-face, wall | ISC during trust interaction > rest | 198 | 99 | 1.00 | 21.05 | 2.47 | ETG-7100 | n.a. | 10 | HbO | .01-.05 | frontRL, tempparR | 3x5, 4x4 | VR, Wang 2019 | 46 | 1 | **0.00** | **0.00** | **0.00** | **0.00** |
| **Cui et al., 2012** | 10.1016/j.neuroimage.2011.09.003 | Cui 2012 | hyper | coop/comp | synchronize button press | side-by-side | ISC synchronize > press faster/rest | 22 | 11 | 0.55 | 26.00 | 6.00 | ETG-4000 | n.a. | 10 | HbO | .08-.3 | frontRL | 3x5 | VR | 22 | 1 | **0.00** | **0.00** | **7.97** | **0.00** |
| **Duan et al., 2020** | 10.1007/s12144-020-01093-5 | Duan 2020 1 | hyper | cooperation | realistic presented problem, lovers dyads | face-to-face, screen | ISC during problem solving in lovers > rest | 40 | 20 | 0.50 | 20.30 | 0.84 | LABNIRS | n.a. | 10 | HbO | .08-.31 | frontRL, tempR | 3x3, 3x2 | Wang 2019 | 19 | 2 | **0.00** | **7.54** | **0.00** | **0.00** |
| | | Duan 2020 2 | hyper | cooperation | realistic presented problem, strangers dyads | face-to-face, screen | ISC during problem solving in strangers > rest | 44 | 22 | 0.50 | 20.30 | 0.84 | LABNIRS | n.a. | 10 | HbO | .08-.31 | frontRL, tempR | 3x3, 3x2 | Wang 2019 | 19 | 0 | **n.a.** | **n.a.** | **n.a.** | **n.a.** |
| **Feng et al., 2020** | 10.1093/scan/nsaa017 | Feng 2020 | hyper | cooperation | simultaneous button pressing after cue | face-to-face, screen | ISC during simultaneous button press > rest | 60 | 30 | n.a. | 20.60 | 1.51 | ETG-4000 | 695, 830 | 10 | HbO | .08-.3 | frontRL | 3x5 | VR | 22 | 1 | **0.00** | **0.00** | **0.03** | **0.00** |
| **Fronda & Balconi, 2020** | 10.1002/brb3.1663 | Fronda & Balconi 2020 | hyper | communication | Repeat cued social vs. affective vs. informative gestures | side-by-side | ISC during affective > social/informative gestures | 34 | 17 | 0.82 | 26.89 | 0.03 | NIRScout | 760, 850 | 6.25 | HbO | .01-.03 | frontRL | 2x 2x2 | AV(3) | 8 | 1 | **0.00** | **0.00** | **0.00** | **0.00** |
| **Hou et al., 2020** | 10.1016/j.neuroimage.2020.116655 | Hou 2020 | pseudo | music | watching/listening video of violinist | video | watching/listening to violinist > rest | 17 | 16 | 0.94 | 20.35 | 1.92 | ETG-7100 | 695, 830 | 10 | HbO | .3-.7 | fronttempparL, fronttempparR | 3x5, 3x5 | VR | 44 | 4 | **0.00** | **0.00** | **0.00** | **0.00** |
| **Hu et al. , 2017** | 10.1093/scan/nsx118 | Hu 2017 | hyper | cooperation | synchronize button press | face-to-face, wall | ISC synchronize > rest | 70 | 35 | 1.00 | n.a. | n.a. | ETG-7100 | n.a. | 10 | HbO | .02-.08 | frontRL | 3x5 | VR | 22 | 1 | **0.00** | **0.01** | **0.00** | **0.00** |
| **Koide & Shimada, 2018** | 10.1111/jpr.12202 | Koide & Shimada 2018 | hyper | communication | cheering during rock-paper-scissors | side by side | ISC player-observer while playing > control | 64 | 32 | 0.00 | 21.30 | 1.60 | ONM-3000 | n.a. | 10 | HbO | n.a. | frontparL | 4x4 | AV(1) | 24 | 2 | **0.00** | **0.00** | **0.00** | **0.00** |
| **Li et al., 2020** | 10.3389/fnhum.2020.00169 | Li 2020 1 | hyper | cooperation | joint drawing task | face-to-face, screen | ISC during cooperation > rest | 24 | 12 | 0.00 | 19.95 | 1.43 | ETG-7100 | 695, 830 | 10 | HbO | .08-.17 | frontRL | 3x5 | VR | 22 | 0 | **n.a.** | **n.a.** | **n.a.** | **n.a.** |
| | | Li 2020 2 | hyper | cooperation | joint drawing task | face-to-face, screen | ISC during cooperation > rest | 24 | 12 | 0.00 | 19.70 | 1.87 | ETG-7100 | 695, 830 | 10 | HbO | .08-.17 | frontRL | 3x5 | VR | 22 | 0 | **n.a.** | **n.a.** | **n.a.** | **n.a.** |
| **Li Ya. et al., 2021** | 10.1093/scan/nsab114 | Li Ya 2021 | hyper | coop/comp | watch emotional or neutral movie; joint button press task | face-to-face, wall | ISC during cooperation after emotional movie > random-ISC | 62 | 31 | 0.61 | 21.96 | 2.64 | LABNIRS | 780, 805, 830 | 24 | HBO | .08-.31 | frontRL | 3x5 | n.a. | 22 | 1 | **0.00** | **0.00** | **0.00** | **0.00** |
| **Li Yu. et al., 2021** | 10.1007/s11682-020-00361-z | Li Yu 2021 | hyper | coop/comp | jenga game, cooperation vs. competition vs. independent | face-to-face | ISC during cooperation > independence | 26 | 13 | 0.00 | 21.14 | 2.01 | FOIRE-3000/16 | 780, 805, 830 | 4 | HbO | .04-.08 | frontRL | 3x5 | Dig | 22 | 1 | **0.00** | **0.02** | **0.00** | **0.00** |
| **Li Z. et al., 2021** | 10.1093/cercor/bhab118 | Li Z 2021 | pseudo | communication | listen to narrative stories, different noise levels | audio | speaker-listener ISC modulated by noise level | 22 | 16 | 0.50 | n.a. | n.a. | NirScan | 785, 808, 850 | 12 | HbO | .01-.03 | frontRL, tempparL, tempparR | custom, 2x4, 2x4 | n.a. | 36 | 20 | **2.60** | **5.78** | **9.53** | **0.00** |
| **Liu N. et al., 2016** | 10.3389/fnhum.2016.00082 | Liu N 2016 | hyper | coop/comp | jenga game, cooperation vs. competition vs. independent | face-to-face | ISC during coop/comp > rest | 18 | 9 | 0.50 | 21.10 | 1.70 | ETG-4000 | n.a. | 10 | HbO | .04-.08 | frontRL, tempparR | 3x3, 3x3 | VR(voxel) | 19 | 3 | **0.00** | **0.06** | **3.65** | **0.00** |
| **Liu T. et al., 2017** | 10.1038/s41598-017-09226-w | Liu T 2017 | hyper | coop/comp | pattern building game | side-by-side | ISC during coop/comp > random ISC | 44 | 22 | 0.00 | 19.00 | 1.40 | LABNIRS | n.a. | 37 | HbO | n.a. | frontparL, frontparR | 4x4, 4x4 | AV(1) | 48 | 6 | **0.00** | **0.00** | **0.01** | **17.28** |
| **Liu Y. et al, 2017** | 10.1038/srep43293 | Liu Y 2017 | pseudo | communication | listen to real-life story, english (native) or turkish | audio | speaker-listener ISC > 0 | 18 | 15 | 0.44 | n.a. | n.a. | fNIR-1100, ETG-4000 | n.a. | 2 | HbO | .01-.05 | frontRL, parL, parR | custom, 3x3, 3x3 | VR | 40 | 15 | **6.27** | **0.00** | **0.00** | **0.00** |
| **Liu J. et al., 2019** | 10.1016/j.neuroimage.2019.03.004 | Liu J 2019 | hyper | learning | teaching face-to-face or pc-mediated | face-to-face vs. back-to-back | ISC during face-to-face teaching > rest | 84 | 42 | 0.76 | 21.00 | 2.30 | ETG-7100 | n.a. | 10 | HbO | .02-1 | frontRL, tempparR | 3x5, 4x4 | VR, Lu 2020b | 46 | 3 | **0.00** | **0.00** | **0.00** | **0.00** |
| **Liu W. et al., 2019** | 10.1016/j.neuroimage.2019.05.035 | Liu W 2019 | hyper | communication | complete sentence via picture, partner rates correspondence | face-to-face vs. back-to-back | ISC during same > different syntactic structures, eye-contact > no eye contact | 180 | 90 | 0.56 | 20.00 | 1.60 | ETG-4000 | 695, 830 | 10 | HbO | .02-.05 | frontparR, frontparL | 2x4, 2x4 | AV(1) | 20 | 2 | **0.00** | **8.03** | **0.00** | **0.00** |
| **Long et al., 2021a/ 2021b** | 10.1093/cercor/bhab413 10.1093/cercor/bhaa316 | Long 2021a + Long 2021b | hyper | communication/ touch | communicate freely or hold each other's hands | face-to-face | ISC interaction of communication mode x topic/ during touch > verbal communication/ in couples > friends | 88 | 44 | 0.50 | 21.27 | 2.04 | LABNIRS | 780, 805, 830 | 55.6 | HbO | .04-.08 | frontparL, frontparL | 2x5, 2x5 | MRI(1) | 26 | 3 | **0.00** | **0.00** | **0.00** | **0.00** |
| **Lu et al., 2019a** | 10.1016/j.neuropsychologia.2019.01.004 | Lu 2019a 1 | hyper | communication | "brainstorming" - neg/pos/no feedback | face-to-face, in triangle | ISC during pos feedback after brainstorming > rest | 40 | 20 | n.a. | 20.72 | 2.47 | ETG-7100 | 696, 830 | 10 | HbO | .018-.048 | frontRL | 3x5 | Lu 2020b | 22 | 12 | **0.00** | **5.24** | **11.32** | **15.63** |
| | | Lu 2019a 2 | hyper | communication | "brainstorming" - neg/pos/no feedback | face-to-face, in triangle | ISC during neg feedback after brainstorming > rest | 38 | 19 | n.a. | 20.72 | 2.47 | ETG-7100 | 696, 830 | 10 | HbO | .018-.048 | frontRL | 3x5 | Lu 2020b | 22 | 7 | **0.00** | **0.00** | **6.04** | **15.63** |
| **Lu et al., 2019a 3/ Lu & Hao, 2019** | 10.1016/j.neuropsychologia.2019.01.004 10.1093/scan/nsz012 | Lu 2019a 3 + Lu & Hao 2019 | hyper | communication | "brainstorming" - neg/pos/no feedback | face-to-face, in triangle | ISC during/ after brainstorming > rest | 38 | 19 | n.a. | 20.72 | 2.47 | ETG-7100 | 696, 830 | 10 | HbO | .018-.048 | frontRL | 3x5 | Lu 2020b | 44 | 2 | **0.00** | **0.00** | **6.07** | **0.05** |
| **Lu et al., 2019b** | 10.1093/cercor/bhy215 | Lu 2019b 1 | hyper | communication | "brainstorming": find alternative uses for everyday objects (AUT) | face-to-face | ISC during cooperative AUT > rest | 50 | 25 | n.a. | 21.00 | 1.52 | ETG-7100 | 695, 830 | 10 | HbO | .042-.045 | frontRL, tempparR | 3x5, 4x4 | Lu 2020b | 46 | 4 | **0.00** | **0.00** | **6.07** | **0.05** |
| | | Lu 2019b 2 | hyper | communication | "brainstorming": find typical uses for everyday objects (OTC) | face-to-face | ISC during cooperative OCT > rest | 52 | 26 | n.a. | 21.00 | 1.52 | ETG-7100 | 695, 830 | 10 | HbO | .042-.045 | frontRL, tempparR | 3x5, 4x4 | Lu 2020b | 46 | 0 | **n.a.** | **n.a.** | **n.a.** | **n.a.** |
| **Lu et al., 2020a** | 10.1007/s00221-020-05799-7 | Lu 2020a | hyper | communication | "brainstorming": AUT vs. OCT | face-to-face | ISC during brainstorming > male & mixed dyads | 132 | 66 | 0.56 | 21.23 | 2.91 | ETG-7100 | 695, 830 | 10 | HbO | .08-.14 | frontRL, tempparR | 3x5, 4x4 | Lu 2020b | 46 | 1 | **0.00** | **0.00** | **0.00** | **0.00** |
| **Lu et al., 2020b** | 10.1016/j.neuroimage.2020.117025 | Lu 2020b | hyper | communication | generating creative uses for everyday objects | face-to-face, wall | ISC during turn taking > normal/virtual communication | 54 | 27 | 0.81 | 20.52 | 2.22 | ETG-7100 | 695, 830 | 10 | HbO | .34-.48 | frontRL, tempparR | 3x5, 4x4 | NA | 46 | 1 | **0.00** | **0.00** | **0.00** | **0.00** |
| **Nozawa et al., 2016** | 10.1016/j.neuroimage.2016.03.059 | Nozawa 2016 | hyper | cooperation | word-chain game | face-to-face, around table | (dyad-ISC during word-chain game > control) > random-ISC | 48 | 12 | 0.42 | 21.90 | NA | self-build | 810 | 10 | HbT | .01-1 | front | 1x3 | AV(3) | 1 | 1 | **0.00** | **0.01** | **0.00** | **0.00** |
| **Nozawa et al., 2019** | 10.1038/s41598-019-49257-z | Nozawa 2019 | hyper | learning | 1. moving arm in sync/indep.; 2. teaching/learning unknown words | face-to-face | ISC during teaching/learning after synchronized > independent movement | 64 | 32 | 0.28 | 21.50 | 1.50 | HOT-1000 | 810 | NA | HbT | NA-.2 | frontR, frontL | 1x3, 1x3 | AV(3) | 2 | 1 | **0.00** | **4.41** | **0.00** | **0.00** |
| **Pan et al., 2017** | 10.1002/hbm.23421 | Pan 2017 1 | hyper | cooperation | synchronize button press; "lovers" | side-by-side, wall | dyad-ISC during joint button press > rest | 34 | 17 | 0.50 | 21.07 | 1.85 | ETG-4000 | 695, 830 | 10 | HbO | .08-.31 | frontparR | 3x5 | VR | 22 | 1 | **0.00** | **0.00** | **0.00** | **0.00** |
| | | Pan 2017 2 | hyper | cooperation | synchronize button press; "friends" | side-by-side, wall | dyad-ISC during joint button press > rest | 32 | 16 | 0.50 | 21.07 | 1.85 | ETG-4000 | 695, 830 | 10 | HbO | .08-.31 | frontparR | 3x5 | VR | 22 | 0 | **n.a.** | **n.a.** | **n.a.** | **n.a.** |
| | | Pan 2017 3 | hyper | cooperation | synchronize button press; "strangers" | side-by-side, wall | dyad-ISC during joint button press > rest | 32 | 16 | 0.50 | 21.07 | 1.85 | ETG-4000 | 695, 830 | 10 | HbO | .08-.31 | frontparR | 3x5 | VR | 22 | 0 | **n.a.** | **n.a.** | **n.a.** | **n.a.** |
| **Pan et al., 2018** | 10.1016/j.neuroimage.2018.08.005 | Pan 2018 1 | hyper | learning | learn to sing a song - "learn phrase by phrase" | face-to-face | dyad-ISC during interactive learning > rest | 13 | 12 | 1.00 | 20.69 | 2.15 | ETG-7100 | 695, 830 | 10 | HbO | .07-.15 | frontparR, frontparL | 3x5, 3x5 | VR | 44 | 4 | **0.08** | **0.00** | **0.00** | **0.00** |
| | | Pan 2018 2 | hyper | learning | learn to sing a song - "whole song at once" | face-to-face | dyad-ISC during interactive learning > rest | 13 | 12 | 1.00 | 20.69 | 2.15 | ETG-7100 | 695, 830 | 10 | HbO | .07-.15 | frontparR, frontparL | 3x5, 3x5 | VR | 44 | 4 | **0.08** | **0.00** | **0.00** | **0.00** |
| **Pan et al., 2020** | 10.1016/j.neuroimage.2020.116657 | Pan 2020 | hyper | learning | dynamic conceptual learning | face-to-face | interactive learning > rest | 48 | 24 | 1.00 | 21.46 | 2.75 | ETG-7100 | 695, 830 | 10 | HbO | .17-.27,.45-.57 | frontRL, tempparL | 3x5, 4x4 | VR | 46 | 22 | **0.00** | **15.23** | **0.04** | **17.28** |
| **Pan et al., 2021** | 10.1016/j.bcp.2020.114111 | Pan 2021 | hyper | learning | teaching of numerical reasoning | side-by-side | ISC during teaching > rest | 18 | 16 | 1.00 | 22.72 | 1.99 | BrainSight | 685, 830 | 10 | HbO | .16-.19 | fronttempparR, fronttempparL | custom | Dig | 26 | 13 | **3.93** | **0.00** | **0.00** | **0.00** |
| **Pinti et al., 2021** | 10.1093/scan/nsaa086 | Pinti 2021 | hyper | deception | card game, telling truth/lying, followed by guessing | face-to-face | predict guesser's from informer's activity | 46 | 23 | n.a. | 26.40 | 10.10 | LIGHTNIRS | 780, 805, 830 | 13.33 | HbT | .01-.4 | frontRL | 2x8 | Dig(1) | 22 | 2 | **0.00** | **4.28** | **1.17** | **0.00** |
| **Quiñones-Camacho et al., 2021** | 10.1002/aur.2513 | Quinones-Camacho 2021 | hyper | communication | free conversation with prompted topics, moderated by experimenter | face-to-face | HC-dyad-ISC during communication > random-ISC | 26 | 23 | n.a. | NA | NA | NIRScout | 760, 850 | 15.63 | HbO | NA | tempR, frontR, frontR, frontR | custom | AV(3) | 8 | 4 | **0.00** | **0.00** | **0.00** | **0.00** |
| **Sun et al., 2020** | 10.1016/j.bandc.2019.105513 | Sun 2020 1 | hyper | cooperation | cooperative solving of math equations, novice-student | face-to-face, wall | ISC during cooperation > rest | 32 | 16 | 0.31 | 22.98 | 3.18 | ETG-4000 | NA | 10 | HbO | .04-.16 | frontRL | 3x5 | VR | 22 | 0 | **n.a.** | **n.a.** | **n.a.** | **n.a.** |
| | | Sun 2020 2 | hyper | cooperation | cooperative solving of math equations, expert-student | face-to-face, wall | ISC during cooperation > rest | 36 | 18 | 0.28 | 29.08 | 2.99 | ETG-4000 | NA | 10 | HbO | .04-.16 | frontRL | 3x5 | VR | 22 | 1 | **0.00** | **0.05** | **0.00** | **0.00** |
| **Sun et al., 2021** | 10.1016/j.bandc.2021.105803 | Sun 2021 | hyper | cooperation | drawing task, control pointer alone or jointly | face-to-face, wall | ISC interaction of student-student vs teacher-student x task condition | 86 | 43 | 0.81 | 25.71 | 3.18 | ETG-4000 | NA | 10 | HbO | .02-.08 | frontRL | 3x5 | VR | 22 | 1 | **0.00** | **0.06** | **0.00** | **0.00** |
| **Tang et al., 2016** | 10.1093/scan/nsv092 | Tang 2016 1 | hyper | decision making | modified ultimatum game - "face to face" | face-to-face, screen | dyad ISC during ultimatum game > rest | 102 | 51 | 0.57 | 22.68 | 2.07 | ETG-4000 | NA | 10 | HbO | .02-.08 | frontR, tempparR | 3x3, 2x3 | AV(1) | 19 | 2 | **4.42** | **0.00** | **0.00** | **0.00** |
| | | Tang 2016 2 | hyper | decision making | modified ultimatum game - "face blocked" | face-to-face, screen | dyad ISC during ultimatum game > rest | 92 | 46 | 0.50 | 22.20 | 2.18 | ETG-4000 | NA | 10 | HbO | .02-.08 | frontR, tempparR | 3x3, 2x3 | AV(1) | 19 | 1 | **1.55** | **0.00** | **0.00** | **0.00** |
| **Wang et al., 2019** | 10.1002/hbm.24592 | Wang 2019 | hyper | coop/comp | synchronize button press (physical pain feedback) | face-to-face, screen | ISC during cooperation with pain feedback > rest | 32 | 16 | 1.00 | 20.90 | 2.10 | ETG-7100 | NA | 10 | HbO | .08-.16 | frontRL, tempparR | 3x5, 4x4 | VR | 46 | 8 | **0.00** | **0.00** | **3.65** | **17.28** |
| **Xue et al., 2018** | 10.1016/j.neuroimage.2018.02.007 | Xue 2018 1 | hyper | cooperation | brainstorming on problem; high-low dyads | face-to-face, in circle | ISC in high-low creativity dyads during brainstorming > rest | 20 | 10 | n.a. | 21.00 | 1.40 | ETG-7100 | 695, 830 | 10 | HbO | .02-.08 | frontRL, tempparR | 3x5, 4x4 | VR, Wang 2019 | 46 | 1 | **0.00** | **7.54** | **0.00** | **0.00** |
| | | Xue 2018 2 | hyper | cooperation | brainstorming on problem; high-high dyads | face-to-face, in circle | ISC in high-high creativity dyads during brainstorming > rest | 20 | 10 | 0.00 | 21.00 | 1.40 | ETG-7100 | 695, 830 | 10 | HbO | .02-.08 | frontRL, tempparR | 3x5, 4x4 | VR, Wang 2019 | 46 | 0 | **n.a.** | **n.a.** | **n.a.** | **n.a.** |
| | | Xue 2018 3 | hyper | cooperation | brainstorming on problem; low-low dyads | face-to-face, in circle | ISC in low-low creativity dyads during brainstorming > rest | 20 | 10 | 0.00 | 21.00 | 1.40 | ETG-7100 | 695, 830 | 10 | HbO | .02-.08 | frontRL, tempparR | 3x5, 4x4 | VR, Wang 2019 | 46 | 4 | **0.00** | **7.54** | **7.97** | **0.00** |
| **Yang et al., 2020** | 10.1038/s41593-020-0630-x | Yang 2020 | hyper(6) | cooperation | incentive comp. game, "attack" vs. "defend", "bonding vs no-bonding" | face-to-face, in circle | within-group ISC interaction of bonding/no-bonding x attack/defend | 516 | 86 | 0.56 | 22.01 | 1.29 | LABNIRS | 780, 805, 830 | 47.62 | HbO | .026-.167 | frontR, tempR | 2x3, 3x2 | MRI(6) | 14 | 4 | **0.01** | **0.00** | **1.07** | **0.02** |
| **Zhang M. et al., 2017a/ 2017b** | 10.1038/s41598-017-06764-1 10.1016/j.bandc.2017.08.008 | Zhang M 2017a + 2017b | hyper | deception, decision making | poker-like card game | face-to-face | ISC during "deception" > "honesty", "risk-seeking" > "risk-averting" | 36 | 18 | 0.50 | 22.30 | 2.40 | LABNIRS | 780, 805, 830 | 57 | HbO | .05-.2 | frontRL, tempL | 3x3, 2x3 | Dig | 27 | 6 | **0.00** | **0.00** | **5.68** | **0.00** |
| **Zhang M. et al., 2020/ 2021a** | 10.3389/fpsyg.2020.542093 10.3389/fnhum.2021.702959 | Zhang M 2020 + 2021a | hyper | cooperation | prisoner's dilemma with high/low incentive reward | side-by-side, screen | ISC during cooperation > rest, high > low reward | 62 | 31 | 0.52 | 22.30 | 2.40 | LABNIRS | NA | 42 | HbO | .01-.2 | frontR, frontRL, frontL | 2x2, 2x3, 2x2 | Dig | 30 | 4 | **0.00** | **0.00** | **5.69** | **0.00** |
| **Zhang M. et al., 2021b** | 10.1016/j.paid.2020.110315 | Zhang M 2021b | hyper | decision making | prisoner's dilemma, dyads act as group & play against experimenter | side-by-side, in front of exper. | ISC during group > individual decision making | 54 | 27 | 0.48 | 20.70 | 3.10 | LABNIRS | NA | NA | HbO | .01-.2 | frontR, frontRL, frontL | 2x2, 2x3, 2x2 | Dig | 15 | 1 | **0.00** | **0.00** | **0.00** | **0.00** |
| **Zhang R. et al., 2021** | 10.1016/j.bandc.2021.105738 | Zhang R 2021 1 | hyper | coop/comp | responding simultaneously to visual cue with vs. without stress | face-to-face, screen | ISC during cooperation with stress > random ISC | 38 | 19 | 1.00 | 20.90 | 2.30 | ETG-7100 | 695, 830 | 10 | HbO | .08-.17 | frontRL, tempparR | 3x5, 4x4 | Wang 2019 | 46 | 4 | **0.00** | **0.01** | **0.01** | **0.00** |
| | | Zhang R 2021 2 | hyper | coop/comp | respondingsimultaneously to visual cue with vs. without stress | face-to-face, screen | ISC during cooperation without stress > random ISC | 42 | 21 | 1.00 | 20.90 | 2.30 | ETG-7100 | 695, 830 | 10 | HbO | .08-.17 | frontRL, tempparR | 3x5, 4x4 | Wang 2019 | 46 | 1 | **0.00** | **0.00** | **0.00** | **0.00** |
| **Zhang Y. et al., 2020** | 10.1523/ENEURO.0236-20.2020 | Zhang Y 2020 1 | hyper | communication | psychological counseling, expert counselor (manual) | face-to-face | ISC during psychological counseling by expert > rest | 17 | 14 | 1.00 | 23.50 | NA | ETG-7100 | 695, 830 | 10 | HbO | .024-.045 | tempparR | 4x4 | Wang 2019 | 24 | 6 | **0.06** | **0.00** | **0.00** | **0.00** |
| | | Zhang Y 2020 2 | hyper | communication | psychological counseling, novice counselor (manual) | face-to-face | ISC during psychological counseling by novice > rest | 21 | 16 | 1.00 | 21.98 | NA | ETG-7100 | 695, 830 | 10 | HbO | .024-.045 | tempparR | 4x4 | Wang 2019 | 24 | 1 | **0.00** | **0.00** | **0.00** | **0.00** |
| **Zhao et al., 2017** | 10.1117/1.JBO.22.2.027004 | Zhao 2017 | hyper | joint action | joint tapping task | back-to-back | ISC during joint tapping > 0 | 48 | 24 | 0.50 | 22.77 | 2.19 | ETG-4000 | NA | NA | HbO | NA | frontRL | 2x4 | AV(1) | 10 | 2 | **0.00** | **0.00** | **0.22** | **0.00** |
| **Zhao et al., 2021** | 10.1037/npe0000138 | Zhao 2021 1 | hyper | decision making | Iowa gambling task after Trier social stress test | side-by-side, single screen | ISC during decision making after stress > rest | 42 | 21 | 1.00 | 21.35 | 2.40 | LABNIRS | 780, 805, 830 | 10 | HbO | .031-.125 | frontRL, tempparL | 3x3, 2x3 | Dig | 19 | 2 | **0.00** | **0.00** | **0.00** | **0.00** |
| | | Zhao 2021 2 | hyper | decision making | Iowa gambling task after control task | side-by-side, single screen | ISC during decision making without stress > rest | 42 | 21 | 1.00 | 21.35 | 2.40 | LABNIRS | 780, 805, 830 | 10 | HbO | .031-.125 | frontRL, tempparL | 3x3, 2x3 | Dig | 19 | 1 | **0.00** | **0.00** | **4.84** | **0.00** |
| **Zheng et al., 2018/ 2021 1** | 10.1002/hbm.24059 10.1037/edu0000707 | Zheng 2018 + 2020 1 | hyper | learning | teaching numerical reasoning, interact vs. no interact vs. recorded | side-by-side, screen | ISC during live > prerecorded video teaching, post > pre interaction | 64 | 60 | 0.50 | 23.13 | 2.31 | ETG-4000 | 686, 830 | 10 | HbO | .06-.07 | tempparR, frontparL | 2x4, 2x4 | MRI(1) | 40 | 3 | **0.02** | **0.00** | **0.00** | **0.00** |
| **Zhu et al., 2021** | 10.1037/edu0000707 | Zhu 2021 1 | hyper | learning | teaching/learning definitions, elaborated feedback from instructor | face-to-face | (dyad-ISC during elaborated feedback > rest) > random-ISC | 36 | 24 | 1.00 | 20.34 | 2.11 | ETG-7100 | 695, 830 | 10 | HbO | .017-.025 | frontRL, tempparL | 3x5, 4x4 | VR | 46 | 5 | **0.00** | **7.55** | **0.00** | **0.00** |
| | | Zhu 2021 2 | hyper | learning | teaching/learning definitions, simple feedback from instructor | face-to-face | (dyad-ISC during simple feedback > rest) > random-ISC | 36 | 24 | 1.00 | 20.28 | 2.16 | ETG-7100 | 695, 830 | 10 | HbO | .01-.7 | frontRL, tempparL | 3x5, 4x4 | VR | 46 | 0 | **n.a.** | **n.a.** | **n.a.** | **n.a.** |
| ***Summary (weighted)*** | | | | | | | | ***3721*** | ***1793*** | ***0.54*** | ***20.53*** | ***2.05*** | | | | | | | | | ***1987*** | ***228*** | ***19.04*** | ***94.70*** | ***89.29*** | ***83.85*** |

**Table S2: fNIRS experiments included for meta-analysis**

See the separate supplementary file for the original table in Excel format. Data from 54 hyperscanning fNIRS publications (*3*, *5*, *60–111*) and 3 pseudohyperscanning fNIRS publications (*112–114*) were included in the analyses. *Publications* refers to the single publications (n = 57) that were included in meta-analyses. *Experiment ID* refers to the aggregated experiments when considering separately analyzed subgroups and reanalyses of existing data (n = 69 experiments).

*Setting* indicates how participants were seated during the experiment. When subjects were seated side-by-side or face-to-face, they nevertheless often were separated by a portable wall or similar. *Length* refers to the wave-length of the light emitted by the fNIRS device. *Band* indicates how band-pass filtering was applied. *Coverage* refers to the general placement of fNIRS probe arrays on the head. Multiple entries indicate multiple separated probe arrays. *Array* refers to the desing of the probe array, “custom” indicates a non-square format. *Sources* lists the source of fNIRS coordinates: AV: AtlasViewer (*115*), Dig: 3D digitizer, VR: virtual registration (*2*), MRI: anatomical MRI-based registration, or from another publication. For “Dig” and “MRI”, numbers in brackets following the abbreviations indicate the number of subjects in the sample from which coordinates were estimated. For “AV”, numbers in brackets show reconstruction quality coded as: 3 = all optodes have assigned 10-10 positions, 2 = at least two 10-10 positions for each optode array, 1 = one 10-10 positions and information about the array alignment. *Contributions* refers to the relative contribution of each study to each cluster in the ALE analysis on the combined fMRI and fNIRS data. Of note, the ALE method incorporates non-linear procedures causing the percentages to not add up to 100%.

Abbreviations: ALE = activation likelihood estimation, hyper = hyperscanning, pseudo = pseudo-hyperscanning, INS = interpersonal neural synchronization, y = years, rTPJ = right temporoparietal junction, r/lSFG = right/left superior frontal gyrus, lMFG = left medial frontal gyrus.

| Target system | Target function | Atlases | | | | |
|---|---|---|---|---|---|---|
| | | Name | Tracer | Subjects (N) | Mean age (y) | Source |
| Serotonin | serotonin receptor 1a | 5HT1a | (11C)WAY100635 | 35 | 26.3 | Savli et al., 2012 (*118*) |
| | serotonin receptor 1b | 5HT1b | (11C)P943 | 65, 23 | 33.7, 28.7 | Gallezot et al., 2010; Savli et al., 2012 (*118*, *119*) |
| | serotonin receptor 2a | 5HT2a | (11C)Cimbi-36 | 29 | 22.6 | Beliveau et al., 2017 (*120*) |
| | serotonin receptor 4 | 5HT4 | (11C)SB207145 | 59 | 25.9 | Beliveau et al., 2017 (*120*) |
| | serotonin receptor 6 | 5HT6 | (11C)GSK215083 | 30 | 36.6 | Radhakrishnan et al., 2018 (*121*) |
| | serotonin transporter | 5HTT | (11C)DASB | 100, 18 | 25.1, 30.5 | Beliveau et al., 2017; Savli et al., 2012 (*118*, *120*) |
| Dopamine | dopamine synthesis | FDOPA | (18F)fluorodopa | 12 | n.a. | García-Gómez et al., 2018 (*122*) |
| | dopamine receptor 1 | D1 | (11C)SCH23390 | 13 | 33 | Kaller et al., 2017 (*123*) |
| | dopamine receptor 2 | D2 | (11C)FLB-457 | 37, 55 | 48.4, 32.5 | Sandiego et al., 2015; Smith et al., 2019 (*124*, *125*) |
| | dopamine transporter | DAT | (123I)FP-CIT | 174, 30 | 61, *n.a.* | Dukart et al., 2018; García-Gómez et al., 2013 (*126*, *127*) |
| Noradrenaline | noradrenaline transporter | NET | (11C)O-MRB | 77, 10 | 33.4, 33.3 | Hesse et al., 2017; Ding et al., 2010 (*128*, *129*) |
| GABA | GABA receptor A | GABAa | (11C)flumazenil | 6, 16 | *n.a.*, 26.6 | Dukart et al., 2018; Nørgaard et al., 2021 (*126*, *130*) |
| Glutamate | metabotropic receptor 5 | mGluR5 | (11C)ABP688 | 73, 22, 28 | 19.9, 67.9, 33.1 | DuBois et al., 2016; Hansen et al., 2022; Smart et al., 2019 (*131–133*) |
| | NMDA receptor | NMDA | (18F)GE-179 | 29 | 41 | Galovic et al., 2021 (*134*) |
| Acetylcholine | α4β2 nicotinic receptor | a4b2 | (F18)flubatine | 30 | 33.5 | Hillmer et al., 2016 (*135*) |
| | muscarinic receptor 1 | M1 | (11C)LSN3172176 | 24 | 40.5 | Naganawa et al., 2021 (*136*) |
| | vesicular Ach transporter | VachT | (18F)FEOBV | 4, 18, 5 | 37, 66.8, 68.3 | Aghourian et al., 2017; Bedard et al., 2019; Hansen et al., 2022 (*132*, *137*, *138*) |
| Endorphins | μ receptor | MU | (11C)carfentanil | 204, 39 | 32.3, *n.a.* | Hansen et al., 2022; Kantonen et al., 2020 (*132*, *139*) |
| Histamine | histamine receptor 3 | H3 | (11C)GSK189254 | 8 | 31.7 | Gallezot et al., 2017 (*140*) |
| Cannabinoid | cannabinoid receptor 1 | CB1 | (11C)OMAR | 77 | 30 | Normandin et al., 2015 (*141*) |
| Synaptic density | vesicle glycoprotein 2A | SV2a | (11C)UCB-J | 10 | 36 | Finnema et al., 2016 (*142*) |
| Oxytocin | oxytocin | OXT | mRNA expression | up to 6 | 24 – 57 | Shen et al., 2012; Markello et al., 2021 (*143*, *20*) |
| | oxytocin receptor | OXTR | | | | |
| | oxytocin release | CD38 | | | | |

**Table S3: Sources of neurotransmitter maps**

All nuclear imaging in vivo neurotransmitter atlases are drawn from JuSpace or neuromaps (*144*, *145*), mRNA expression atlases are derived from the Allen brain atlas (*143*) using the abagen toolbox with default settings (*20*). Sources listed in the table are the original sources of the atlases. For spatial correlation analyses, the atlases were parcellated, Z-standardized, and the weighted mean was calculated if multiple atlases using the same tracer were available.
Abbreviations: y = years.

| **Analysis** | **Dataset** | **Sources** | **N (categories)** | | **N (genes)** |
|---|---|---|---|---|---|
| | | | **all** | **included** | |
| **Neuronal cell types** | *PsychENCODE* cell markers (transcripts per kilobase counts) | Wang et al., 2018; Darmanis et al., 2015; Lake et al., 2016 (*146*, *147*, *155*) | 24 | 24 | 2 – 83 |
| **Developmental regional enrichment** | *BrainSpan* expression data (expression > 0.9$^{th}$ quantile, annotated to ≤ 20% of categories) | Miller et al., 2014; Grote et al., 2016 (*149*, *156*) | 80 | 80 | 5 – 224 |
| **Psychiatric disorders** | *DisGeNET* disease markers (manually curated dataset) | Piñero et al., 2020; Jiao et al., 2012 (*150*, *157*) | 332 | 150 | 5 – 697 |
| **Biological functions** | *Gene Ontology* biological processes (annotations propagated through the hierarchy, retrieved March 2022) | Ashburner et al., 2000; The Gene Ontology Consortium et al., 2021; Jiao et al., 2012 (*151*, *152*, *157*) | 15,039 | 6,947 | 5 – 200 |

**Table S4: Gene category enrichment datasets**

All datasets, except for *BrainSpan* data, were available as genetic "markers". Gene-wise expression values in the *BrainSpan* dataset were thresholded according to the above settings to identify markers for each of 80 categories. *DisGeNET* and *BrainSpan* categories were thresholded to contain at least 5 genes, *Gene Ontology* categories were thresholded to contain between 5 and 200 genes (*153*).
URLs: *PsychENCODE*: http://resource.psychencode.org/, *BrainSpan*: https://www.brainspan.org/, *ABAEnrichment*: https://bioconductor.org/packages/ABAEnrichment/, *DisGeNET*: https://www.disgenet.org/, *Gene Ontology*: http://geneontology.org/, *DAVID*: https://david.ncifcrf.gov/.

| Analysis | Cluster | X | Y | Z | Estimate | Volume | AAL regions |
|---|---|---|---|---|---|---|---|
| **ALE (p < .001)** | rTPJ | 62 | -48 | 16 | 0.019 | 2856 | 58.26% Temporal_Mid_R; 36.41% Temporal_Sup_R |
| | rSTG | 50 | -20 | -6 | 0.017 | 752 | 82.98% Temporal_Sup_R; 15.96% Temporal_Mid_R |
| **ALE (p < .01)** | rTPJ | 62 | -48 | 16 | 0.041 | 7856 | 47.45% Temporal_Mid_R; 25.25% Temporal_Sup_R; 12.02% SupraMarginal_R; 8.66% Temporal_Inf_R |
| | rSTG | 50 | -20 | -6 | 0.025 | 3368 | 88.84% Temporal_Sup_R; 8.55% Temporal_Mid_R |
| | rIns | 38 | 20 | -2 | 0.021 | 3296 | 52.43% Insula_R; 18.69% Frontal_Inf_Orb_2_R; 13.11% Frontal_Inf_Tri_R; 10.19% Frontal_Inf_Oper_R; 5.58% no_label |
| **ALE (no pseudo)** | rTPJ | 62 | -48 | 16 | 0.016 | 2152 | 63.94% Temporal_Mid_R; 32.71% Temporal_Sup_R |
| **ALE & fNIRS (all fNIRS)** | rTPJ | 62 | -48 | 16 | 0.019 | 3440 | 63.95% Temporal_Mid_R; 34.42% Temporal_Sup_R |
| | lSFG | -26 | 66 | 8 | 0.021 | 2248 | 64.41% Frontal_Sup_2_L; 20.28% no_label; 10.32% Frontal_Sup_Medial_L |
| | rSFG | 28 | 54 | 32 | 0.020 | 2040 | 47.84% Frontal_Sup_2_R; 23.92% Frontal_Sup_Medial_R; 23.14% Frontal_Mid_2_R; 5.10% no_label |
| | rMFG | 36 | 38 | 42 | 0.021 | 784 | 100.00% Frontal_Mid_2_R |
| **ALE & fNIRS (restricted fNIRS)** | rTPJ | 60 | -48 | 16 | 0.022 | 3368 | 63.42% Temporal_Mid_R; 34.92% Temporal_Sup_R |
| | lSFG | -26 | 66 | 8 | 0.021 | 2064 | 64.34% Frontal_Sup_2_L; 19.38% no_label; 10.85% Frontal_Sup_Medial_L |
| | rMFG/ rSFG | 28 | 54 | 32 | 0.021 | 824 | 51.46% Frontal_Mid_2_R; 42.72% Frontal_Sup_2_R; 5.83% no_label |
| | rMFG | 36 | 38 | 42 | 0.021 | 792 | 100.00% Frontal_Mid_2_R |
| **MACM** | rTPJ | 60 | -48 | 16 | 0.085 | 28576 | 39.39% Temporal_Mid_R; 18.20% Temporal_Sup_R; 11.62% SupraMarginal_R; 9.55% Temporal_Inf_R; 7.84% Fusiform_R; 6.63% Occipital_Inf_R |
| | lTPJ | -58 | -42 | 22 | 0.066 | 26280 | 35.43% Temporal_Mid_L; 17.08% SupraMarginal_L; 15.68% Temporal_Sup_L; 8.80% Fusiform_L; 5.02% Temporal_Inf_L |
| | lPFCIns | -32 | 20 | 2 | 0.068 | 22456 | 24.90% Frontal_Inf_Tri_L; 21.66% Insula_L; 18.28% Precentral_L; 15.57% Frontal_Inf_Oper_L; 12.22% Frontal_Inf_Orb_2_L |
| | rPFCIns | 36 | 20 | -2 | 0.069 | 19760 | 22.43% Frontal_Inf_Oper_R; 19.64% Insula_R; 19.51% Precentral_R; 18.70% Frontal_Inf_Tri_R; 6.80% Frontal_Inf_Orb_2_R; 5.71% no_label |
| | SMA | -4 | 12 | 48 | 0.070 | 11416 | 32.94% Supp_Motor_Area_L; 24.04% Supp_Motor_Area_R; 20.88% Cingulate_Mid_R; 9.39% Cingulate_Mid_L; 8.20% Frontal_Sup_Medial_L |
| | lTh | -10 | -18 | 4 | 0.066 | 5672 | 54.72% Thalamus_L; 16.50% no_label; 14.10% Hippocampus_L; 10.86% Amygdala_L |
| | lIPL | -34 | -50 | 46 | 0.061 | 5448 | 64.32% Parietal_Inf_L; 27.02% Parietal_Sup_L; 5.87% Precuneus_L |
| | rTh | 10 | -16 | 6 | 0.061 | 4664 | 49.40% Thalamus_R; 24.19% no_label; 17.67% Pallidum_R; 6.69% Caudate_R |
| | rIPL | 32 | -58 | 48 | 0.061 | 2872 | 37.33% Parietal_Inf_R; 31.20% Angular_R; 30.08% Parietal_Sup_R |
| | lPrec | -2 | -56 | 34 | 0.063 | 1520 | 50.53% Precuneus_L; 31.58% Cingulate_Post_L; 15.26% Precuneus_R |

#### Table S5: Information on clusters resulting from ALE and MACM analyses

Data derived from ALE maps after application of a cluster-level threshold at family-wise error-corrected $p < .05$. AtlasReader (*116*) was used to estimate peak MNI coordinates (*X, Y, Z*), the average ALE value (*Estimate*), the cluster volume in mm (*Volume*), and coverage of anatomical regions relative to the cluster according the the AAL atlas (*117*). Abbreviations: ALE = activation likelihood estimation, pseudo = pseudo-hyperscanning, fNIRS = functional near-infrared spectroscopy, MACM = meta-analytic connectivity modelling, AAL = automated anatomic labelling atlas, r/lTPJ = right/left temporoparietal junction, rSTG = right superior temporal gyrus, rIns = right insula, r/lPFCIns = right/left prefrontal cortex-insula, SMA = supplementary motor area, r/lTh = left thalamus, r/lIPL = left inferior parietal lobule, lPrec = left precuneus.

**All included fNIRS experiments**

| Parcel | N (ch) | | | | | | N (exp) | | N (sub) | | Ratio * N (sub-all) | | | | Ratio * N (exp-all) | | | | AAL region |
|---|---|---|---|---|---|---|---|---|---|---|---|---|---|---|---|---|---|---|---|
| | *all* | *sync* | *ratio* | *p* | *p (M)* | *p (%)* | *all* | *sync* | *all* | *sync* | *value* | *p* | *p (M)* | *p (%)* | *value* | *p* | *p (M)* | *p (%)* | |
| RH_Vis_3 | 19 | 4 | 0.21 | 0.169 | 0.320 | 0.024 | 18 | 4 | 1339 | 592 | 281.89 | 0.014 | 0.052 | 0.481 | 3.79 | 0.075 | 0.236 | 0.056 | 52.97% Temporal_Inf_R |
| LH_Default_PFC_2 | 56 | 11 | 0.20 | 0.057 | 0.092 | 0.295 | 44 | 7 | 2205 | 268 | 433.13 | 0.018 | 0.116 | 0.181 | 8.64 | 0.007 | 0.091 | 0.303 | 37.52% Frontal_Inf_Tri_L |
| LH_DorsAttn_Post_6 | 5 | 3 | 0.60 | 0.013 | 0.058 | 0.480 | 4 | 3 | 154 | 118 | 92.40 | 0.047 | 0.149 | 0.114 | 2.40 | 0.012 | 0.102 | 0.288 | 57.52% Parietal_Sup_L |
| RH_SalVentAttn_TempOccPar_1 | 35 | 5 | 0.14 | 0.437 | 0.556 | <0.001 | 31 | 4 | 2145 | 176 | 306.43 | 0.048 | 0.280 | 0.079 | 4.43 | 0.188 | 0.484 | 0.001 | 53.36% Temporal_Sup_R |
| RH_Default_PFCdPFCm_1 | 38 | 6 | 0.16 | 0.264 | 0.477 | 0.011 | 38 | 6 | 1844 | 280 | 291.16 | 0.076 | 0.317 | 0.027 | 6.00 | 0.044 | 0.293 | 0.066 | 29.90% Frontal_Med_Orb_R |
| LH_SalVentAttn_FrOperIns_2 | 4 | 1 | 0.25 | 0.396 | 0.999 | 0.004 | 3 | 1 | 242 | 18 | 60.50 | 0.111 | 0.999 | 0.107 | 0.75 | 0.395 | 0.999 | 0.004 | 64.87% Insula_L |
| RH_Default_PFCdPFCm_2 | 114 | 17 | 0.15 | 0.151 | 0.295 | 0.006 | 46 | 13 | 2818 | 1002 | 420.23 | 0.124 | 0.339 | 0.002 | 6.86 | 0.278 | 0.555 | <0.001 | 41.38% Frontal_Sup_Medial_R |
| LH_Default_Temp_2 | 6 | 1 | 0.17 | 0.544 | 0.768 | <0.001 | 6 | 1 | 444 | 48 | 74.00 | 0.142 | 0.612 | <0.001 | 1.00 | 0.457 | 0.733 | <0.001 | 88.43% Temporal_Mid_L |
| RH_Cont_PFCl_3 | 87 | 12 | 0.14 | 0.319 | 0.349 | 0.034 | 47 | 11 | 2756 | 912 | 380.14 | 0.146 | 0.215 | 0.066 | 6.48 | 0.259 | 0.388 | 0.020 | 65.37% Frontal_Mid_2_R |
| LH_Default_PFC_4 | 113 | 21 | 0.19 | 0.015 | 0.029 | 0.691 | 43 | 12 | 2098 | 506 | 389.89 | 0.205 | 0.228 | 0.007 | 7.99 | 0.102 | 0.132 | 0.086 | 55.26% Frontal_Sup_2_L |
| RH_Cont_PFCl_2 | 61 | 7 | 0.11 | 0.599 | 0.652 | <0.001 | 42 | 5 | 2545 | 206 | 292.05 | 0.222 | 0.377 | 0.014 | 4.82 | 0.413 | 0.557 | 0.001 | 52.90% Frontal_Mid_2_R |
| LH_Cont_Par_1 | 9 | 3 | 0.33 | 0.076 | 0.196 | 0.178 | 6 | 2 | 212 | 66 | 70.67 | 0.258 | 0.270 | 0.024 | 2.00 | 0.186 | 0.225 | 0.124 | 71.29% Parietal_Inf_L |
| RH_Cont_Par_2 | 63 | 9 | 0.14 | 0.311 | 0.397 | 0.002 | 32 | 8 | 2015 | 503 | 287.86 | 0.263 | 0.476 | 0.001 | 4.57 | 0.459 | 0.650 | <0.001 | 59.87% Angular_R |
| RH_Limbic_OFC_1 | 4 | 1 | 0.25 | 0.376 | 0.404 | 0.056 | 4 | 1 | 146 | 18 | 36.50 | 0.263 | 0.248 | 0.080 | 1.00 | 0.341 | 0.241 | 0.104 | 28.24% Rectus_R |
| RH_Default_Temp_3 | 10 | 1 | 0.10 | 0.713 | 0.761 | <0.001 | 9 | 1 | 734 | 180 | 73.40 | 0.278 | 0.456 | 0.028 | 0.90 | 0.657 | 0.660 | 0.004 | 48.97% Temporal_Mid_R |
| RH_Default_Temp_2 | 4 | 1 | 0.25 | 0.378 | 0.504 | <0.001 | 4 | 1 | 131 | 88 | 32.75 | 0.297 | 0.430 | <0.001 | 1.00 | 0.343 | 0.470 | <0.001 | 43.90% Temporal_Pole_Sup_R |
| LH_Default_PFC_6 | 9 | 2 | 0.22 | 0.300 | 0.287 | 0.063 | 8 | 2 | 308 | 66 | 68.44 | 0.301 | 0.304 | 0.003 | 1.78 | 0.283 | 0.232 | 0.066 | 73.85% Frontal_Mid_2_L |
| RH_Default_PFCv_2 | 23 | 5 | 0.22 | 0.117 | 0.038 | 0.590 | 14 | 3 | 631 | 79 | 137.17 | 0.307 | 0.135 | 0.107 | 3.04 | 0.241 | 0.103 | 0.188 | 49.04% Frontal_Inf_Tri_R |
| LH_Cont_PFCl_1 | 63 | 8 | 0.13 | 0.444 | 0.401 | 0.021 | 45 | 7 | 2075 | 226 | 263.49 | 0.337 | 0.384 | 0.007 | 5.71 | 0.232 | 0.325 | 0.030 | 54.13% Frontal_Inf_Tri_L |
| LH_SalVentAttn_ParOper_1 | 18 | 5 | 0.28 | 0.053 | 0.183 | 0.108 | 12 | 4 | 399 | 148 | 110.83 | 0.341 | 0.609 | 0.001 | 3.33 | 0.107 | 0.385 | 0.008 | 54.68% SupraMarginal_L |
| RH_Default_Par_1 | 48 | 5 | 0.10 | 0.688 | 0.718 | <0.001 | 33 | 4 | 2063 | 120 | 214.90 | 0.384 | 0.481 | 0.005 | 3.44 | 0.602 | 0.702 | <0.001 | 39.79% Angular_R |
| RH_DorsAttn_PrCv_1 | 22 | 5 | 0.23 | 0.096 | 0.400 | <0.001 | 16 | 5 | 513 | 186 | 116.59 | 0.389 | 0.499 | <0.001 | 3.64 | 0.135 | 0.417 | <0.001 | 44.81% Frontal_Inf_Oper_R |
| LH_DorsAttn_Post_2 | 18 | 3 | 0.17 | 0.339 | 0.369 | 0.013 | 14 | 3 | 531 | 102 | 88.50 | 0.427 | 0.517 | <0.001 | 2.33 | 0.329 | 0.379 | 0.007 | 29.06% Parietal_Inf_L |
| LH_SalVentAttn_PFCl_1 | 76 | 10 | 0.13 | 0.384 | 0.551 | <0.001 | 43 | 10 | 2042 | 436 | 268.68 | 0.440 | 0.674 | <0.001 | 5.66 | 0.338 | 0.607 | <0.001 | 58.16% Frontal_Mid_2_L |
| LH_Default_Par_1 | 16 | 2 | 0.13 | 0.579 | 0.296 | 0.042 | 15 | 2 | 647 | 197 | 80.88 | 0.455 | 0.288 | 0.017 | 1.88 | 0.401 | 0.262 | 0.046 | 69.08% Temporal_Mid_L |
| LH_Vis_7 | 6 | 1 | 0.17 | 0.503 | 0.666 | <0.001 | 6 | 1 | 163 | 48 | 27.17 | 0.478 | 0.661 | <0.001 | 1.00 | 0.419 | 0.661 | <0.001 | 42.60% Temporal_Mid_L |
| RH_Vis_7 | 31 | 4 | 0.13 | 0.496 | 0.587 | 0.004 | 20 | 3 | 1040 | 119 | 134.19 | 0.493 | 0.537 | 0.013 | 2.58 | 0.514 | 0.646 | 0.001 | 74.44% Occipital_Mid_R |
| RH_SomMot_4 | 23 | 3 | 0.13 | 0.494 | 0.754 | <0.001 | 17 | 3 | 741 | 65 | 96.65 | 0.514 | 0.702 | <0.001 | 2.22 | 0.477 | 0.714 | <0.001 | 62.76% Postcentral_R |
| LH_DorsAttn_Post_3 | 8 | 1 | 0.13 | 0.641 | 0.549 | 0.007 | 4 | 1 | 138 | 18 | 17.25 | 0.641 | 0.608 | <0.001 | 0.50 | 0.641 | 0.602 | <0.001 | 48.76% Parietal_Sup_L |
| LH_SomMot_5 | 19 | 4 | 0.21 | 0.165 | 0.223 | 0.055 | 6 | 4 | 246 | 166 | 51.79 | 0.722 | 0.705 | <0.001 | 1.26 | 0.69 | 0.654 | <0.001 | 55.07% Postcentral_L |
| RH_Default_PFCdPFCm_3 | 22 | 2 | 0.09 | 0.736 | 0.813 | <0.001 | 13 | 1 | 584 | 32 | 53.09 | 0.759 | 0.743 | <0.001 | 1.18 | 0.736 | 0.780 | <0.001 | 61.82% Frontal_Sup_2_R |
| RH_SomMot_1 | 31 | 2 | 0.06 | 0.895 | 0.926 | <0.001 | 25 | 2 | 1333 | 105 | 86.00 | 0.766 | 0.807 | <0.001 | 1.61 | 0.793 | 0.882 | <0.001 | 61.86% Temporal_Sup_R |
| RH_DorsAttn_Post_3 | 19 | 1 | 0.05 | 0.915 | 0.824 | <0.001 | 17 | 1 | 826 | 32 | 43.47 | 0.777 | 0.744 | <0.001 | 0.89 | 0.774 | 0.808 | <0.001 | 42.84% Parietal_Inf_R |
| LH_SomMot_4 | 25 | 2 | 0.08 | 0.812 | 0.930 | <0.001 | 18 | 2 | 657 | 44 | 52.56 | 0.832 | 0.928 | <0.001 | 1.44 | 0.802 | 0.927 | <0.001 | 73.10% Postcentral_L |
| RH_Cont_Par_1 | 49 | 3 | 0.06 | 0.940 | 0.887 | <0.001 | 30 | 3 | 1845 | 550 | 112.96 | 0.855 | 0.808 | <0.001 | 1.84 | 0.936 | 0.875 | <0.001 | 59.43% SupraMarginal_R |

| | | | | | | | | | | | | | | | | | | | |
|---|---|---|---|---|---|---|---|---|---|---|---|---|---|---|---|---|---|---|---|
| RH_DorsAttn_Post_5 | 25 | 2 | 0.08 | 0.831 | 0.806 | <0.001 | 15 | 2 | 606 | 49 | 48.48 | 0.859 | 0.804 | <0.001 | 1.20 | 0.831 | 0.803 | <0.001 | 46.68% Precuneus_R |
| RH_DorsAttn_Post_2 | 17 | 1 | 0.06 | 0.894 | 0.887 | <0.001 | 12 | 1 | 534 | 44 | 31.41 | 0.860 | 0.827 | <0.001 | 0.71 | 0.891 | 0.853 | 0.001 | 85.28% Postcentral_R |
| RH_SalVentAttn_TempOccPar_2 | 50 | 4 | 0.08 | 0.835 | 0.780 | <0.001 | 29 | 3 | 1295 | 112 | 103.60 | 0.884 | 0.822 | <0.001 | 2.32 | 0.838 | 0.799 | <0.001 | 68.61% SupraMarginal_R |
| LH_Default_Par_2 | 47 | 10 | 0.21 | 0.037 | 0.020 | 0.741 | 13 | 6 | 447 | 132 | 95.11 | 0.890 | 0.878 | <0.001 | 2.77 | 0.743 | 0.685 | <0.001 | 36.97% Angular_L |
| RH_DorsAttn_Post_1 | 47 | 3 | 0.06 | 0.918 | 0.553 | 0.005 | 26 | 2 | 1365 | 152 | 87.13 | 0.905 | 0.432 | 0.018 | 1.66 | 0.916 | 0.675 | <0.001 | 64.75% Temporal_Mid_R |
| RH_DorsAttn_Post_4 | 40 | 3 | 0.08 | 0.871 | 0.945 | <0.001 | 20 | 3 | 978 | 191 | 73.35 | 0.915 | 0.957 | <0.001 | 1.50 | 0.907 | 0.956 | <0.001 | 56.13% Parietal_Sup_R |
| RH_Cont_PFCl_1 | 169 | 19 | 0.11 | 0.617 | 0.491 | <0.001 | 43 | 10 | 2052 | 398 | 230.70 | 0.959 | 0.813 | <0.001 | 4.83 | 0.913 | 0.768 | <0.001 | 44.35% Frontal_Sup_2_R |
| LH_SomMot_1 | 28 | 1 | 0.04 | 0.968 | 0.931 | <0.001 | 17 | 1 | 747 | 18 | 26.68 | 0.967 | 0.919 | <0.001 | 0.61 | 0.968 | 0.929 | <0.001 | 77.45% Temporal_Sup_L |
| RH_SomMot_6 | 50 | 3 | 0.06 | 0.940 | 0.955 | <0.001 | 21 | 2 | 1022 | 50 | 61.32 | 0.977 | 0.951 | <0.001 | 1.26 | 0.987 | 0.955 | <0.001 | 54.07% Precentral_R |
| LH_Default_PFC_5 | 159 | 9 | 0.06 | 0.999 | 0.993 | <0.001 | 47 | 7 | 2226 | 350 | 126.00 | 0.999 | 0.997 | <0.001 | 2.66 | 0.999 | 0.996 | <0.001 | 51.49% Frontal_Sup_Medial_L |

**fNIRS experiments restricted to specific INS > rest, control, or randomization**

| Parcel | N (ch) | | | | N (exp) | | N (sub) | | Ratio * N (sub-all) | | Ratio * N (exp-all) | | AAL region |
|---|---|---|---|---|---|---|---|---|---|---|---|---|---|
| | *all* | *sync* | *ratio* | *p* | *all* | *sync* | *all* | *sync* | *value* | *p* | *value* | *p* | |
| LH_Default_PFC_4 | 100 | 19 | 0.19 | 0.008 | 33 | 6 | 1514 | 280 | 275.27 | 0.008 | 7.03 | 0.051 | 55.26% Frontal_Sup_2_L |
| RH_Cont_PFCl_3 | 76 | 11 | 0.14 | 0.205 | 4 | 3 | 154 | 118 | 92.40 | 0.021 | 5.93 | 0.130 | 65.37% Frontal_Mid_2_R |
| RH_Default_PFCdPFCm_1 | 33 | 6 | 0.18 | 0.147 | 37 | 11 | 1704 | 484 | 323.76 | 0.027 | 6.00 | 0.015 | 29.90% Frontal_Med_Orb_R |
| RH_Default_PFCdPFCm_2 | 95 | 12 | 0.13 | 0.357 | 37 | 5 | 1613 | 216 | 245.46 | 0.052 | 5.05 | 0.384 | 41.38% Frontal_Sup_Medial_R |
| LH_Default_PFC_2 | 46 | 7 | 0.15 | 0.246 | 41 | 10 | 1946 | 396 | 281.66 | 0.063 | 5.63 | 0.048 | 37.52% Frontal_Inf_Tri_L |
| LH_SalVentAttn_PFCl_1 | 60 | 8 | 0.13 | 0.315 | 15 | 3 | 637 | 76 | 127.40 | 0.088 | 4.40 | 0.306 | 58.16% Frontal_Mid_2_L |
| RH_Cont_PFCl_2 | 50 | 6 | 0.12 | 0.465 | 5 | 2 | 182 | 66 | 78.00 | 0.090 | 4.44 | 0.210 | 52.90% Frontal_Mid_2_R |
| LH_Cont_PFCl_1 | 46 | 6 | 0.13 | 0.386 | 23 | 3 | 1059 | 154 | 162.92 | 0.112 | 4.57 | 0.176 | 54.13% Frontal_Inf_Tri_L |
| RH_Cont_PFCl_1 | 154 | 17 | 0.11 | 0.544 | 35 | 5 | 1569 | 174 | 204.65 | 0.144 | 4.19 | 0.843 | 44.35% Frontal_Sup_2_R |
| RH_SalVentAttn_TempOccPar_1 | 26 | 4 | 0.15 | 0.313 | 37 | 5 | 1727 | 206 | 207.24 | 0.153 | 3.54 | 0.143 | 53.36% Temporal_Sup_R |
| RH_Cont_Par_2 | 51 | 7 | 0.14 | 0.343 | 4 | 1 | 176 | 48 | 44.00 | 0.156 | 3.57 | 0.475 | 59.87% Angular_R |
| RH_Vis_3 | 15 | 3 | 0.20 | 0.225 | 2 | 1 | 62 | 18 | 31.00 | 0.161 | 3.00 | 0.081 | 52.97% Temporal_Inf_R |
| RH_Default_Par_1 | 35 | 3 | 0.09 | 0.783 | 40 | 10 | 1972 | 428 | 249.09 | 0.207 | 2.23 | 0.634 | 39.79% Angular_R |
| RH_Default_PFCv_2 | 16 | 5 | 0.31 | 0.039 | 33 | 8 | 1566 | 382 | 208.80 | 0.217 | 3.13 | 0.100 | 49.04% Frontal_Inf_Tri_R |
| LH_DorsAttn_Post_6 | 5 | 3 | 0.60 | 0.007 | 4 | 1 | 146 | 18 | 36.50 | 0.224 | 2.40 | 0.006 | 57.52% Parietal_Sup_L |
| RH_Vis_7 | 27 | 3 | 0.11 | 0.579 | 10 | 3 | 311 | 79 | 97.19 | 0.238 | 1.78 | 0.587 | 74.44% Occipital_Mid_R |
| RH_DorsAttn_Post_1 | 36 | 3 | 0.08 | 0.769 | 11 | 3 | 365 | 134 | 73.00 | 0.369 | 1.75 | 0.768 | 64.75% Temporal_Mid_R |
| LH_Default_PFC_5 | 135 | 6 | 0.04 | 0.997 | 9 | 2 | 311 | 90 | 51.83 | 0.389 | 1.69 | 0.999 | 51.49% Frontal_Sup_Medial_L |
| LH_Default_Par_2 | 35 | 9 | 0.26 | 0.013 | 6 | 1 | 163 | 48 | 27.17 | 0.456 | 2.57 | 0.502 | 36.97% Angular_L |
| LH_Cont_Par_1 | 7 | 3 | 0.43 | 0.035 | 26 | 6 | 1131 | 361 | 155.24 | 0.461 | 2.14 | 0.035 | 71.29% Parietal_Inf_L |
| RH_DorsAttn_PrCv_1 | 15 | 3 | 0.20 | 0.231 | 12 | 2 | 415 | 43 | 51.88 | 0.541 | 2.20 | 0.227 | 44.81% Frontal_Inf_Oper_R |
| RH_SomMot_6 | 46 | 3 | 0.07 | 0.901 | 16 | 2 | 760 | 55 | 84.44 | 0.548 | 1.24 | 0.939 | 54.07% Precentral_R |
| RH_SomMot_4 | 16 | 2 | 0.13 | 0.544 | 4 | 1 | 138 | 18 | 17.25 | 0.591 | 1.50 | 0.530 | 62.76% Postcentral_R |
| LH_SalVentAttn_ParOper_1 | 12 | 2 | 0.17 | 0.376 | 26 | 3 | 1157 | 98 | 99.17 | 0.601 | 1.50 | 0.371 | 54.68% SupraMarginal_L |

| | | | | | | | | | | | | | | |
|---|---|---|---|---|---|---|---|---|---|---|---|---|---|---|
| LH_SomMot_5 | 19 | 4 | 0.21 | 0.133 | 13 | 2 | 417 | 44 | 46.33 | 0.614 | 1.26 | 0.636 | 55.07% | Postcentral_L |
| RH_DorsAttn_Post_5 | 25 | 2 | 0.08 | 0.800 | 6 | 4 | 246 | 166 | 51.79 | 0.632 | 1.20 | 0.800 | 46.68% | Precuneus_R |
| LH_SomMot_4 | 18 | 2 | 0.11 | 0.594 | 17 | 1 | 826 | 32 | 43.47 | 0.669 | 1.44 | 0.588 | 73.10% | Postcentral_L |
| LH_Default_Temp_2 | 4 | 1 | 0.25 | 0.366 | 10 | 5 | 307 | 110 | 78.94 | 0.725 | 1.00 | 0.333 | 88.43% | Temporal_Mid_L |
| RH_DorsAttn_Post_4 | 36 | 2 | 0.06 | 0.909 | 21 | 2 | 997 | 152 | 83.08 | 0.733 | 1.00 | 0.909 | 56.13% | Parietal_Sup_R |
| RH_DorsAttn_Post_3 | 19 | 1 | 0.05 | 0.891 | 11 | 1 | 327 | 17 | 27.25 | 0.740 | 0.89 | 0.714 | 42.84% | Parietal_Inf_R |
| RH_Limbic_OFC_1 | 4 | 1 | 0.25 | 0.376 | 10 | 1 | 355 | 44 | 27.31 | 0.744 | 1.00 | 0.346 | 28.24% | Rectus_R |
| RH_SomMot_1 | 21 | 1 | 0.05 | 0.913 | 18 | 1 | 763 | 17 | 36.33 | 0.789 | 0.86 | 0.785 | 61.86% | Temporal_Sup_R |
| RH_DorsAttn_Post_2 | 17 | 1 | 0.06 | 0.861 | 15 | 2 | 606 | 49 | 48.48 | 0.794 | 0.71 | 0.856 | 85.28% | Postcentral_R |
| LH_SalVentAttn_FrOperIns_2 | 2 | 1 | 0.50 | 0.219 | 12 | 1 | 534 | 44 | 31.41 | 0.801 | 1.00 | 0.216 | 64.87% | Insula_L |
| RH_Cont_Par_1 | 40 | 1 | 0.03 | 0.991 | 38 | 9 | 1722 | 376 | 190.09 | 0.804 | 0.63 | 0.984 | 59.43% | SupraMarginal_R |
| LH_DorsAttn_Post_2 | 13 | 1 | 0.08 | 0.774 | 18 | 2 | 792 | 59 | 44.00 | 0.908 | 0.77 | 0.760 | 29.06% | Parietal_Inf_L |
| LH_Default_Par_1 | 12 | 1 | 0.08 | 0.758 | 12 | 1 | 391 | 18 | 18.62 | 0.928 | 0.92 | 0.627 | 69.08% | Temporal_Mid_L |
| LH_Vis_7 | 6 | 1 | 0.17 | 0.497 | 19 | 2 | 836 | 50 | 54.52 | 0.936 | 1.00 | 0.391 | 42.60% | Temporal_Mid_L |
| RH_SalVentAttn_TempOccPar_2 | 39 | 1 | 0.03 | 0.988 | 25 | 1 | 1113 | 34 | 27.83 | 0.977 | 0.59 | 0.988 | 68.61% | SupraMarginal_R |
| LH_SomMot_1 | 21 | 1 | 0.05 | 0.928 | 23 | 1 | 905 | 26 | 23.21 | 0.987 | 0.57 | 0.928 | 77.45% | Temporal_Sup_L |
| LH_DorsAttn_Post_3 | 8 | 1 | 0.13 | 0.591 | 38 | 5 | 1778 | 242 | 79.02 | 0.997 | 0.50 | 0.591 | 48.76% | Parietal_Sup_L |

**Table S6: Region-wise fNIRS meta-analyses**

Results from regionwise meta-analysis of fNIRS INS experiments, for the complete and for a restricted set of experiments. *Parcel*: brain region label drawn from a functionally defined 100-parcel cortical parcellation (*21*). *N (ch/exp/sub)*: number of channels, experiments, and subjects associated to the respective region. *Ratio*: (number of INS channels / number of all channels). *Ratio * N (sub-all)*: (Ratio * number of subjects contributing to a parcel). *Ratio * N (exp-all)*: (Ratio * number of experiments contributing to a parcel). *p (M)*: Median *p* value resulting from iterative (1,000 iterations) recalculation of fNIRS meta-analysis after fNIRS coordinate randomization within a 1cm cortical radius. *p (%)*: Percentage of sub-threshold (< 0.05) *p*-values after coordinate randomization. Randomization analysis was only performed for the main fNIRS coordinate dataset. *P* values are estimated from null distributions generated by permuting (5,000 iterations) the fNIRS channel coordinate-atlas parcel assignment and recalculating the respective index. *AAL region*: Anatomical region with the largest coverage from the automated anatomic labelling atlas (*117*). The order is based on *p* values associated to "*Ratio * N (sub-all)*" and only those parcels in which at least one INS channel was observed are shown.
Abbreviations: fNIRS = functional near-infrared spectroscopy, INS = interpersonal neural synchronization.

| Topic | rTPJ | | | | | | Whole-brain | | |
|---|---|---|---|---|---|---|---|---|---|
| | Reverse inference | | | Forward inference | | | Spearman correlation | | |
| | q | z | prob. | q | z | lik. | p | q | Z(rho) |
| 145_mind_mental_social | <0.001 | 6.81 | 0.02 | <0.001 | 5.40 | 1.91 | 0.379 | 0.679 | 0.06 |
| 143_action_actions_observation | <0.001 | 5.32 | 0.02 | <0.001 | 3.71 | 1.56 | <0.001 | 0.005 | 0.64 |
| 82_motion_mt_moving | <0.001 | 5.25 | 0.01 | <0.001 | 4.29 | 1.84 | 0.001 | 0.007 | 0.54 |
| 193_mirror_video_imitation | 0.007 | 2.68 | 0.01 | 0.211 | 1.25 | 1.26 | 0.001 | 0.009 | 0.51 |
| 115_face_faces_fusiform | 0.011 | 2.55 | 0.02 | 0.048 | 1.98 | 1.34 | 0.224 | 0.498 | 0.18 |
| 154_social_interactions_interaction | 0.012 | 2.50 | 0.02 | 0.012 | 2.51 | 1.43 | 0.777 | 1.000 | -0.19 |
| 138_real_virtual_reality | 0.021 | 2.31 | 0.01 | 0.027 | -2.21 | 1.54 | 0.012 | 0.075 | 0.44 |
| 30_events_future_personal | 0.072 | 1.80 | 0.01 | 0.090 | -1.70 | 1.45 | 0.838 | 1.000 | -0.23 |
| 99_detection_novelty_oddball | 0.072 | 1.80 | 0.01 | 0.025 | -2.24 | 1.71 | 0.187 | 0.458 | 0.17 |
| 175_dopamine_dopaminergic_striatum | 0.145 | -1.46 | 0.00 | 0.997 | 0.00 | 0.58 | 1.000 | 1.000 | -0.66 |
| 149_olfactory_taste_odor | 0.204 | -1.27 | 0.00 | 0.997 | 0.00 | 0.34 | 0.992 | 1.000 | -0.46 |
| 100_gestures_abstract_race | 0.243 | 1.17 | 0.01 | 0.210 | -1.25 | 1.43 | 0.042 | 0.146 | 0.38 |
| 142_scene_scenes_perspective | 0.243 | 1.17 | 0.01 | 0.217 | -1.23 | 1.30 | 0.242 | 0.512 | 0.17 |
| 26_time_delay_temporal | 0.243 | 1.17 | 0.02 | 0.217 | 1.23 | 1.17 | 0.381 | 0.679 | 0.07 |
| 64_attention_attentional_visual | 0.244 | 1.16 | 0.03 | 0.217 | 1.23 | 1.14 | <0.001 | 0.005 | 0.82 |
| 139_faces_emotional_facial | 0.246 | 1.16 | 0.02 | 0.142 | 1.47 | 1.28 | 0.881 | 1.000 | -0.22 |
| 157_somatosensory_stimulation_cortex | 0.248 | -1.15 | 0.01 | 0.997 | 0.00 | 0.65 | 0.394 | 0.689 | 0.07 |
| 191_eye_gaze_saccade | 0.248 | 1.15 | 0.01 | 0.437 | -0.78 | 1.17 | 0.059 | 0.180 | 0.35 |
| 97_adaptation_selective_stimulus | 0.248 | 1.15 | 0.03 | 0.210 | 1.25 | 1.16 | <0.001 | 0.005 | 0.64 |
| 8_cues_cue_target | 0.252 | 1.14 | 0.01 | 0.335 | 0.96 | 1.17 | 0.041 | 0.146 | 0.35 |
| 181_creative_creativity_generation | 0.261 | 1.12 | 0.00 | 0.185 | -1.32 | 1.56 | 0.435 | 0.719 | 0.04 |
| 130_risk_high_taking | 0.292 | -1.05 | 0.00 | 0.997 | 0.00 | 0.72 | 0.998 | 1.000 | -0.50 |
| 74_feedback_negative_performance | 0.292 | 1.05 | 0.01 | 0.368 | -0.90 | 1.22 | 0.470 | 0.750 | 0.03 |
| 11_cognitive_function_performance | 0.292 | -1.05 | 0.05 | 0.906 | 0.12 | 0.98 | 0.523 | 0.821 | -0.02 |
| 108_visual_auditory_sensory | 0.387 | 0.87 | 0.02 | 0.449 | 0.76 | 1.10 | <0.001 | 0.005 | 0.61 |
| 111_memory_encoding_hippocampal | 0.387 | -0.87 | 0.01 | 0.997 | 0.00 | 0.84 | 0.835 | 1.000 | -0.21 |
| 118_goal_goals_planning | 0.387 | -0.87 | 0.00 | 0.997 | 0.00 | 0.65 | 0.744 | 1.000 | -0.15 |
| 125_expertise_experts_ic | 0.387 | -0.87 | 0.00 | 0.997 | 0.00 | 0.67 | 0.026 | 0.125 | 0.36 |
| 152_perceptual_perception_visual | 0.387 | -0.87 | 0.01 | 0.997 | 0.00 | 0.87 | 0.001 | 0.008 | 0.56 |
| 159_inhibition_response_inhibitory | 0.387 | 0.87 | 0.01 | 0.217 | 1.23 | 1.23 | 0.461 | 0.749 | 0.03 |
| 161_women_men_sex | 0.387 | -0.87 | 0.01 | 0.997 | 0.00 | 0.81 | 0.999 | 1.000 | -0.49 |
| 166_pictures_picture_images | 0.387 | -0.87 | 0.01 | 0.997 | 0.00 | 0.83 | 0.928 | 1.000 | -0.20 |
| 184_spatial_location_space | 0.387 | 0.87 | 0.02 | 0.476 | 0.71 | 1.09 | 0.010 | 0.073 | 0.48 |
| 198_repetition_priming_suppression | 0.387 | -0.87 | 0.01 | 0.997 | 0.00 | 0.84 | 0.057 | 0.180 | 0.36 |
| 3_task_switching_set | 0.387 | 0.87 | 0.01 | 0.515 | -0.65 | 1.11 | 0.100 | 0.281 | 0.27 |
| 39_light_shed_alertness | 0.387 | 0.87 | 0.01 | 0.335 | -0.96 | 1.27 | 0.688 | 1.000 | -0.13 |
| 55_music_musical_pitch | 0.387 | -0.87 | 0.00 | 0.997 | 0.00 | 0.62 | 0.425 | 0.717 | 0.05 |
| 6_implicit_explicit_cd | 0.387 | 0.87 | 0.01 | 0.216 | -1.24 | 1.43 | 0.364 | 0.676 | 0.06 |
| 196_personality_trait_scores | 0.433 | -0.78 | 0.01 | 0.872 | -0.16 | 0.96 | 0.998 | 1.000 | -0.48 |
| 79_training_practice_trained | 0.433 | -0.78 | 0.01 | 0.997 | 0.00 | 0.85 | 0.338 | 0.641 | 0.11 |
| 187_speech_auditory_temporal | 0.437 | 0.78 | 0.02 | 0.599 | 0.53 | 1.06 | 0.212 | 0.494 | 0.18 |
| 31_hearing_deaf_sign | 0.442 | 0.77 | 0.00 | 0.416 | -0.81 | 1.33 | 0.025 | 0.125 | 0.40 |
| 28_memory_retrieval_episodic | 0.448 | 0.76 | 0.01 | 0.745 | 0.33 | 1.03 | 0.836 | 1.000 | -0.23 |
| 197_reward_striatum_anticipation | 0.466 | -0.73 | 0.01 | 0.906 | 0.12 | 0.96 | 1.000 | 1.000 | -0.58 |
| 117_object_objects_visual | 0.484 | 0.70 | 0.01 | 0.449 | 0.76 | 1.12 | 0.030 | 0.125 | 0.42 |
| 58_autonomic_arousal_rate | 0.484 | -0.70 | 0.00 | 0.997 | 0.00 | 0.76 | 0.988 | 1.000 | -0.44 |
| 20_wm_load_memory | 0.491 | -0.69 | 0.01 | 0.997 | 0.00 | 0.79 | 0.191 | 0.458 | 0.19 |
| 93_language_sentences_comprehension | 0.491 | 0.69 | 0.01 | 0.449 | 0.76 | 1.12 | 0.102 | 0.281 | 0.28 |
| 128_prediction_error_outcome | 0.500 | 0.67 | 0.01 | 0.281 | 1.08 | 1.21 | 0.762 | 1.000 | -0.12 |
| 137_regulation_emotion_reappraisal | 0.503 | 0.67 | 0.01 | 0.449 | -0.76 | 1.16 | 0.997 | 1.000 | -0.36 |
| 182_experience_subjective_ratings | 0.503 | 0.67 | 0.01 | 0.416 | 0.81 | 1.14 | 0.967 | 1.000 | -0.28 |
| 5_body_bodies_eba | 0.503 | 0.67 | 0.01 | 0.335 | -0.96 | 1.29 | 0.049 | 0.165 | 0.37 |
| 127_task_performance_cognitive | 0.513 | -0.65 | 0.14 | 0.821 | 0.23 | 1.00 | 0.029 | 0.125 | 0.34 |
| 158_pet_glucose_metabolism | 0.520 | 0.64 | 0.00 | 0.772 | -0.29 | 1.01 | 1.000 | 1.000 | -0.56 |
| 95_verbs_verb_nouns | 0.567 | 0.57 | 0.00 | 0.745 | -0.33 | 1.03 | 0.056 | 0.180 | 0.34 |
| 186_decision_making_choice | 0.625 | 0.49 | 0.02 | 0.449 | 0.76 | 1.11 | 0.811 | 1.000 | -0.19 |

| | | | | | | | | | |
|---|---|---|---|---|---|---|---|---|---|
| 23_reasoning_relational_relations | 0.628 | 0.48 | 0.00 | 0.452 | -0.75 | 1.21 | 0.289 | 0.585 | 0.14 |
| 173_touch_ct_tactile | 0.630 | -0.48 | 0.00 | 0.997 | 0.00 | 0.72 | 0.010 | 0.073 | 0.47 |
| 43_conflict_interference_incongruent | 0.630 | 0.48 | 0.01 | 0.519 | 0.64 | 1.10 | 0.023 | 0.125 | 0.40 |
| 146_pain_painful_chronic | 0.657 | -0.44 | 0.01 | 0.997 | 0.00 | 0.78 | 0.903 | 1.000 | -0.29 |
| 18_color_shape_shapes | 0.718 | 0.36 | 0.01 | 0.449 | -0.76 | 1.20 | 0.001 | 0.007 | 0.55 |
| 57_learning_learned_sequence | 0.718 | -0.36 | 0.01 | 0.997 | 0.00 | 0.85 | 0.791 | 1.000 | -0.17 |
| 90_imagery_mental_rotation | 0.718 | -0.36 | 0.00 | 0.997 | 0.00 | 0.74 | 0.093 | 0.272 | 0.33 |
| 150_reading_phonological_readers | 0.730 | -0.35 | 0.01 | 0.997 | 0.00 | 0.82 | 0.001 | 0.007 | 0.57 |
| 153_phase_women_menstrual | 0.853 | -0.19 | 0.00 | 0.916 | -0.11 | 0.87 | 0.947 | 1.000 | -0.33 |
| 172_verbal_fluency_overt | 0.853 | -0.19 | 0.01 | 0.997 | 0.00 | 0.81 | 0.030 | 0.125 | 0.37 |
| 179_memory_working_task | 0.853 | -0.19 | 0.01 | 0.957 | 0.05 | 0.94 | 0.137 | 0.346 | 0.24 |
| 2_problem_problems_arithmetic | 0.853 | -0.19 | 0.00 | 0.957 | -0.05 | 0.88 | 0.134 | 0.346 | 0.26 |
| 52_control_cognitive_task | 0.853 | -0.19 | 0.01 | 0.821 | 0.23 | 0.99 | 0.323 | 0.625 | 0.11 |
| 121_tool_tools_knowledge | 0.857 | 0.18 | 0.00 | 0.636 | -0.47 | 1.11 | 0.035 | 0.136 | 0.41 |
| 13_fear_conditioning_extinction | 0.857 | 0.18 | 0.00 | 0.732 | -0.34 | 1.05 | 0.999 | 1.000 | -0.44 |
| 131_test_performance_intelligence | 0.857 | 0.18 | 0.01 | 0.599 | -0.53 | 1.10 | 0.855 | 1.000 | -0.23 |
| 160_bias_biases_spd | 0.857 | -0.18 | 0.00 | 0.821 | -0.23 | 0.97 | 0.825 | 1.000 | -0.14 |
| 163_language_chinese_english | 0.857 | -0.18 | 0.01 | 0.957 | -0.05 | 0.90 | 0.010 | 0.073 | 0.47 |
| 167_human_humans_animal | 0.857 | 0.18 | 0.01 | 0.817 | 0.23 | 1.00 | 0.814 | 1.000 | -0.20 |
| 177_mood_rumination_induction | 0.857 | 0.18 | 0.00 | 0.519 | -0.64 | 1.18 | 0.983 | 1.000 | -0.38 |
| 180_amygdala_threat_fear | 0.857 | -0.18 | 0.01 | 0.821 | -0.23 | 0.99 | 0.998 | 1.000 | -0.45 |
| 65_recognition_correct_familiarity | 0.857 | 0.18 | 0.01 | 0.719 | -0.36 | 1.05 | 0.304 | 0.602 | 0.11 |
| 9_semantic_word_knowledge | 0.857 | -0.18 | 0.01 | 0.965 | 0.04 | 0.92 | 0.286 | 0.585 | 0.13 |
| 42_negative_positive_vmpfc | 0.891 | 0.14 | 0.01 | 0.599 | 0.53 | 1.07 | 1.000 | 1.000 | -0.58 |
| 116_task_matching_strategy | 0.912 | -0.11 | 0.01 | 0.906 | -0.12 | 0.95 | 0.015 | 0.092 | 0.42 |
| 91_movement_motor_movements | 0.926 | -0.09 | 0.01 | 0.997 | 0.00 | 0.74 | 0.221 | 0.498 | 0.22 |
| 92_executive_control_functions | 0.926 | -0.09 | 0.01 | 0.957 | -0.05 | 0.91 | 0.403 | 0.693 | 0.06 |
| 183_orientation_colour_separation | 0.959 | -0.05 | 0.00 | 0.821 | -0.23 | 0.97 | 0.238 | 0.512 | 0.17 |
| 162_words_word_lexical | 0.960 | -0.05 | 0.01 | 0.997 | 0.00 | 0.87 | 0.029 | 0.125 | 0.39 |
| 169_acupuncture_stimulation_vstm | 0.960 | -0.05 | 0.00 | 0.946 | -0.07 | 0.82 | 0.816 | 1.000 | -0.18 |
| 112_context_empathy_contextual | 0.962 | 0.05 | 0.01 | 0.722 | -0.36 | 1.05 | 0.759 | 1.000 | -0.17 |
| 147_target_search_targets | 0.963 | -0.05 | 0.01 | 0.821 | 0.23 | 0.99 | 0.036 | 0.136 | 0.34 |
| 189_category_categories_categorization | 0.974 | 0.03 | 0.01 | 0.745 | -0.33 | 1.03 | 0.117 | 0.314 | 0.28 |
| 19_illusion_physical_perceived | 0.974 | 0.03 | 0.00 | 0.772 | -0.29 | 1.02 | <0.001 | 0.007 | 0.53 |
| 66_emotional_negative_amygdala | 0.988 | 0.02 | 0.02 | 0.775 | 0.29 | 1.01 | 0.998 | 1.000 | -0.45 |

**Table S7: Functional-decoding results**

Functional decoding of the rTPJ cluster using reverse and forward inference estimates, and the whole-brain INS distributions using spatial Spearman correlations (Z-transformed), sorted by reverse inference *q* values. Topics are selected from 200 latent Dirichlet allocation topics distributed with the Neurosynth database (version 7). For whole-brain correlations, *p* values are derived from comparison to correlations with topicwise null maps (10,000 permutations) and were false discovery rate-corrected across all topic (*q*). All associations significant after multiple comparison correction in any analysis are marked in bold.

Abbreviations: rTPJ = right temporo-parietal junction, prob. = probability, lik. = likelihood.

**In-vivo & post-mortem neurotransmitter atlases**

| Atlas | Whole-brain | | | Cortex-only | | Baseline-adjusted | |
|---|---|---|---|---|---|---|---|
| | *Z (ρ)* | *p* | *q* | *Z (ρ)* | *p* | *Z (ρ)* | *p* |
| GABAa | 0.51 | <0.001 | 0.005 | 0.43 | 0.002 | 0.51 | <0.001 |
| SV2a | 0.40 | 0.002 | 0.022 | 0.29 | 0.047 | 0.41 | 0.001 |
| mGluR5 | 0.37 | 0.003 | 0.026 | 0.26 | 0.089 | 0.36 | 0.005 |
| 5HT2a | 0.43 | 0.011 | 0.065 | 0.19 | 0.160 | 0.49 | 0.005 |
| M1 | 0.27 | 0.032 | 0.154 | 0.28 | 0.063 | 0.26 | 0.040 |
| 5HT6 | 0.19 | 0.084 | 0.320 | | | | |
| 5HT1a | 0.27 | 0.093 | 0.320 | | | | |
| 5HT1b | 0.18 | 0.115 | 0.346 | | | | |
| NET | 0.17 | 0.180 | 0.480 | | | | |
| NMDA | -0.06 | 0.590 | 1.000 | | | | |
| D1 | -0.07 | 0.623 | 1.000 | | | | |
| D2 | -0.12 | 0.697 | 1.000 | | | | |
| CB1 | -0.08 | 0.726 | 1.000 | | | | |
| 5HT4 | -0.13 | 0.737 | 1.000 | | | | |
| a4b2 | -0.16 | 0.755 | 1.000 | | | | |
| FDOPA | -0.19 | 0.782 | 1.000 | | | | |
| DAT | -0.19 | 0.788 | 1.000 | | | | |
| OXT | -0.19 | 0.791 | 1.000 | | | | |
| H3 | -0.20 | 0.908 | 1.000 | | | | |
| MU | -0.23 | 0.929 | 1.000 | | | | |
| 5HTT | -0.35 | 0.942 | 1.000 | | | | |
| VAChT | -0.29 | 0.956 | 1.000 | | | | |
| OXTR | -0.39 | 0.984 | 1.000 | | | | |
| CD38 | -0.57 | 1.000 | 1.000 | | | | |

**GABA-related gene co-expression clusters**

| Atlas | Whole-brain | | |
|---|---|---|---|
| | *Z (ρ)* | *p* | *q* |
| Cluster 3 | 0.45 | 0.005 | 0.014 |
| Cluster 2 | 0.43 | 0.007 | 0.014 |
| Cluster 4 | -0.19 | 0.932 | 0.952 |
| Cluster 1 | -0.23 | 0.952 | 0.952 |

#### Table S8: Results of neurotransmitter association analyses

Spatial correlations between interpersonal neural synchronization Z maps and neurotransmitter receptor distributions derived from in vivo nuclear imaging atlases (*144*, *145*) or post-mortem mRNA expression data (*19*, *20*). Correlation coefficients are r-to-Z transformed partial Spearman's ρ, *p* values are uncorrected and derived from null-corelations, *q* values are false discovery-corrected *p* values according to the Benjamini-Hochberg procedure. The two right main columns show sensitivity analyses of the significant positive associations (i) calculated only on cortical parcels and (ii) calculated after additional inclusion of functional baseline activation rate (BL) in partial correlations of INS and nuclear imaging maps. Rows are sorted by main *p* values.

**Neuronal cell types**

| Category | N (genes) | Average Z (ρ) | p (norm) | q (norm) | p (perm) | q (perm) |
|---|---|---|---|---|---|---|
| Adult-Ex3 | 9 | 0.36 | < 0.001 | 0.001 | < 0.001 | < 0.001 |
| Adult-In5 | 7 | 0.12 | 0.001 | 0.013 | 0.001 | 0.012 |
| Adult-In6 | 5 | 0.19 | 0.005 | 0.039 | 0.004 | 0.032 |
| Adult-Ex4 | 15 | 0.12 | 0.030 | 0.147 | 0.025 | 0.125 |
| Adult-OtherNeuron | 38 | 0.07 | 0.031 | 0.147 | 0.026 | 0.125 |
| Adult-Ex5 | 21 | 0.11 | 0.051 | 0.206 | 0.049 | 0.197 |
| Adult-Ex2 | 9 | 0.06 | 0.095 | 0.325 | 0.095 | 0.326 |
| Adult-Ex1 | 17 | -0.01 | 0.566 | 1.000 | 0.563 | 1.000 |
| Adult-Ex6 | 34 | -0.11 | 0.993 | 1.000 | 0.994 | 1.000 |
| Adult-Ex7 | 6 | -0.07 | 0.908 | 1.000 | 0.906 | 1.000 |
| Adult-Ex8 | 54 | -0.07 | 0.931 | 1.000 | 0.928 | 1.000 |
| Adult-In1 | 2 | -0.30 | 0.994 | 1.000 | 0.993 | 1.000 |
| Adult-In2 | 7 | -0.10 | 0.991 | 1.000 | 0.993 | 1.000 |
| Adult-In3 | 17 | -0.02 | 0.751 | 1.000 | 0.751 | 1.000 |
| Adult-In4 | 11 | -0.04 | 0.788 | 1.000 | 0.791 | 1.000 |
| Adult-In7 | 10 | -0.03 | 0.759 | 1.000 | 0.762 | 1.000 |
| Adult-In8 | 3 | 0.02 | 0.368 | 1.000 | 0.369 | 1.000 |
| Adult-Astro | 37 | -0.23 | 1.000 | 1.000 | 1.000 | 1.000 |
| Adult-Endo | 76 | -0.16 | 0.992 | 1.000 | 0.992 | 1.000 |
| Dev-quiescent | 15 | -0.04 | 0.785 | 1.000 | 0.780 | 1.000 |
| Dev-replicating | 25 | -0.05 | 0.971 | 1.000 | 0.973 | 1.000 |
| Adult-Micro | 20 | -0.25 | 0.998 | 1.000 | 0.999 | 1.000 |
| Adult-OPC | 39 | -0.19 | 0.999 | 1.000 | 0.999 | 1.000 |
| Adult-Oligo | 35 | -0.03 | 0.649 | 1.000 | 0.651 | 1.000 |

**Developmental regional enrichment**

| Category | N (genes) | Average Z (ρ) | p (norm) | q (norm) | p (perm) | q (perm) |
|---|---|---|---|---|---|---|
| child_CBC | 165 | 0.16 | < 0.001 | 0.001 | < 0.001 | < 0.001 |
| adolescent_CBC | 155 | 0.15 | < 0.001 | 0.001 | < 0.001 | < 0.001 |
| adult_A1C | 31 | 0.32 | < 0.001 | 0.001 | < 0.001 | < 0.001 |
| adult_V1C | 67 | 0.32 | < 0.001 | 0.001 | < 0.001 | < 0.001 |
| adult_CBC | 193 | 0.14 | < 0.001 | 0.001 | < 0.001 | < 0.001 |
| adolescent_STC | 5 | 0.50 | < 0.001 | < 0.001 | < 0.001 | 0.001 |
| adult_STC | 67 | 0.28 | < 0.001 | 0.001 | < 0.001 | 0.001 |
| adult_S1C | 45 | 0.31 | < 0.001 | 0.001 | < 0.001 | 0.001 |
| adult_IPC | 55 | 0.27 | < 0.001 | 0.001 | < 0.001 | 0.001 |
| infant_CBC | 121 | 0.10 | < 0.001 | 0.001 | < 0.001 | 0.001 |
| adult_M1C | 30 | 0.27 | < 0.001 | 0.002 | < 0.001 | 0.001 |
| adolescent_DFC | 27 | 0.19 | < 0.001 | 0.001 | < 0.001 | 0.002 |
| adolescent_VFC | 8 | 0.24 | < 0.001 | 0.001 | < 0.001 | 0.002 |
| adult_VFC | 43 | 0.26 | < 0.001 | 0.001 | < 0.001 | 0.002 |
| infant_V1C | 47 | 0.21 | < 0.001 | 0.001 | 0.001 | 0.003 |
| adolescent_OFC | 14 | 0.26 | < 0.001 | 0.001 | 0.001 | 0.003 |
| adolescent_IPC | 17 | 0.31 | < 0.001 | 0.001 | 0.001 | 0.004 |
| adult_OFC | 41 | 0.25 | < 0.001 | 0.001 | 0.001 | 0.004 |
| infant_ITC | 8 | 0.19 | 0.001 | 0.005 | 0.001 | 0.005 |
| adult_DFC | 57 | 0.21 | 0.001 | 0.005 | 0.002 | 0.007 |
| adolescent_V1C | 11 | 0.18 | 0.002 | 0.008 | 0.002 | 0.007 |
| infant_IPC | 16 | 0.21 | 0.002 | 0.008 | 0.003 | 0.010 |
| infant_MFC | 8 | 0.18 | 0.004 | 0.013 | 0.004 | 0.014 |
| adult_ITC | 37 | 0.17 | 0.004 | 0.014 | 0.006 | 0.019 |
| child_V1C | 31 | 0.12 | 0.006 | 0.019 | 0.006 | 0.019 |
| infant_S1C | 35 | 0.15 | 0.008 | 0.024 | 0.007 | 0.022 |
| prenatal_V1C | 171 | 0.07 | 0.009 | 0.025 | 0.008 | 0.024 |
| adolescent_S1C | 9 | 0.15 | 0.010 | 0.028 | 0.009 | 0.025 |
| adolescent_ITC | 15 | 0.13 | 0.010 | 0.028 | 0.010 | 0.026 |

**DisGeNET psychiatric disease markers**

| Category | N (genes) | Average Z (ρ) | p (norm) | q (norm) | p (perm) | q (perm) |
|---|---|---|---|---|---|---|
| Seasonal Affective Disorder | 14 | 0.200 | < 0.001 | 0.001 | < 0.001 | < 0.001 |

| | | | | | | |
|---|---|---|---|---|---|---|
| Psychosis, Brief Reactive | 11 | 0.238 | < 0.001 | 0.001 | < 0.001 | < 0.001 |
| Schizophreniform Disorders | 11 | 0.238 | < 0.001 | 0.001 | < 0.001 | < 0.001 |
| Language Development Disorders | 11 | 0.305 | < 0.001 | 0.001 | < 0.001 | 0.006 |
| Speech Delay | 11 | 0.305 | < 0.001 | 0.001 | < 0.001 | 0.006 |
| Semantic-Pragmatic Disorder | 11 | 0.305 | < 0.001 | 0.001 | < 0.001 | 0.006 |
| Neurodevelopmental Disorders | 89 | 0.151 | < 0.001 | 0.001 | < 0.001 | 0.006 |
| Schizoaffective Disorder | 25 | 0.136 | < 0.001 | 0.003 | < 0.001 | 0.006 |
| Abnormal behavior | 15 | 0.173 | < 0.001 | 0.003 | < 0.001 | 0.006 |
| Global developmental delay | 125 | 0.081 | < 0.001 | 0.006 | < 0.001 | 0.006 |
| Major Affective Disorder 2 | 13 | 0.156 | < 0.001 | 0.003 | 0.001 | 0.011 |
| Opioid-Related Disorders | 5 | 0.128 | 0.001 | 0.012 | 0.003 | 0.022 |
| Opioid abuse | 5 | 0.128 | 0.001 | 0.012 | 0.003 | 0.022 |
| Narcotic Abuse | 5 | 0.128 | 0.001 | 0.012 | 0.003 | 0.022 |
| Opiate Addiction | 5 | 0.128 | 0.001 | 0.012 | 0.003 | 0.022 |
| Narcotic Dependence | 5 | 0.128 | 0.001 | 0.012 | 0.003 | 0.022 |
| Opiate Abuse | 5 | 0.128 | 0.001 | 0.012 | 0.003 | 0.022 |
| Manic Disorder | 65 | 0.061 | 0.003 | 0.025 | 0.003 | 0.022 |
| Phencyclidine Abuse | 7 | 0.146 | 0.004 | 0.025 | 0.003 | 0.023 |
| Phencyclidine-Related Disorders | 7 | 0.146 | 0.004 | 0.025 | 0.003 | 0.023 |
| Depression, Postpartum | 6 | 0.189 | 0.004 | 0.025 | 0.004 | 0.027 |
| Intellectual Disability | 416 | 0.047 | 0.004 | 0.026 | 0.004 | 0.027 |
| Developmental Coordination Disorder | 7 | 0.117 | 0.005 | 0.034 | 0.005 | 0.031 |
| Motor Skills Disorders | 7 | 0.117 | 0.005 | 0.034 | 0.005 | 0.031 |
| Profound Mental Retardation | 126 | 0.041 | 0.008 | 0.043 | 0.006 | 0.036 |
| Mental Retardation, Psychosocial | 126 | 0.041 | 0.008 | 0.043 | 0.006 | 0.036 |
| Mental deficiency | 126 | 0.041 | 0.008 | 0.043 | 0.006 | 0.036 |
| Manic | 71 | 0.048 | 0.010 | 0.054 | 0.008 | 0.045 |

**Gene ontology biological processes**

| Category | N (genes) | Average Z (ρ) | p (norm) | q (norm) | p (perm) | q (perm) |
|---|---|---|---|---|---|---|
| neg. reg. of voltage-gated calcium channel activity | 5 | 0.23 | < 0.001 | 0.008 | < 0.001 | < 0.001 |
| pos. reg. of skeletal muscle cell differentiation | 5 | 0.42 | < 0.001 | 0.008 | < 0.001 | < 0.001 |
| reg. of water loss via skin | 9 | 0.21 | < 0.001 | 0.011 | < 0.001 | < 0.001 |
| pos. reg. of nonmotile primary cilium assembly | 6 | 0.24 | < 0.001 | 0.011 | < 0.001 | < 0.001 |
| deadenylation-dependent decapping of nuclear-transcribed mRNA | 8 | 0.28 | < 0.001 | 0.011 | < 0.001 | < 0.001 |
| protein K6-linked ubiquitination | 7 | 0.26 | < 0.001 | 0.011 | < 0.001 | < 0.001 |
| reg. of macromitophagy | 5 | 0.28 | < 0.001 | 0.011 | < 0.001 | < 0.001 |
| cerebellar Purkinje cell differentiation | 10 | 0.23 | < 0.001 | 0.011 | < 0.001 | < 0.001 |
| reg. of Golgi to plasma membrane protein transport | 7 | 0.37 | < 0.001 | 0.011 | < 0.001 | < 0.001 |
| reg. of mRNA splicing, via spliceosome | 65 | 0.14 | < 0.001 | 0.011 | < 0.001 | < 0.001 |
| phosphorelay signal transduction system | 7 | 0.26 | < 0.001 | 0.011 | < 0.001 | < 0.001 |
| cardiac muscle adaptation | 17 | 0.16 | < 0.001 | 0.011 | < 0.001 | < 0.001 |
| muscle hypertrophy in response to stress | 16 | 0.19 | < 0.001 | 0.011 | < 0.001 | < 0.001 |
| cardiac muscle hypertrophy in response to stress | 16 | 0.19 | < 0.001 | 0.011 | < 0.001 | < 0.001 |
| response to parathyroid hormone | 7 | 0.24 | < 0.001 | 0.011 | < 0.001 | < 0.001 |
| reg. of protein sumoylation | 21 | 0.14 | < 0.001 | 0.011 | < 0.001 | < 0.001 |
| cell differentiation in hindbrain | 16 | 0.17 | < 0.001 | 0.013 | < 0.001 | < 0.001 |
| reg. of Rac protein signal transduction | 12 | 0.21 | < 0.001 | 0.013 | 0.001 | 0.020 |
| histone H2A monoubiquitination | 12 | 0.21 | < 0.001 | 0.013 | < 0.001 | < 0.001 |
| N-terminal protein lipidation | 6 | 0.30 | < 0.001 | 0.013 | < 0.001 | < 0.001 |
| nephric duct development | 8 | 0.19 | < 0.001 | 0.013 | < 0.001 | 0.010 |
| neg. reg. of glucocorticoid receptor signaling pathway | 6 | 0.34 | < 0.001 | 0.013 | < 0.001 | 0.010 |
| histone H4-K5 acetylation | 16 | 0.17 | < 0.001 | 0.013 | < 0.001 | < 0.001 |
| histone H4-K8 acetylation | 16 | 0.17 | < 0.001 | 0.013 | < 0.001 | < 0.001 |
| cerebellar Purkinje cell layer formation | 11 | 0.19 | < 0.001 | 0.013 | < 0.001 | < 0.001 |
| reg. of skeletal muscle cell differentiation | 14 | 0.15 | < 0.001 | 0.013 | < 0.001 | < 0.001 |
| neg. reg. of mitophagy | 6 | 0.31 | < 0.001 | 0.013 | < 0.001 | < 0.001 |
| reg. of alternative mRNA splicing, via spliceosome | 35 | 0.13 | < 0.001 | 0.013 | < 0.001 | 0.015 |
| reg. of production of miRNAs involved in gene silencing by miRNA | 9 | 0.15 | < 0.001 | 0.013 | < 0.001 | < 0.001 |
| serine phosphorylation of STAT3 protein | 6 | 0.35 | < 0.001 | 0.013 | < 0.001 | < 0.001 |

| | | | | | | |
|---|---|---|---|---|---|---|
| pyrimidine nucleotide catabolic process | 11 | 0.15 | < 0.001 | 0.013 | < 0.001 | 0.010 |
| Golgi ribbon formation | 11 | 0.15 | < 0.001 | 0.013 | < 0.001 | 0.010 |
| startle response | 24 | 0.14 | < 0.001 | 0.013 | < 0.001 | < 0.001 |
| histone deubiquitination | 22 | 0.16 | < 0.001 | 0.013 | < 0.001 | 0.010 |
| neg. reg. of peptidyl-tyrosine phosphorylation | 32 | 0.13 | < 0.001 | 0.013 | < 0.001 | < 0.001 |
| establishment of meiotic spindle localization | 5 | 0.19 | < 0.001 | 0.013 | < 0.001 | < 0.001 |
| pos. reg. of circadian rhythm | 13 | 0.14 | < 0.001 | 0.013 | < 0.001 | < 0.001 |
| serine phosphorylation of STAT protein | 8 | 0.23 | < 0.001 | 0.013 | < 0.001 | < 0.001 |
| locomotion involved in locomotory behavior | 7 | 0.15 | < 0.001 | 0.013 | < 0.001 | < 0.001 |
| histone H4-K12 acetylation | 7 | 0.22 | < 0.001 | 0.013 | < 0.001 | 0.010 |
| macromitophagy | 17 | 0.26 | < 0.001 | 0.013 | < 0.001 | 0.010 |
| neg. reg. of histone H3-K9 methylation | 10 | 0.21 | < 0.001 | 0.013 | < 0.001 | < 0.001 |
| cerebellar Purkinje cell layer morphogenesis | 14 | 0.18 | < 0.001 | 0.013 | < 0.001 | < 0.001 |
| establishment of skin barrier | 7 | 0.18 | < 0.001 | 0.014 | < 0.001 | 0.010 |
| mRNA 3'-splice site recognition | 6 | 0.26 | < 0.001 | 0.014 | < 0.001 | 0.015 |
| reg. of circadian rhythm | 82 | 0.10 | < 0.001 | 0.014 | < 0.001 | 0.015 |
| response to muscle inactivity involved in reg. of muscle adaptation | 5 | 0.23 | < 0.001 | 0.014 | < 0.001 | < 0.001 |
| response to denervation involved in reg. of muscle adaptation | 5 | 0.23 | < 0.001 | 0.014 | < 0.001 | < 0.001 |
| reg. of type B pancreatic cell development | 7 | 0.39 | < 0.001 | 0.014 | 0.001 | 0.020 |
| protein desumoylation | 8 | 0.20 | < 0.001 | 0.014 | < 0.001 | < 0.001 |

##### Table S9: Results of gene category enrichment analyses

Gene category enrichment analyses were calculated using ABAnnotate (*32*) on different databases; neuronal cell type markers (*146–148*), regionwise developmental mRNA expression data (*149*), psychiatric disease markers (*150*) and Gene Ontology biological process annotations (*151*, *152*). The method was developed by Fulcher et al. (*153*). Spatial Spearman correlations were calculated between the interpersonal neural synchrony (INS) Z map and the gene expression profiles drawn from the Allen Brain Atlas (*19*) of each gene annotated to a category. Correlation coefficients were averaged for each category. This process was repeated for 5,000 null maps derived from the INS map and significance was evaluated by comparison of the “real” category-wise average correlation with the 5,000 null correlations (*p(perm)*). To approximate p-values below 1/5000, a Gassian distribution of categorywise average correlations was fitted to the distribution of permutation-derived scores (*p(norm)*). Resulting *p* values were false discovery rate-corrected according to the Benjamini-Hochberg procedure (*q*). Rows are sorted according to the parametric uncorrected *p* values. Shown are all results for cell types, only significant for regional developmental enrichment, and the top 50 Gene Ontology categories. Full results are provided in the GitHub repository accompanying this publication.

Abbreviations: perm = permutation, norm = Gaussian normal distribution, CBC = cerebellar cortex, A1C = primary auditory cortex, V1C = primary visual cortex, STC = posterior (caudal) superior temporal cortex, S1C = primary somatosensory cortex, IPC = posteroventral (inferior) parietal cortex, M1C = primary motor cortex, DFC = dorsolateral prefrontal cortex, VFC = ventrolateral prefrontal cortex, OFC = orbital frontal cortex, ITC = inferolateral temporal cortex, MFC = anterior (rostral) cingulate (medial prefrontal) cortex, reg = regulation, pos = positive, neg = negative.

| Cluster ID | N (terms) | *Representative GO term* | | | |
|---|---|---|---|---|---|
| | | GO term ID | GO term description | p (norm) | Average Z (ρ) |
| 1 | 16 | GO:1901386 | negative regulation of voltage-gated potassium channel activity | 2.120E-06 | 0.23 |
| 2 | 49 | GO:2001016 | positive regulation of skeletal muscle cell differentiation | 2.370E-06 | 0.42 |
| 3 | 36 | GO:0033561 | regulation of water loss via skin | 8.690E-06 | 0.21 |
| 4 | 80 | GO:0000290 | deadenylation-dependent decapping of nuclear-transcribed mRNA | 1.180E-05 | 0.28 |
| 5 | 13 | GO:0085020 | protein K6-linked ubiquitination | 1.210E-05 | 0.26 |
| 6 | 18 | GO:1901524 | regulation of mitophagy | 1.280E-05 | 0.28 |
| 7 | 26 | GO:0021702 | cerebellar Purkinje cell differentiation | 1.330E-05 | 0.23 |
| 8 | 19 | GO:0071107 | response to parathyroid hormone | 2.450E-05 | 0.24 |
| 9 | 30 | GO:0033233 | regulation of protein sumoylation | 2.470E-05 | 0.14 |
| 10 | 21 | GO:0006498 | N-terminal protein lipidation | 4.560E-05 | 0.30 |
| 11 | 9 | GO:2000323 | negative regulation of glucocorticoid receptor signaling pathway | 4.700E-05 | 0.34 |
| 12 | 15 | GO:0051295 | establishment of meiotic spindle localization | 7.030E-05 | 0.19 |
| 13 | 21 | GO:0042501 | serine phosphorylation of STAT protein | 7.190E-05 | 0.23 |
| 14 | 5 | GO:0031987 | locomotion involved in locomotory behavior | 7.250E-05 | 0.15 |
| 15 | 31 | GO:0000389 | mRNA 3'-splice site recognition | 9.360E-05 | 0.26 |
| 16 | 13 | GO:0014894 | response to denervation involved in regulation of muscle adaptation | 1.003E-04 | 0.23 |
| 17 | 16 | GO:0030213 | hyaluronan biosynthetic process | 1.069E-04 | 0.19 |
| 18 | 6 | GO:0032042 | mitochondrial DNA metabolic process | 1.802E-04 | 0.16 |
| 19 | 2 | GO:0086027 | AV node cell to bundle of His cell signaling | 1.835E-04 | 0.30 |
| 20 | 6 | GO:0000301 | retrograde transport, vesicle recycling within Golgi | 2.339E-04 | 0.13 |
| 21 | 19 | GO:0015851 | nucleobase transport | 2.574E-04 | 0.34 |
| 22 | 2 | GO:0042118 | endothelial cell activation | 3.987E-04 | 0.17 |
| 23 | 1 | GO:0010659 | cardiac muscle cell apoptotic process | 6.881E-04 | 0.08 |
| 24 | 1 | GO:0008228 | opsonization | 1.870E-03 | 0.21 |
| 25 | 2 | GO:0039703 | RNA replication | 1.968E-03 | 0.08 |
| 26 | 1 | GO:0039694 | viral RNA genome replication | 1.968E-03 | 0.08 |
| 27 | 2 | GO:0086003 | cardiac muscle cell contraction | 3.916E-03 | 0.07 |

**Table S10: Results of GeneOntology category clustering**

The data were generated using GO-Figure! (*154*). GO categories significantly associated to the interpersonal neural synchrony Z map (see Table S9) were clustered based on semantic similarity (similarity threshold ≥ .2) and for each cluster a representative term was selected in a data-driven fashion. *N (terms)* shows the number of GO categories annoatated to each cluster. The list is ordered according to the p-values of the representative terms and shows only GO terms that could be clustered. See Table S9 for further descriptions.
Abbreviations: GO = GeneOntology.

### 5 Supplementary Figures

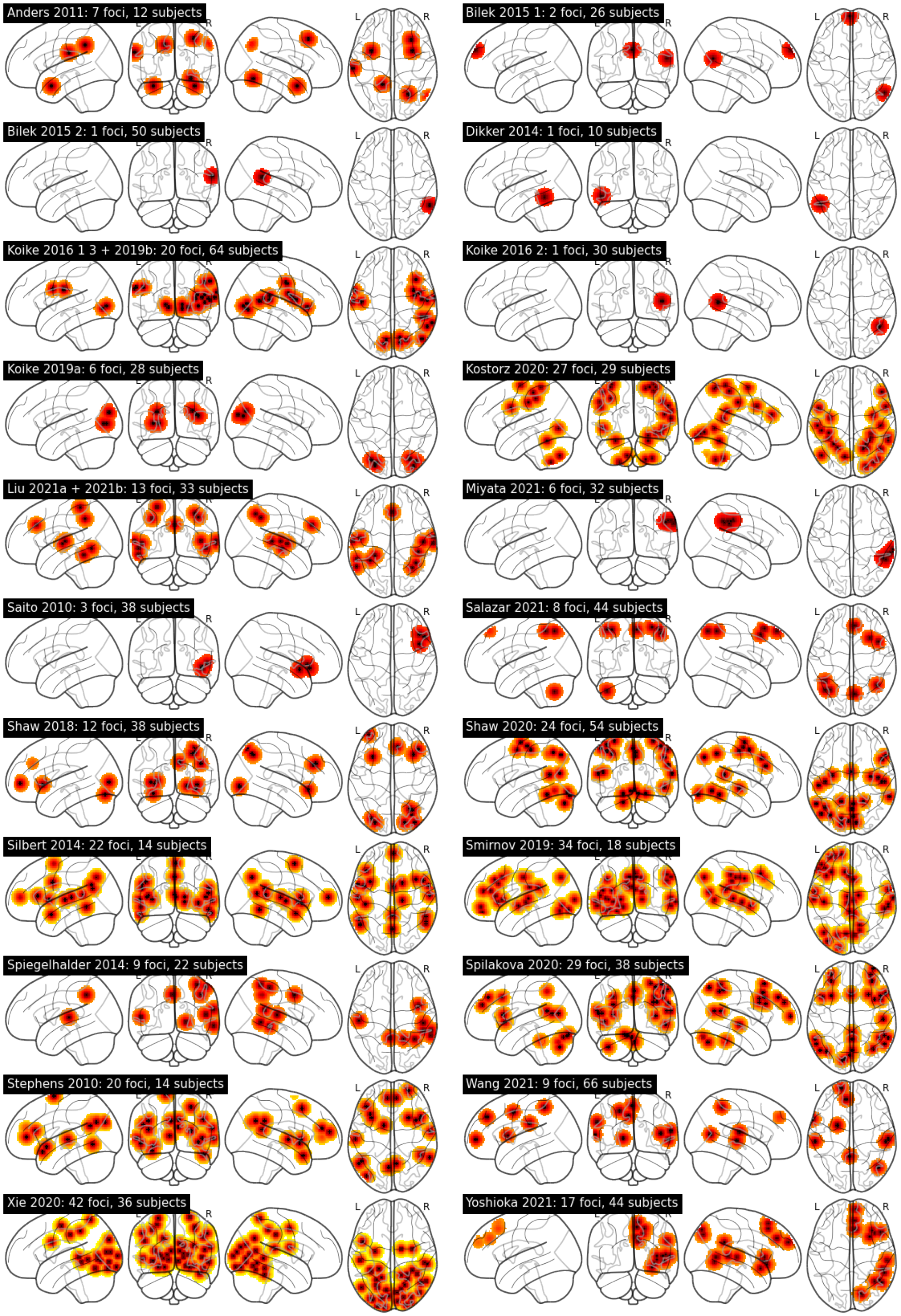

**Figure S1: INS foci reported in each fMRI experiment**

Foci are plotted individually for each experiment and smoothed using the activation likelihood estimation kernel depending on the reported sample size.

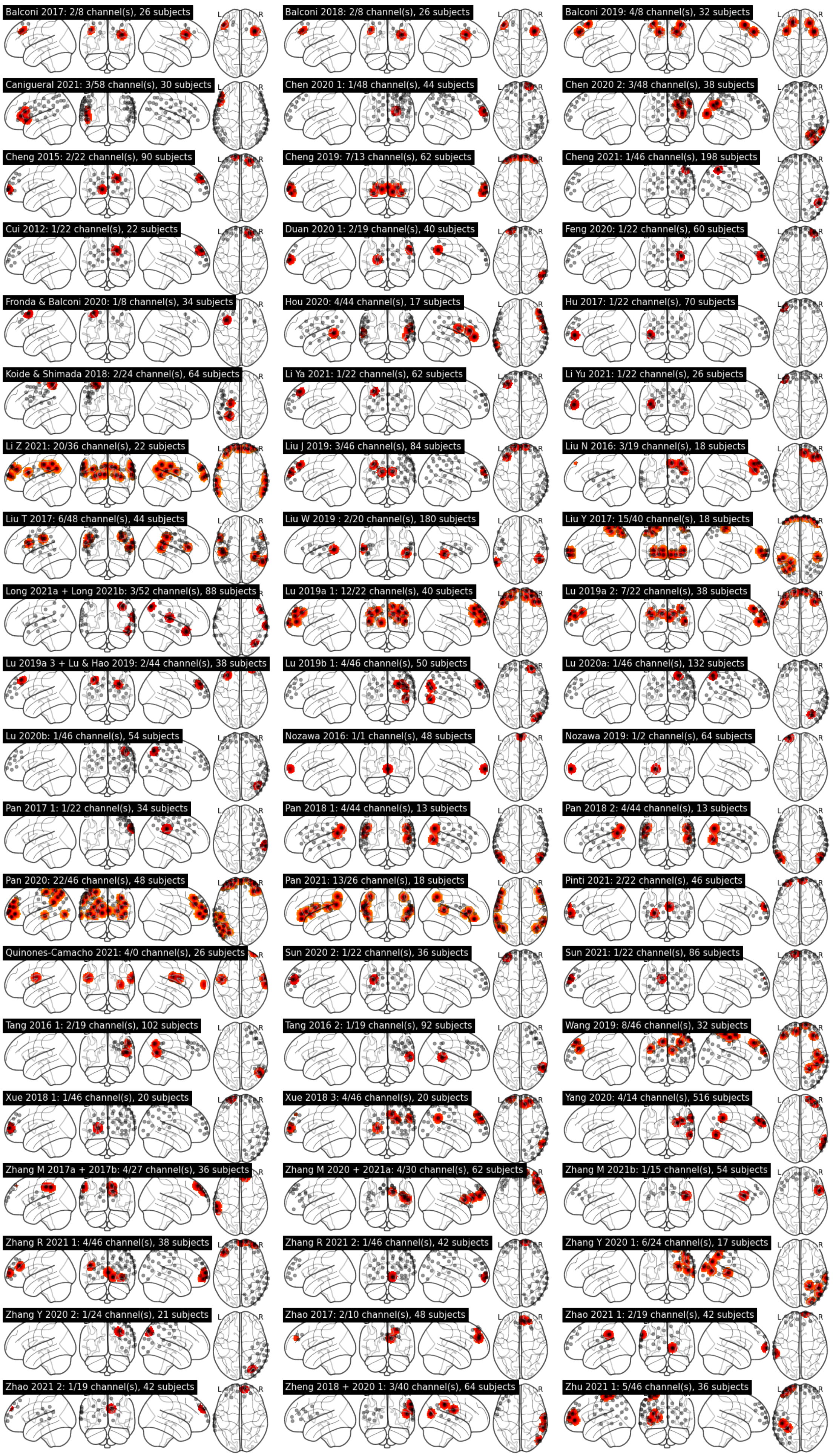
Balconi 2017: 2/8 channel(s), 26 subjects
Balconi 2018: 2/8 channel(s), 26 subjects
Balconi 2019: 4/8 channel(s), 32 subjects
Canigueral 2021: 3/58 channel(s), 30 subjects
Chen 2020 1: 1/48 channel(s), 44 subjects
Chen 2020 2: 3/48 channel(s), 38 subjects
Cheng 2015: 2/22 channel(s), 90 subjects
Cheng 2019: 7/13 channel(s), 62 subjects
Cheng 2021: 1/46 channel(s), 198 subjects
Cui 2012: 1/22 channel(s), 22 subjects
Duan 2020 1: 2/19 channel(s), 40 subjects
Feng 2020: 1/22 channel(s), 60 subjects
Fronda & Balconi 2020: 1/8 channel(s), 34 subjects
Hou 2020: 4/44 channel(s), 17 subjects
Hu 2017: 1/22 channel(s), 70 subjects
Koide & Shimada 2018: 2/24 channel(s), 64 subjects
Li Ya 2021: 1/22 channel(s), 62 subjects
Li Yu 2021: 1/22 channel(s), 26 subjects
Li Z 2021: 20/36 channel(s), 22 subjects
Liu J 2019: 3/46 channel(s), 84 subjects
Liu N 2016: 3/19 channel(s), 18 subjects
Liu T 2017: 6/48 channel(s), 44 subjects
Liu W 2019 : 2/20 channel(s), 180 subjects
Liu Y 2017: 15/40 channel(s), 18 subjects
Long 2021a + Long 2021b: 3/52 channel(s), 88 subjects
Lu 2019a 1: 12/22 channel(s), 40 subjects
Lu 2019a 2: 7/22 channel(s), 38 subjects
Lu 2019a 3 + Lu & Hao 2019: 2/44 channel(s), 38 subjects
Lu 2019b 1: 4/46 channel(s), 50 subjects
Lu 2020a: 1/46 channel(s), 132 subjects
Lu 2020b: 1/46 channel(s), 54 subjects
Nozawa 2016: 1/1 channel(s), 48 subjects
Nozawa 2019: 1/2 channel(s), 64 subjects
Pan 2017 1: 1/22 channel(s), 34 subjects
Pan 2018 1: 4/44 channel(s), 13 subjects
Pan 2018 2: 4/44 channel(s), 13 subjects
Pan 2020: 22/46 channel(s), 48 subjects
Pan 2021: 13/26 channel(s), 18 subjects
Pinti 2021: 2/22 channel(s), 46 subjects
Quinones-Camacho 2021: 4/0 channel(s), 26 subjects
Sun 2020 2: 1/22 channel(s), 36 subjects
Sun 2021: 1/22 channel(s), 86 subjects
Tang 2016 1: 2/19 channel(s), 102 subjects
Tang 2016 2: 1/19 channel(s), 92 subjects
Wang 2019: 8/46 channel(s), 32 subjects
Xue 2018 1: 1/46 channel(s), 20 subjects
Xue 2018 3: 4/46 channel(s), 20 subjects
Yang 2020: 4/14 channel(s), 516 subjects
Zhang M 2017a + 2017b: 4/27 channel(s), 36 subjects
Zhang M 2020 + 2021a: 4/30 channel(s), 62 subjects
Zhang M 2021b: 1/15 channel(s), 54 subjects
Zhang R 2021 1: 4/46 channel(s), 38 subjects
Zhang R 2021 2: 1/46 channel(s), 42 subjects
Zhang Y 2020 1: 6/24 channel(s), 17 subjects
Zhang Y 2020 2: 1/24 channel(s), 21 subjects
Zhao 2017: 2/10 channel(s), 48 subjects
Zhao 2021 1: 2/19 channel(s), 42 subjects
Zhao 2021 2: 1/19 channel(s), 42 subjects
Zheng 2018 + 2020 1: 3/40 channel(s), 64 subjects
Zhu 2021 1: 5/46 channel(s), 36 subjects

**Figure S2: INS foci reported in each fNIRS experiment**

Interpersonal neural synchrony (INS) channels are plotted individually for each experiment and, for vizualization purposes, smoothed using the activation likelihood estimation kernel set to a fixed “sample size” of n = 10. Gray markers show all channels reported by, or reconstructed for, each study. Only those experiments reporting at least one INS channel are shown.

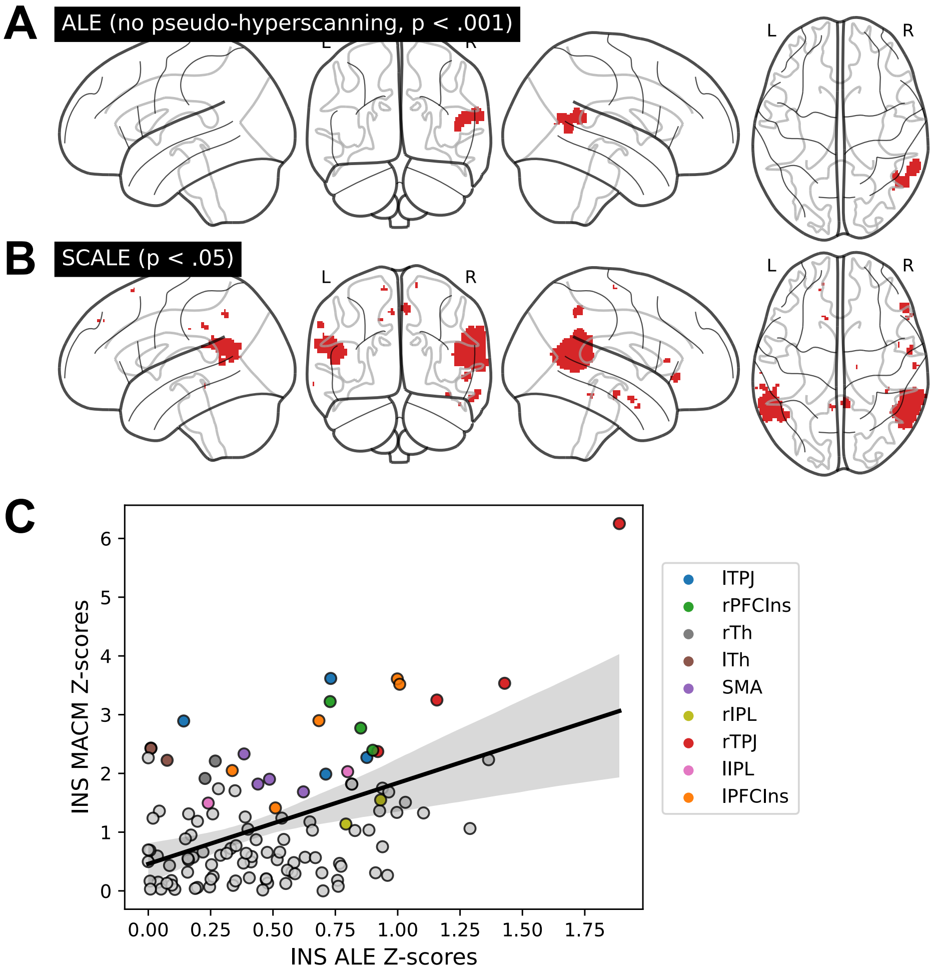

**Figure S3: ALE and MACM sensitivity analyses**

**A**: ALE without pseudo-hyperscanning studies. **B**: MACM corrected for baseline activation (SCALE). The method as implemented in NiMARE (*158*) does not support multiple comparison correction and was thresholded at voxel-level $p < .05$, uncorrected, for demonstration purposes. **C**: Spatial correlation between whole-brain distributions of INS and the rTPJ-associated MACM map to demonstrate comparable distributions patterns going beyond rTPJ activation.
Abbreviations: ALE = activation likelihood estimation, SCALE = specific coactivation likelihood estimation, INS = interpersonal neural synchronization, MACM = meta-analytic coactivation modeling, r/lTPJ = right/left temporoparietal junction, rSTG = right superior temporal gyrus, rIns = right insula, r/lPFCIns = right/left prefrontal cortex-insula, SMA = supplementary motor area, r/lTh = left thalamus, r/lIPL = left inferior parietal lobule, lPrec = left precuneus.

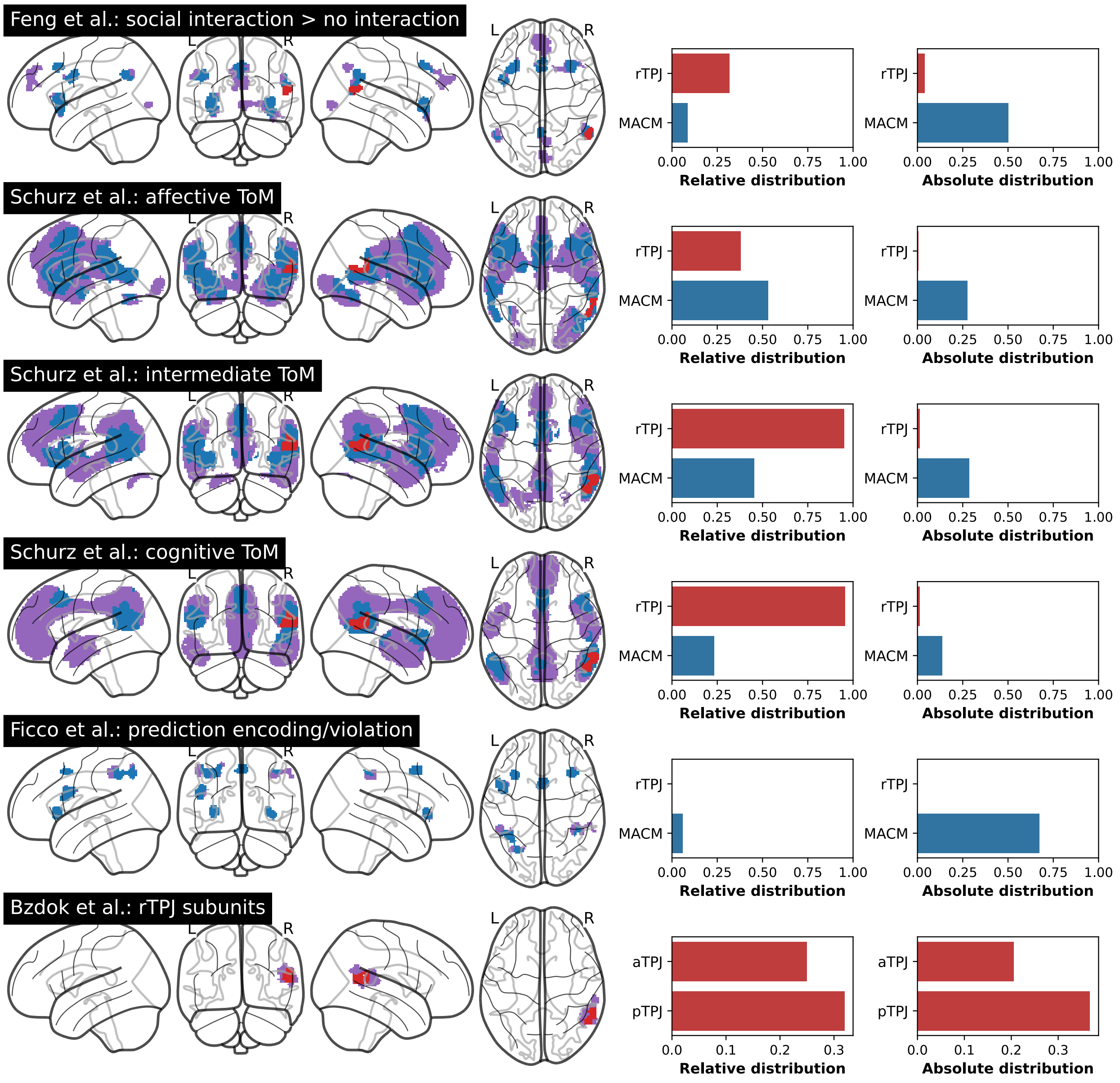

**Figure S4: Spatial overlap with prior meta-analytically derived networks and rTPJ subunits**

Sources: Feng et al., 2021 (*159*); Schurz et al., 2021 (*160*); Ficco et al., 2021 (*161*); Bzdok et al., 2013 (*162*). Original networks are plotted in purple, overlayed by conjunctions with the rTPJ-MACM network (blue) and the rTPJ cluster (red). For the upper five rows, the right side shows relative and absolute distributions of rTPJ cluster and MACM network within the corresponding meta-analytically derived network. The lowest row shows distributions of the INS-rTPJ cluster within two subunits of the rTPJ. Relatived distribution: proportion of "INS-voxels" within a given network vs. all "INS-voxels". Absolute distribution: proportion of "INS-voxels" within a given network vs. all voxels within the network.
Abbreviations: rTPJ = right temporoparietal junction, MACM = meta-analytic connectivity modeling, ToM = theory of mind, INS = interpersonal neural synchronization.

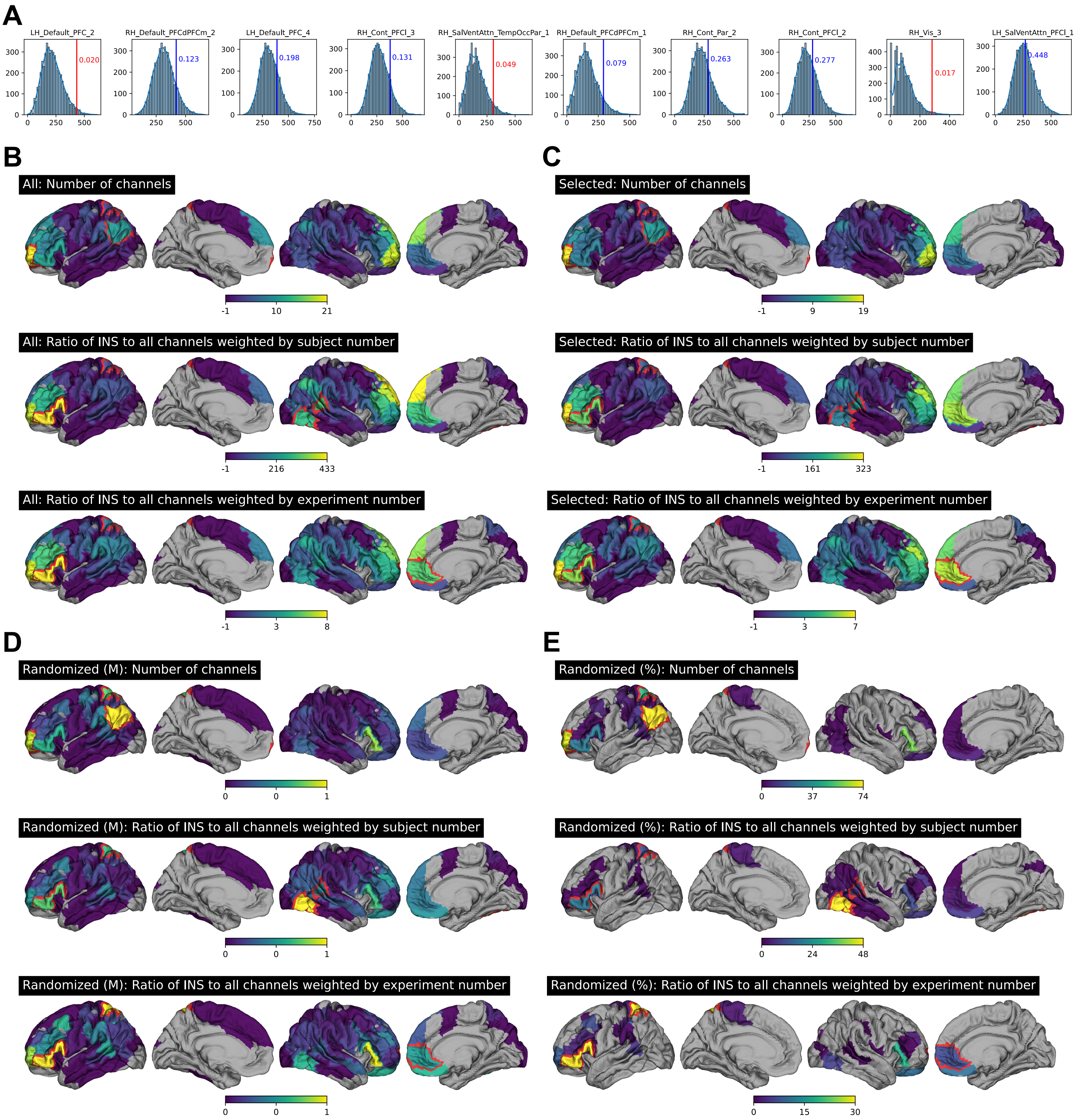

**Figure S5: Complete meta-analytic results of INS fNIRS experiments**

**A**: Histograms for the top ten regions resulting from the permutations regarding the meta-analytic fNIRS INS analysis that included all experiments and focused on the parcelwise ratio of INS vs. all channels weighted by the number of subjects. Vertical lines and numbers show the "real" result and the uncorrected *p* value estimated from each null distribution. **B**: Full fNIRS meta-analysis result for all included experiments. The middle row is shown in main Figure 2B. **C**: Full fNIRS meta-analysis result restricted to experiments explicitly assessing INS compared to rest, control, or randomization. **D**: Median (M) $\log_{10}$-transformed *p* values resulting from fNIRS meta-analysis after repeated randomization of fNIRS coordinates (radius = 1cm, 1,000 iterations). **E**: Percentage of sub-threshold *p* values after coordinate randomization.

Abbreviations: INS = interpersonal neural synchrony, fNIRS = functional near-infrared spectroscopy.

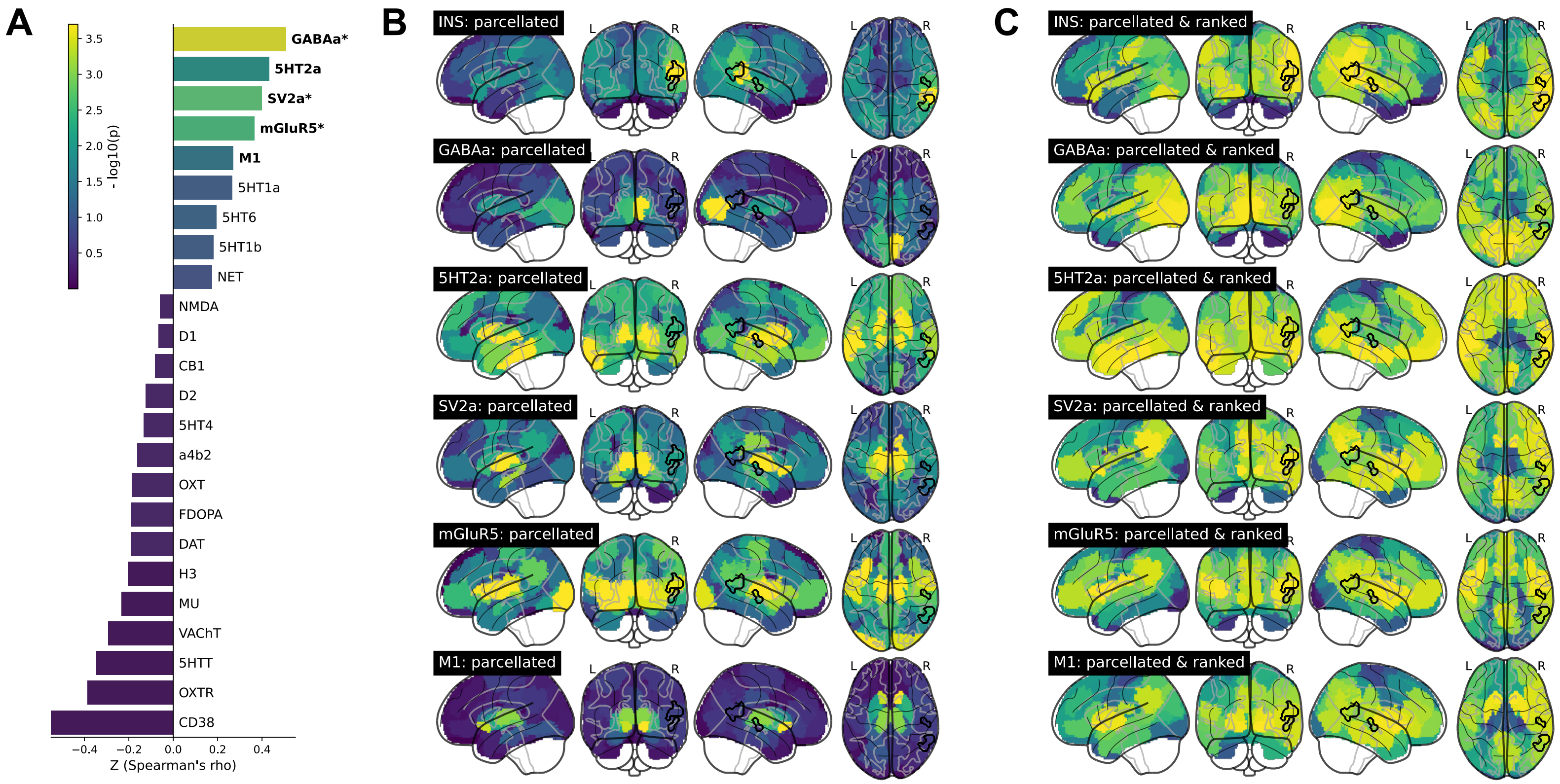

**Figure S6: Spatial associations of INS to neurotransmitter distributions**

**A**: Spatial correlations between whole-brain INS distribution and in vivo neurotransmitter receptor/ postmortem gene expression distributions. Color-coded $p$ values are uncorrected, correlations significant at uncorrected $p < .05$ derived from nonparametric permutation are printed in bold, $p$ values surviving false discovery rate correction ($q < .05$) are marked by asterisks. Nuclear imaging-derived maps were correlated with the INS map using partial spearman correlations controlling for local gray matter volume to account for partial volume effects. Correlation coefficients were compared to those calculated from 5,000 spatial autocorrelation-corrected null maps derived from each indicidual transmitter map using JuSpyce. **B**: Parcellated and Z-transformed transmitter PET significantly associated to INS. **C**: To better demonstrate the spatial alignment between INS and PET maps, parcellated values were ranked and the ranks were reassigned to the brain volumes (yellow = highest rank). Black outline shows significant clusters resulting from the main activation likelihood estimation analysis.
See Table S3 for full neurotransmitter receptor names, nuclear imaging tracers, and template sources, and Table S8 for full JuSpyce results.
Abbreviations: INS = interpersonal neural synchrony, PET = positron emission tomography.

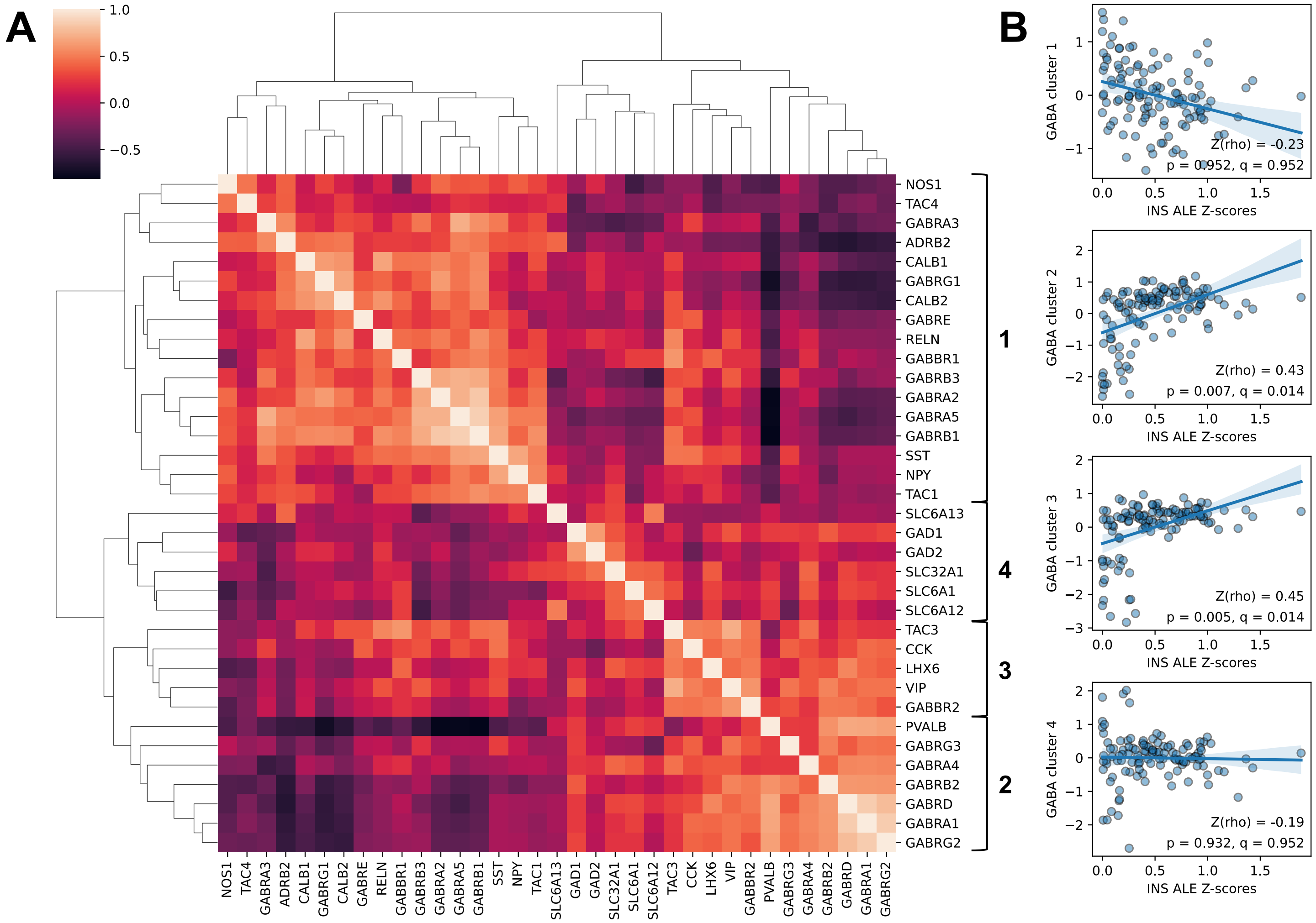

**Figure S7: Validation of INS-GABA$_A$ associations using mRNA expression data**

**A**: Clustered correlation heatmap of expression profiles of GABA-related genes. Colors represent Spearman correlations. Coexpression clusters were estimated using a hierarchical clustering algorithm based on euclidean pairwise distances. **B**: Correlation between the INS distribution and gene clusterwise averaged expression values, the correlation with clusters 2 and 3 are significant ($q < .05$).
Abbreviations: INS = interpersonal neural synchronization, ALE = activation likelihood estimation.

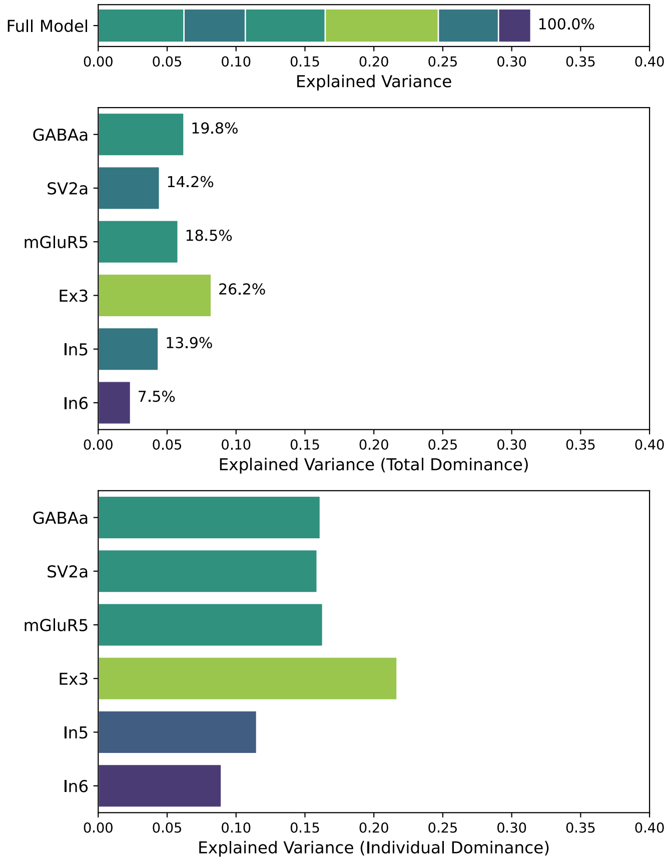

**Figure S8: Variance of the INS distribution explained by significantly associated nuclear imaging and neuronal cell type data**

Results from a dominance analysis using significantly positively associated nuclear imaging and neuronal cell type atlases as predictors and the INS Z map as target. As in the main univariate correlation analyses, local gray matter volume was regressed out of both predictors and target before performing the dominance analysis. Upper: $R^2$ of the complete multilinear regression model ($R^2 = .31$). Middle: "Total dominance" which can be interpreted as the relative contribution of a predictor to the explained variance of the complete model in interaction with the remaining predictors and therefore can also be expressed in percent of total $R^2$. Lower: "Individual dominance" which can be interpreted as each individual predictor's explained variance independent of all other predictors.
Abbreviations: INS = interpersonal neural synchronization.

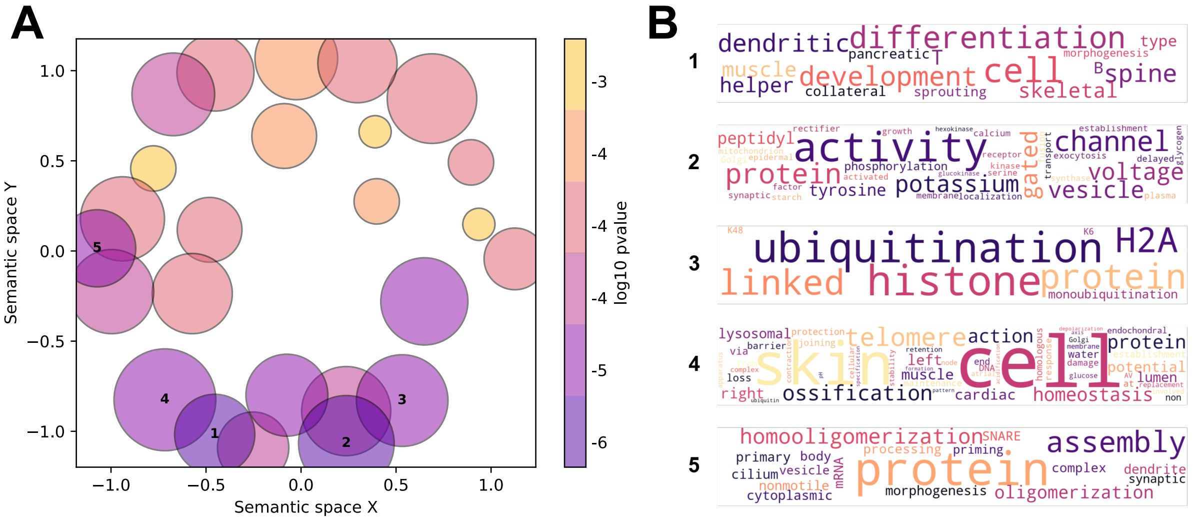

**Figure S9: Clustering of GeneOntology biological process categories spatially associated with INS**

**A**: two-dimensional reduction of the pairwise semantic similarity matrix of 474 Gene Ontology categories spatially associated to INS. Each point represents one cluster of n biologial process categories formed by a similarity threshold of .2 with the point size depicting n. Point color represents $log(p)$ associated with a representive GO term identified for each cluster. **B**: Word clouds build from combined category descriptions of the five semantic clusters most strongly associated to INS (marked by numbers 1-5 in both panels). Word size is based on word frequency within each semantic cluster.
